# Reprogramming tumour-associated macrophages from immune suppressive to inflammatory state by Checkpoint kinase 1 inhibitor combination treatment

**DOI:** 10.64898/2026.05.13.724422

**Authors:** Zhen Zeng, Anastasia Gandini, Ritu Bhatt, Martina Proctor, Jintao Guo, Susan Millard, Sherry Wu, Riccardo Dolcetti, James W. Wells, Jazmina Gonzalez Cruz, Katharine Irvine, Brian Gabrielli

**Author notes:** Corresponding authors: Prof Brian Gabrielli, Dr Zhen Zeng.

## Abstract

**Background:** Tumour-associated macrophages (TAMs) play critical roles within the tumour microenvironment regulating immune evasion and therapeutic response. Previously, we have shown that the combination of Checkpoint kinase 1 inhibitor (CHK1i) with a subclinical dose of hydroxyurea (LDHU) reprograms the tumour immune microenvironment to a pro-inflammatory status.

**Methods:** We investigated a tumour-restricted Fcgr4 (Cd16.2) expressing macrophage population in multiple murine tumour models and the impact of CHK1i+LDHU on this population, using conventional and imaging flow cytometry as well as single-cell sequencing.

**Results:** Transcriptional profiling using CITE-seq and single-cell RNA sequencing reveals that Fcgr4⁺ TAMs closely resemble Fcgr4⁻ TAMs but display modest enrichment of interferon-associated and inflammatory gene programs, consistent with a functionally biased state rather than a distinct lineage. Importantly, we show that a highly tumour selective CHK1i+LDHU therapy shifts TAMs toward a more inflammatory phenotype while preserving dominant immunosuppressive features. Depletion of CSF1R⁺ macrophages enhanced CD8⁺ T cell activation without influencing tumour growth but significantly augmented therapeutic efficacy of CHK1i+LDHU.

**Conclusion:** Together, these findings define a novel TAM population and establish how targeted therapy reshapes, but does not fully overcome, TAM-mediated immune regulation.

## Introduction

Tumour-associated macrophages (TAMs) are a dominant immune population within the tumour microenvironment where they play critical roles in regulating tumour growth, immune evasion, angiogenesis, metastasis, and therapeutic response [1]. Across multiple solid malignancies, including breast, lung, pancreatic, ovarian, and colorectal cancers, increased TAM infiltration correlates with poorer clinical outcomes [2, 3]. These observations underscore the central contribution of macrophages to tumour progression and highlight the need to better define their functional states within tumours, and importantly, changes in response to anti-cancer treatments.

Macrophages are highly plastic innate immune cells derived from circulating monocytes or tissue-resident precursors and are capable of adopting diverse functional states in response to environmental cues [4–7]. Although macrophage activation was historically simplified into M1/M2 polarization model, it is now clear that TAMs exist along a dynamic continuum shaped by tumour-derived signals, hypoxia, metabolic stress, and interactions with stromal and immune cells [1, 4, 8].

Mechanistically, TAMs suppress anti-tumour immunity through secretion of immunoregulatory cytokines such as interleukin-10 (IL-10) and transforming growth factor-β (TGF-β), expression of immune checkpoint ligands such as Tim3, and metabolic regulation of the tumour microenvironment [9–12]. Elevated TAM expression of Arginase (*Arg1*) and heme oxygenase 1 (HO-1; *Hmox1*), among other mediators, contributes to impaired cytotoxic T cell and natural killer (NK) cell function by altering the tumour microenvironment [13]. In parallel, TAMs promote tumour angiogenesis and metastasis through production of vascular endothelial growth factor (VEGF), matrix metalloproteinases (MMPs), and factors that facilitate extracellular matrix remodelling and pre-metastatic niche formation [1]. Together, these multifaceted pro-tumoral functions underscore the central contribution of TAMs to tumour immune suppression and disease progression. TAMs have also been implicated in resistance to chemotherapy, radiotherapy, and immune checkpoint blockade, positioning them as attractive targets for combination therapies [14, 15].

We have previously reported that the combination of Checkpoint kinase 1 inhibitor (CHK1i) with a subclinical dose of the ribonucleotide inhibitor hydroxyurea (low dose HU, LDHU) not only selectively kills a range of tumour types with little normal tissue toxicity but appears to reprogram TAMs from a pro-tumour to pro-inflammatory phenotype [16–18]. Here, we have investigated this change in macrophage phenotype in more detail using single cell RNA sequencing to assess the contribution of these reprogrammed TAMs to tumour growth and treatment response.

## Materials and Methods

### Cell lines

Immune-edited mouse melanoma cell lines YUMMUV1.7ie and YUMMUV3.3ie were generated as previously described [17]. Additional murine tumour cell lines used in this study included ID8p53^-/-^, B16F10. Cells were cultured in RPMI 1640 (Gibco) or high-glucose DMEM (Gibco) supplemented with 10% heat-inactivated fetal bovine serum (FBS; Bovogen), 1 mM sodium pyruvate (Gibco), GlutaMAX (Gibco), 20 mM HEPES (Sigma-Aldrich), and 1× antibiotic-antimycotic (Gibco). Cultures were maintained at 37°C in a humidified incubator (Binder) with 5% CO₂ and 2% O₂.

To generate CSF1 knockdown YUMMUV1.7ie clones, cells were transduced with lentiviral particles (Cat# 169120940296; Applied Biological Materials Inc., Richmond, BC, Canada) encoding a CSF1-specific short hairpin RNA (shRNA), GFP reporter, and puromycin resistance gene. Following puromycin selection, resistant colonies were expanded and assessed for CSF1 knockdown by fluorescence microscopy (GFP expression). GFP-positive cells were further purified by fluorescence-activated cell sorting.

### CSF1 quantification by LEGENDplex

Cytokine secretion was measured from untreated cell lines. Briefly, 3,000 cells were seeded in 96-well plates in the appropriate culture medium and incubated for 48h as previously described. After incubation, culture supernatants were collected and stored at −20 °C until analysis. M-CSF levels (740687) in the supernatants were quantified using a LEGENDplex bead-based immunoassay kit from BioLegend, following the manufacturer’s instructions for the LEGENDplex Multi-Analyte Flow Assay Kit. Samples were acquired on a BD LSRFortessa X-20 flow cytometer, and cytokine concentrations were calculated using LEGENDplex Qognit analysis software.

### RT-qPCR

Total RNA was extracted from cell lines using TRIzol (Invitrogen), and cDNA was generated from 2 µg RNA using the High-Capacity cDNA Reverse Transcription Kit (Applied Biosystems). qPCR was performed using PowerUp SYBR Green Master Mix (Applied Biosystems) on a ViiA 7 or QuantStudio 7 Flex system with KiCqStart primers (Merck). Relative gene expression was calculated using the 2^–ΔCt method with Actin (mouse) as reference gene.

### Mouse tumour assays

Experiments were performed with approval from The University of Queensland Animal Ethics Committee (2021/AE000249, 2022/AE000025 and 2022/AE000652). Tumour establishment: Syngeneic mouse YUMMUV1.7ie, YUMMUV3.3ie and ID8-p53^−/−^ tumours were established in C57BL/6J mice as described previously [17, 18]. B16F10 cells (2.5 × 10^4^) in 100 μl of Hank’s balanced salt solution were subcutaneously (s.c.) injected into C57BL6/J mice (RRID:IMSR JAX:000664). Following tumour establishment, mice were treated with CHK1i (SRA737) + LDHU as described previously [17]. Briefly, mice were treated every other day for 3 days per week for 3 weeks with vehicle (10% DMSO, 5% Tween80, 5% PEG400 in clinical-grade saline) or 50 mg/kg CHK1i (SRA737, Sierra Oncology, San Mateo, CA, USA) combined with 100 mg/kg HU in the vehicle by oral gavage, then 4 h later by i.p. injection of 50 mg/kg HU (in saline). To examine the influence of macrophages on treatment responses, anti-Csf1r (clone AFS98, Cat# BE0213, BioXcell) or appropriate isotype control injections were initiated 3 days prior to CHK1i+LDHU treatments and then repeated twice a week. At the experimental endpoint, tumours were collected, and other organ sites were dissected in a double-blinded manner.

### Immune Profiling

Tumours were harvested, mechanically minced, and enzymatically digested with DNase I and collagenase IV for 30 minutes at 37°C. The resulting cell suspensions were passed through a 40 µm cell strainer to generate single-cell suspensions. For splenocyte isolation, harvested spleens were gently dissociated through a 70 µm cell strainer using a syringe plunger. Red blood cells were lysed, and cells were washed and resuspended in staining buffer. Processed cells were incubated with Fc block TruStain FcX™ (anti-mouse CD16/32; Biolegend 101320) prior to staining with Live/Dead Aqua (Thermo Fisher Scientific, Waltham, MA, USA) and a panel of fluorochrome-conjugated antibodies (1:100 dilution) for immune cell profiling (Supplementary Table S1). Intracellular FoxP3 staining was performed using the FoxP3 Staining Kit (eBioscience) according to the manufacturer’s instructions. Absolute cell counts were determined using Flow-Count Fluorospheres (Beckman Coulter, Miami, FL, USA). Flow cytometric analyses were primarily performed using BD LSRFortessa instruments. For extended panels, stained cells were analysed using a Cytek Aurora spectral flow cytometer (Cytek Biosciences) with SpectroFlo software. Spectral unmixing was performed as described in the Supplementary Materials. Cell sorting was performed using BD FACSAria Fusion sorters or a MoFlo Astrios cell sorter.

### Imaging flow cytometry

Sorted live GFP^+^ cells were stained with antibody cocktails (as outlined in the gating strategy and detailed in Supplementary? Table xx) in the presence of CD16/CD32 blocking reagent (TruStain FcX™ anti-mouse CD16/32 antibody; BioLegend) for 30 minutes on ice. Cells were then washed with 2% FBS/PBS and resuspended at 2 x 10^7^ cells/mL for acquisition. Samples were analysed using a three-laser (405 nm, 488 nm, 642 nm) Amnis ImageStreamX Mk II imaging flow cytometer (Luminex Corporation, Austin, TX, USA). Data were analysed using Amnis IDEAS Software (version 6.2).

### CITEseq + scRNAseq

#### Single cell capture

Sorted live CD45.2⁺ cells from YUMMUV1.7ie tumours were labelled with BD AbSeq Ab-Oligos (Supplementary Table S2) according to the manufacturer’s instructions (BD Doc ID: 23-24262(01)). Labelled cells were washed three times with BD Pharmingen™ Stain Buffer (500 × g, 10 min) and preserved in Omics Guard buffer (1 mL per 10⁴–10⁷ cells). Single-cell capture was performed using the BD Rhapsody™ Single-Cell Analysis System with the BD Rhapsody™ Scanner (BD Doc ID: 23-24252(01)). For a complete list of materials, see Supplementary Table 2. A total of 3.3–9.4 × 10⁴ cells per sample were captured, as determined by the BD Rhapsody Scanner.

#### Library preparation

Abseq and targeted mRNA libraries were generated according to the manufacturer’s instructions (BD Doc ID: 23-24514(01)) at Central Analytical Research Facility (CARF), Queensland University of Technology (QUT). Libraries were quantified and assessed for fragment size distribution using a high-sensitivity NGS Fragment Analyzer assay (DNF-474-33, 1 - 6000 bp; Agilent) prior to pooling and sequencing.

#### Sequencing

For each sample, AbSeq and targeted mRNA were uniquely indexed, pooled, and sequenced on a NovaSeq X (Illumina; 25B flow cell). Target sequencing depths were 5,000 reads per cell for AbSeq libraries and 15,000 reads per cell for targeted mRNA libraries. Following Illumina demultiplexing, separate FASTQ files were generated for each library type per sample and processed using the BD Rhapsody™ Targeted Analysis Pipeline (Seven Bridges platform; BD Doc ID: 23-24580(01)). Processing included read quality control, barcode and UMI extraction, alignment to the mm10 reference genome, gene assignment, and molecule counting with recursive substitution error correction (RSEC) and distribution-based error correction (DBEC).

#### Data analysis

DBEC count matrices were imported into R using Read10X and CreateSeuratObject (Seurat v4.5.1). RNA data were normalised using SCTransform prior to integration across samples. AbSeq antibody counts were normalised using centred log-ratio (CLR) transformation. Macrophages were extracted based on AbSeq expression of CD11B and F4/80. For macrophage subsets, RNA counts from the targeted panel were log-normalised and scaled prior to principal component analysis. Given the targeted 397-gene panel, all genes were used for downstream dimensionality reduction. UMAP was performed using the first 20 principal components, followed by graph-based clustering (resolution = 0.4). Cluster marker genes were identified using the Wilcoxon rank-sum test (FindAllMarkers). Genes detected in more than two clusters were excluded, and the top-ranked markers per cluster were used for annotation and heatmap visualisation.

### Statistical analysis

All statistical analyses were performed using GraphPad Prism 9 (RRID:SCR_002798). Bar graphs display mean values and standard deviation (SD).

### Data Availability

The original contributions presented in this study are included in the article/supplementary material. Raw and processed single cell CITE-seq and RNA-seq data described in the study have been submitted to the NCBI Gene Expression Omnibus repository and will be available upon manuscript acceptance. Further inquiries can be directed to the corresponding author(s).

## Results

### Tumour macrophage content correlates with tumour expressed cytokines

F4/80^+^ macrophages and monocytes can represent a high proportion of the tumour stroma. In mouse tumour models, we have found F4/80+ monocyte/macrophage content ranged from 20% of total live cell number in YUMMUV1.7 to ∼1 % in B16F10 melanomas in agreement with published ranges for these tumours (Figure 1A; Supp. Figure S1) [19–21]. Csf1 secretion <u>in</u> <u>vitro</u> was highest in the YUMMUV1.7 and lowest in B16F10, with the level of Csf1 expressed correlating with the level of macrophage content <u>in vivo</u> in four models including the ovarian cancer ID8 p53^-/-^ and three melanoma models (Figure 1B). Csf1 is a critical growth and survival factor for macrophages [5]. Depletion of Csf1 expression using shRNA (Clone A and Clone C) in the high expressing YUMMUV1.7 model reduced Csf1 to a level similar to YUMMUV3.3, but this reduction in Csf1 expression did not affect macrophage content <u>in vivo</u> (Figure 1C,D), although more complete depletion of Csf1 has been reported to reduce TAM content in other tumours [22]. These tumour lines also had strong basal expression of the macrophage chemo-attractants *Ccl2, Ccl5* and *Ccl7* expression, and although the levels of individual chemokines varied between tumours, each tumour exhibited high expression of at least one of these chemokines (Figure 1E) [23]. No expression of the other Csf1r ligand Il34 was found in any of the human or mouse models, and Il34 has been reported to have little contribution of TAM levels [22]. The strong correlation between Csf1 levels and TAM abundance suggests that tumour expressed Csf1 is a determinant of the level of tumour associated macrophage recruitment. However, the lack of effect on TAM content of Csf1 depletion in YUMMUV1.7 cells to levels comparable with other tumour lines suggests that either a minimal level of Csf1 expression is required for TAM recruitment and/or other tumour-derived tumour-expressed chemokines are also likely to contribute to recruitment.

**Figure 1:**
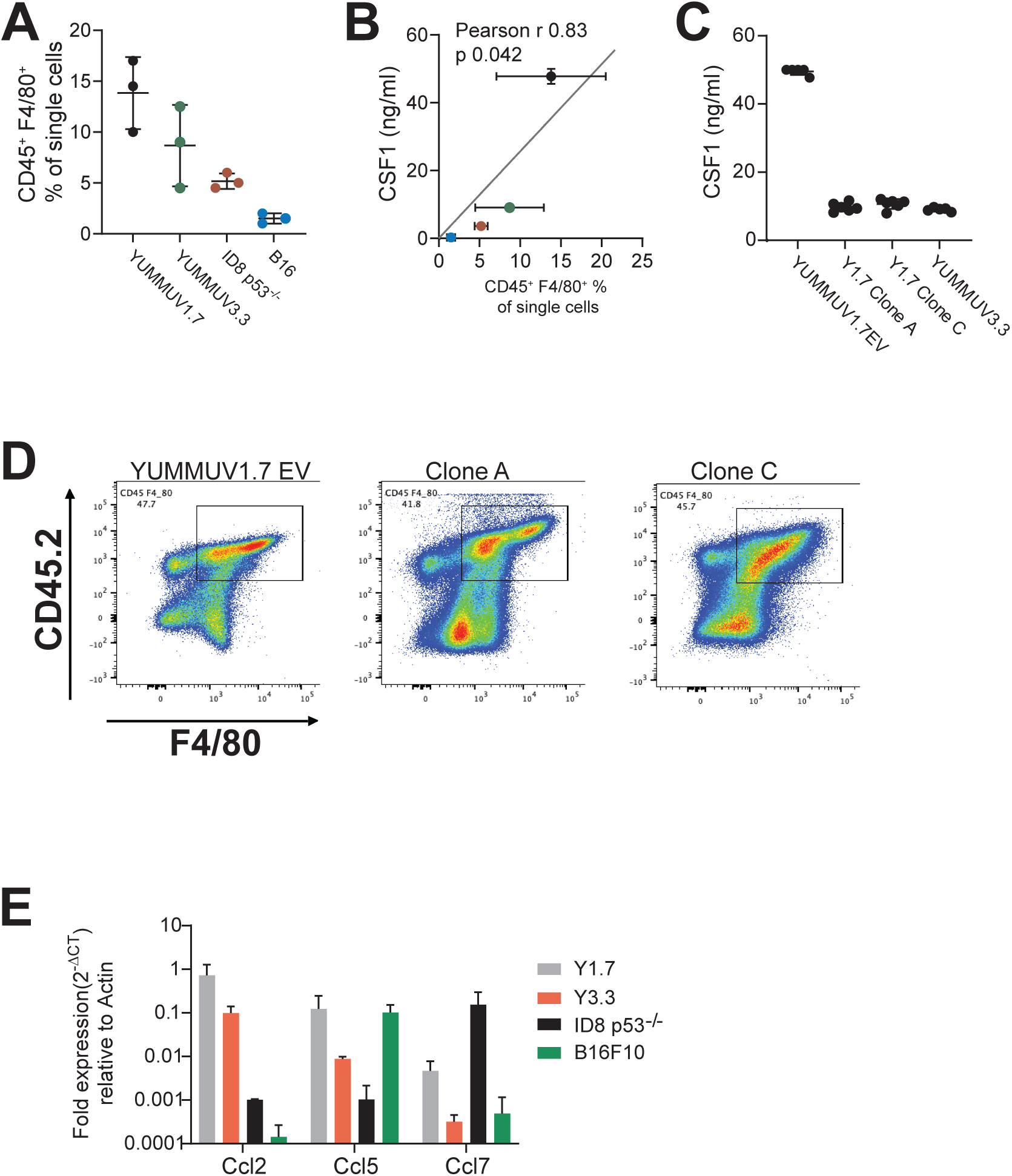
**A;** The proportion of CD45^+^ F4/80^+^ cells in the indicated syngenic mouse tumours of total cells. **B;** Correlation of secreted Csf1 in media plotted against the CD45^+^ F4/80^+^ population for mouse tumours in A. the indicated mouse tumour cell lines. The data are the mean and SD of 4 replicates for the Csf1 measurements. **C;** The secreted Csf1 levels in YUMMUV1.7 cells expressing empty vector (EV) or two different Csf1 shRNA (clone A, C) compared to YUMMUV3.3. **D;** Representative flow cytometry dot plots identifying the CD45^+^ F4/80^+^ cells in YUMMUV1.7 EV or Csf1 shRNA clone A and clone C expressing tumours. **E;** Expression level of the indicated chemokines from the indicated mouse tumour cell lines treated in in vitro with CHK1i+LDHU for 24h. The data is expressed relative to Actin and are the mean and SD from 4 replicates.

### Tumour-associated macrophages express Fcrg4

When analysing the tumour immune microenvironment in a panel of syngeneic mouse tumours, an antibody panel containing both lymphoid and myeloid markers identified populations of CD3^-^ TCRβ^-^ cells expressing the macrophage marker F4/80 and natural killer (NK) cell marker NK1.1 (Figure 2A, gate 1) in additional to F4/80^−^ NK1.1^+^ NK cells (gate 2). When the forward and side scatter profiles of the F4/80^+^ NK1.1^+^ and F4/80^−^ NK1.1^+^ cells were compared, the F4/80^−^ populations were smaller and had less complexity, whereas the F4/80^+^ NK1.1^+^ cells had size and complexity comparable to F4/80^+^ NK1.1^−^ macrophages (Figure 2B). Since tissue disaggregation can lead to macrophage fragmentation and erroneous identification of marker expression [24], Amnis imaging flow cytometry was used to further investigate these unexpected F4/80^+^ NK1.1^+^ cells using MacGreen mice that express EGFP from the *Csf1r* promoter. Gating for CD45^+^ NK1.1^−^ cells revealed lymphocyte sized CD3^+^ F4/80^−^ GFP^−^ T cells (Supp. Figure S2) while gating for the CD45^+^ NK1.1^+^ cells revealed similarly sized CD3^-^NK1.1^+^ F4/80^−^ GFP^−^ NK cells (Figure 2C). The NK1.1^+^ population also contained GFP ^Low/neg^F4/80^+^ and GFP^+^ F4/80^+^ populations that were significantly larger than the lymphocytes (Figure 2C). The NK1.1 staining was generally uniform and fully overlapped the F4/80 staining on the cells indicating this was not NK cell fragments but NK1.1 antibody binding to these large cells. This suggested that, despite the use of conventional Fc-blocking reagents, the NK1.1 antibody (a mouse IgG2a antibody) was binding to the Fcgr4 expressed on these large, predominantly Csf1r-GFP-expressing macrophages, as previously reported [25]. This was demonstrated by the NK1.1 and mouse IgG2a isotype control antibodies producing identical staining of the F4/80 population from tumours (Figure 2D). The presence of Fcgr4-expressing macrophages appeared to be specific for tumour as this was not found in the spleen of tumour bearing mice (Figure 2E). The IgG2a binding Fcgr4 (CD16.2) expressing macrophages were a common feature of multiple cancer models representing approximately 40% of all macrophages in range of tumour models including ascites of ID8 p53^-/-^ tumour bearing mice (Figure 2F).

**Figure 2:**
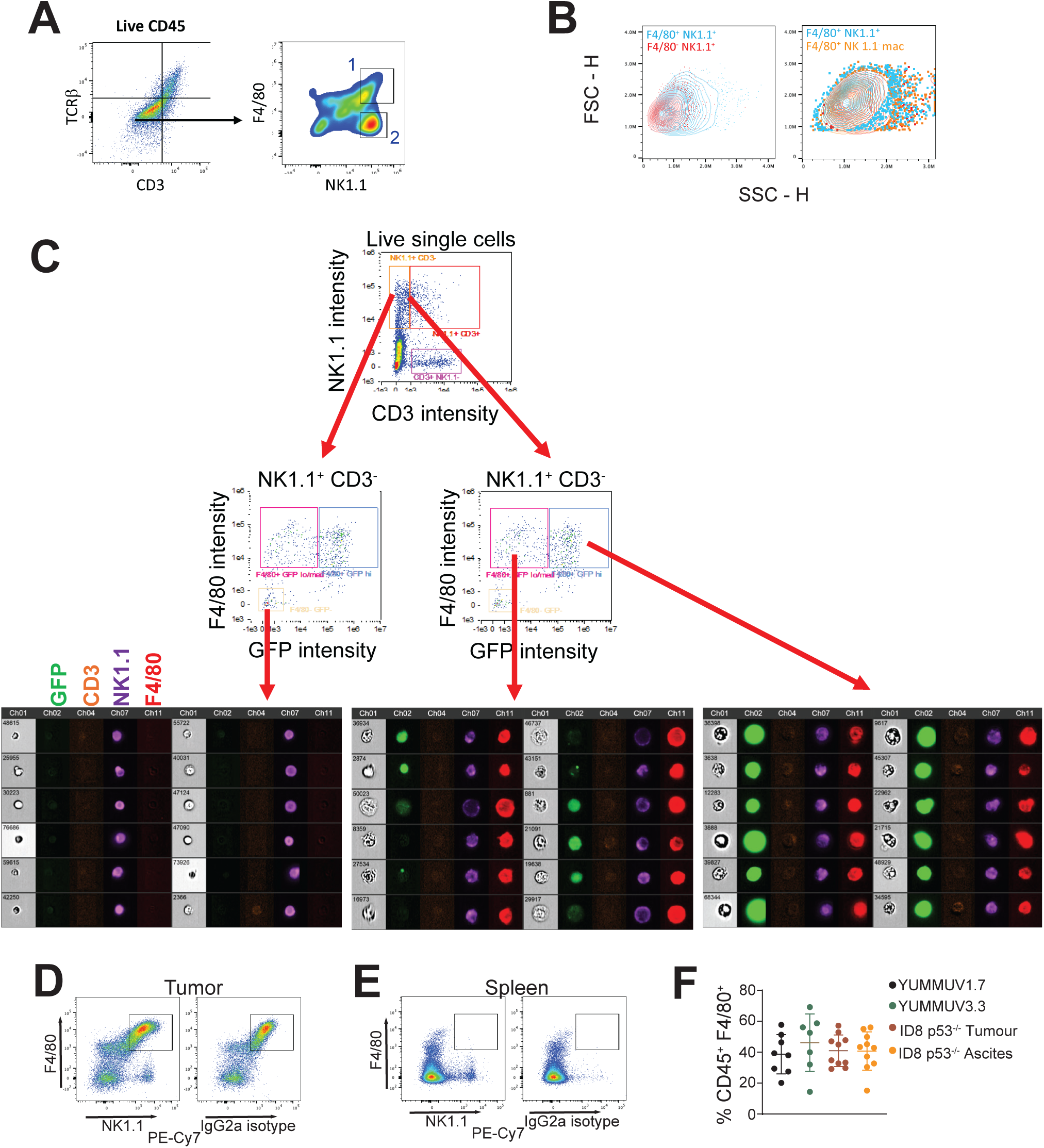
F4/80 expression on NK1.1^+^ populations. **A;** Mouse melanoma YUMMUV1.7 tumours were analysed using multiparameter flow cytometry. Representative gating strategy for gating live CD45^+^ cells. CD3- TCRβ- cells were analysed for F4/80 and NK1.1 staining. Gate 1 are F4/80^+^ NK1.1^+^ cells and gate 2 F4/80- NK1.1^+^ NK cells. **B;** Flow cytometry analysis of size (FSC) and complexity (SSC) comparisons of gate 2 F4/80- NK1.1^+^ NK cells compared to gate 1 F4/80^+^ NK1.1^+^ cells (left panel) and gate 1 F4/80^+^ NK1.1^+^ cells with F4/80^-^ NK1.1^-^ macrophages. **C;** Amnis imaging flow cytometry of CD45^+^ tumour associated cells from MacGreen mice. YUMMUV1.7 tumours were grown in MacGreen mice that express EGFP from the Csf1r promoter. The CD45+ cells were stained for CD3, F4/80 and NK1.1 and GFP fluorescence was used to identify Csf1r^+^ cells. The CD3^-^ NK1.1^+^ cells were gated for F4/80 and GFP staining intensity and images of cells from the indicated gates are shown. **D, E;** Flow cytometry of live CD45^+^ cells immunostained for F4/80 and either NK1.1 or IgG2a isotype antibody from YUMMUV1.7 tumour or normal spleen. **F;** The proportion of live CD45^+^ F4/80^+^ cells expressing FcRγ4 (identified by IgG2a binding) in the indicated mouse tumour models.

We performed CITEseq and single cell RNAseq (scRNAseq) of YUMMUV1.7 tumour derived CD45^+^ cells FACS sorted from control (62,980 cells) and CHK1i+LDHU treated mice (146,106 cells; Figure 3A). ScRNAseq was performed using BD Rhapsody imaging based single cell targeted mRNA amplification which selectively amplified 390 immune related and housekeeping genes (Supp. Figure S3, Supp. Table S3), and BD AbSeq for CITEseq detection of marker antibody binding to further investigate the nature of these Fcγr4-expressing macrophages. Initially, the control samples were analysed separately. In control tumours, CITEseq identified 10,379 F4/80^+^ CD11B^+^ macrophage/monocytes, and these were further divided into CD16.2/Fcgr4^+^ and ^−^ populations. Both populations expressed similar levels of the other indicated markers including CD16/Fcgr3 (Figure 3B). UMAP dimensionality reduction analysisof the control F4/80^+^ CD11B^+^ macrophage/monocyte of the scRNAseq data revealed a single major and several minor clusters, but overlaying CD16.2/Fcgr4 staining showed that cells expressing this marker were uniformly spread across all clusters as were the Fcgr4^-^ macrophages (Figure 3C), demonstrating little difference in gene expression patterns of these subsets (Supp. Figure S3). Only nine genes were significantly differentially expressed genes between the Fcgr4^+^ and ^−^ macrophage/monocyte populations, including increased expression of *Cxcl10*, *CD72*, *CD74*, *Irf7*, and *Stat1* which are associated with inflammatory macrophages [26, 27], and reduced expression of *Lmna* in Fcrg4^+^ macrophages (Figure 3D). However, the common highly expressed genes suggested the control TAM population possessed a dominant pro-tumoral phenotype [27–30], with strong expression of macrophage markers *Apoe*, *Lgals3*, *Trem2*, *Spp1*, *Arg1,* and *C1qa/b*, *Fn1*, *CD63* relatively uniformly across the clusters (Figure 3E, Supp. Figure S4), and low or no expression of pro-inflammatory markers such as *Cxcl9/10*, *Il1b*, *Lap3*, *Stat1*, *Tnfrsf13b*, although the Fcrg4+ cells express higher levels of *Cxcl10* and *Stat1* (Supp. Figure S3, Figure 3D). Only a few genes such as the MHCII genes e.g. *H2Aa* demonstrated enhanced expression in a subset of macrophages, whereas MHCI genes e. g. H2K1 were uniformly expressed (Figure 3E). Other pro-tumoral macrophage markers such type 2 chemokines *Ccl6*, *Ccl9* and *Cxcl16* [31] were also expressed by all macrophages (Supp. Figure S4). This suggests the Fcrg4^+^ macrophage have increased expression of a few pro-inflammatory markers, but the strong pro-tumoral signals in the control TAM population are likely to dominate.

**Figure 3:**
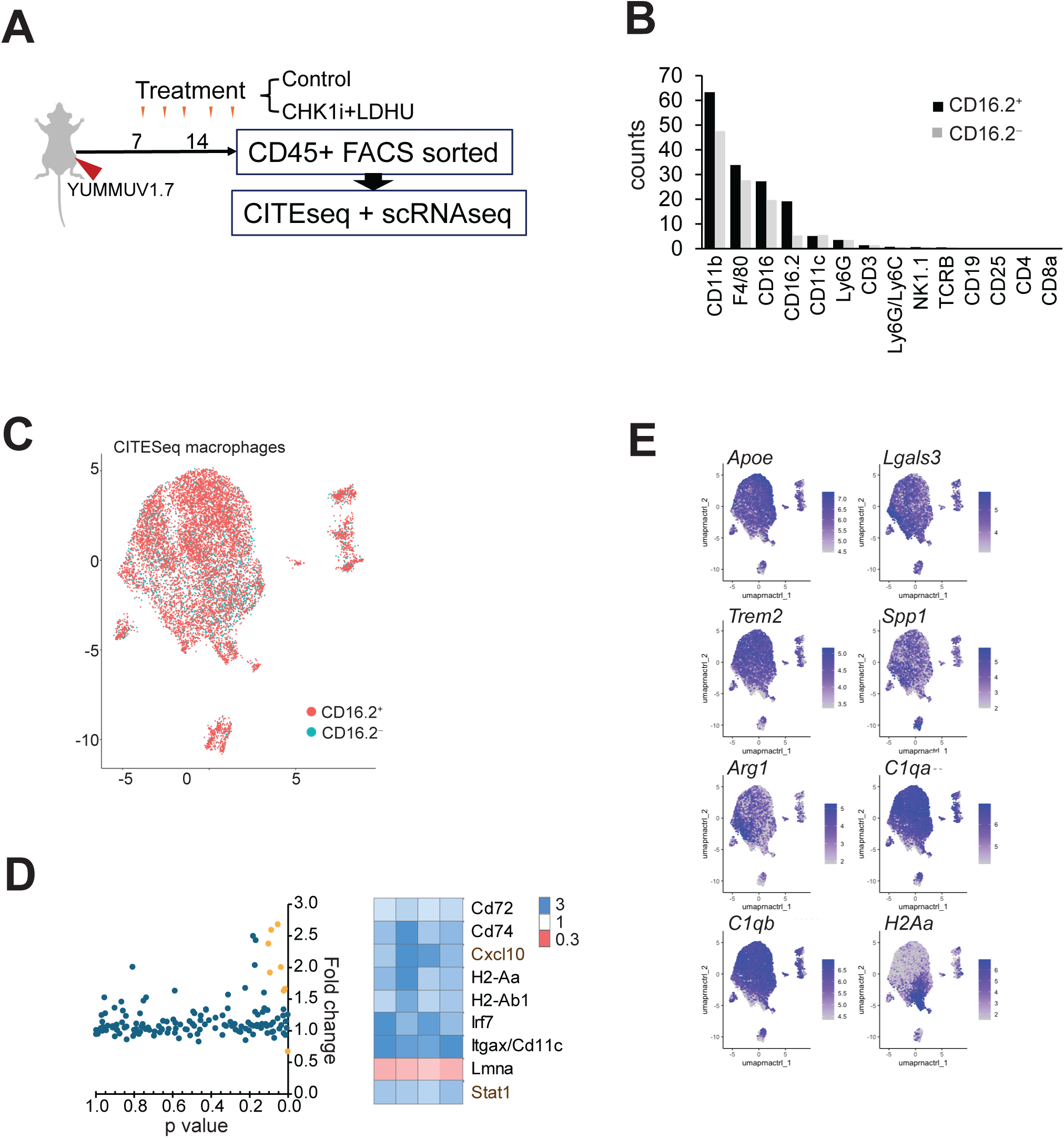
**A;** Diagram of the mouse experiment. **B;** CITEseq analysis of the CD11b^+^ F4/80^+^ monocyte/ macrophage populations (CD16.2^-^/FcRγ4^-^ and CD16.2^+^/FcRγ4^+^) from YUMMUV1.7 tumours for the indicated markers. The data are mean of 4 tumours. **C;** UMAP clustering from scRNAseq of control (untreated tumour) CD11b^+^ F4/80^+^ cells overlaid for CITEseq CD16.2^-^/FcRγ4^-^ and CD16.2^+^/FcRγ4^+^ staining. **D;** The dot plot of fold change against adjusted p value (t-test, left panel) in gene expression from the comparison of control FcRγ4^-^ and FcRγ4^+^ macrophages. These data are for the combination of all four control tumours. The yellow dots are significant (p<0.05) differentially expressed genes. Fold change of the significant differentially expressed genes in the individual tumours (right panel). **E;** Marker of the suppressive macrophage phenotype overlaid on the control CD11b^+^ F4/80^+^ macrophage UMAP clustering.

### Tumour targeted treatment reprograms macrophages phenotype

We have previously reported a combination of CHK1 inhibitor (CHK1i) with low dose of hydroxyurea (LDHU) demonstrated strong and selective anti-tumour activity in vitro [16], and in vivo triggered an anti-tumour immune response characterized by altered macrophage gene express suggestive of a change from pro-tumour to anti-tumour pro-inflammatory phenotype, for example decreased expression of *Mrc1*/CD206 and increased expression of *Nos2*/iNos [17].

To further investigate the effect of this treatment on the tumour associated macrophages, scRNAseq was performed comparing tumours from control and treated mice. CITEseq identified 10,379 control and 49,475 treated tumour-derived CD45^+^ CD11b^+^ F4/80^+^ cells. The difference in the numbers of cells was due to differences in batch processing and not increased number of macrophages with treatment. All CD45^+^ CD11b^+^ F4/80^+^ cells expressed high levels of macrophage markers *Apoe*, *C1qa/b*, *Ccr1*, *CD63*, *Fcgr3*, *Lgals1*, *Lyz2*, *Spp1*, and immunosuppressive genes e.g. *Arg1*, *Ccl2/9*, *CD52*, *Ctsd*, *Hmox1*, *Lagals3* and *Trem2* [30] (Supp Table S3; Supp Figure S3). Many of the strongly suppressive markers (*Apoe, C1qa/b, Ctsd, Lgals1, Spp1*) were relatively unchanged with treatment (Supp Figure S3). UMAP analysis revealed 12 clusters in the scRNAseq data (Figure 4A). Most macrophages from untreated mice clustered tightly together (Figure 4B, cluster 2), which appears similar to the major cluster in the control only UMAP (Figure 3C). Cluster 2 was distinguished from the other clusters by high expression of suppressive markers such as *CD163*, *Fcna*, *F13a1* and *Maf* and strongly downregulated expression of the inflammatory markers *Irf7*, *Stat1* and MHC-I *H2-K1* (Figure 5C; Supp. Figure S5; Supp. Table S4). Cells from treated mice formed a large cluster comprising: clusters 0 and 1, characterized by suppressive genes *Arg1* and *Mmp12* respectively (Figure 5C; Supp Figure S5; Supp Table S4); cluster 3 with increased expression of proliferative markers such as *Aurkb*, *Mcmc2/4*, *Tyms*, *Ube2c*, *Pclaf*; and cluster 4 with modestly increased expression of both inflammatory markers *Nlrp3* and *Tnf* and suppressive markers *Il4ra, Cd72, C1qa/b* indicating a mixed response. Macrophages from treated mice were also enriched in clusters 5, 6 and 11, which were separated from the major clusters and characterized by strongly increased expression of NK (cluster 5; *Gzma/b, Prf1, Klra1/3/7/21, Nkg7*), T cell (cluster 6; *CD3, CD8, Trac, Lat, CD247*) and B cell (cluster 11; *CD19, Igkc, Mzb1, Pou2af1*) genes. As these genes are expressed at extremely low levels in control macrophage populations (Supp Table S3), the very strong enrichment (up to 70-fold) in macrophages from treated mice to levels that could be readily detected (Figure 4D), whereas tumour associated macrophage markers such as *C1qa*, *Lyz2* and *Lgals1* were strongly expressed in all clusters (Figure 4E). This suggested that the macrophages are directly in contact with the NK, T and B cells in treated mice and co-isolated during tissue disruption. The presence of *Gzma/b* and *Prf1* in the NK cluster 6 and *Gzmb*, *Icos* and *PdCD1* in the T cell cluster 5 suggest the NK and T cells may be activated (Figure 4D). Cluster 7 was characterised by markers of multiple stromal cell type fibroblast (*CD34*, *Tgfb3*) and neutrophils (*Mmp9*) (Supp Table S3). Cluster 8 appears to have a set of genes that are induced through Tlr4 signalling (*S100a8/9, Mmp9, Tnf*); cluster 9 is marked by increase possible dendritic and mast cells markers *Flt3, Ccl17*, *CD40*, *CD86* and *Kit* expression (Figure 5C; Supp Figure S5; Supp Tables S4); cluster 10 is strongly pro-inflammatory with increased expression of *Nlrp3, Il6ra, Nod2, Casp1*. Together, the data suggest that treatment of tumours with CHK1i+LDHU reprogrammed the macrophage phenotype to a more pro-inflammatory, anti-tumour phenotype having increased interaction with effector cells such as NK and T cells.

**Figure 4:**
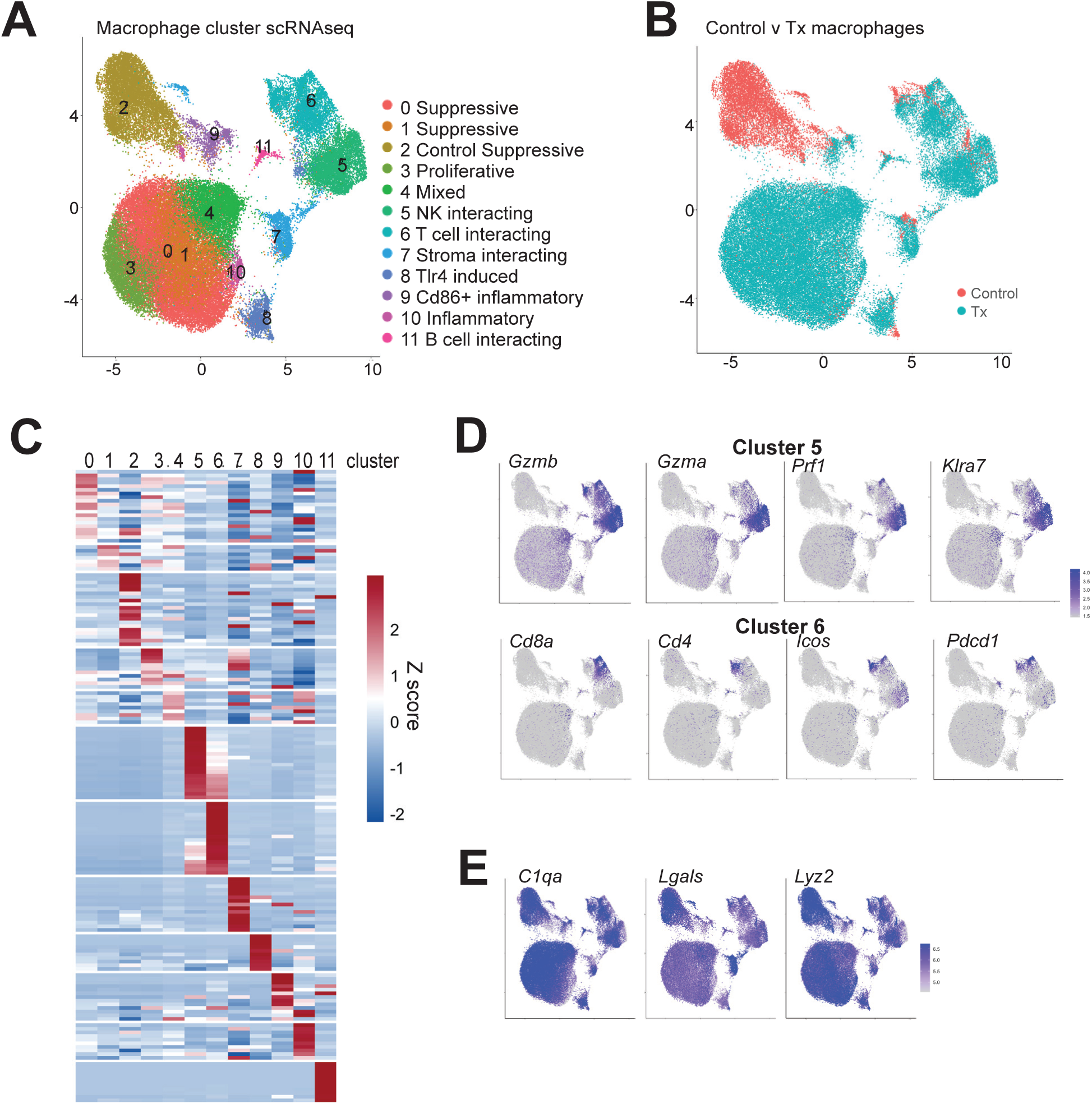
**A;** UMAP clustering of scRNAseq data of macrophages from control and treated YUMMUV1.7 tumours (n=4 for each arm). **B;** The control and CHK1i+LDHU treated (Tx) cluster are identified. **C;** Heat maps showing the z score of gene expression in the individual clusters compared to the rest of the clusters combined. A full-size version of this heatmap with gene names is shown in Supp Figure S5). **D;** Overlay of the indicated T and NK cell markers on the macrophage UMAP. The scale bar indicates the log2 gene expression. **E;** Overlay of the indicated macrophage gene expression on the macrophage UMAP. The scale bar indicates the log2 gene expression.

**Figure 5:**
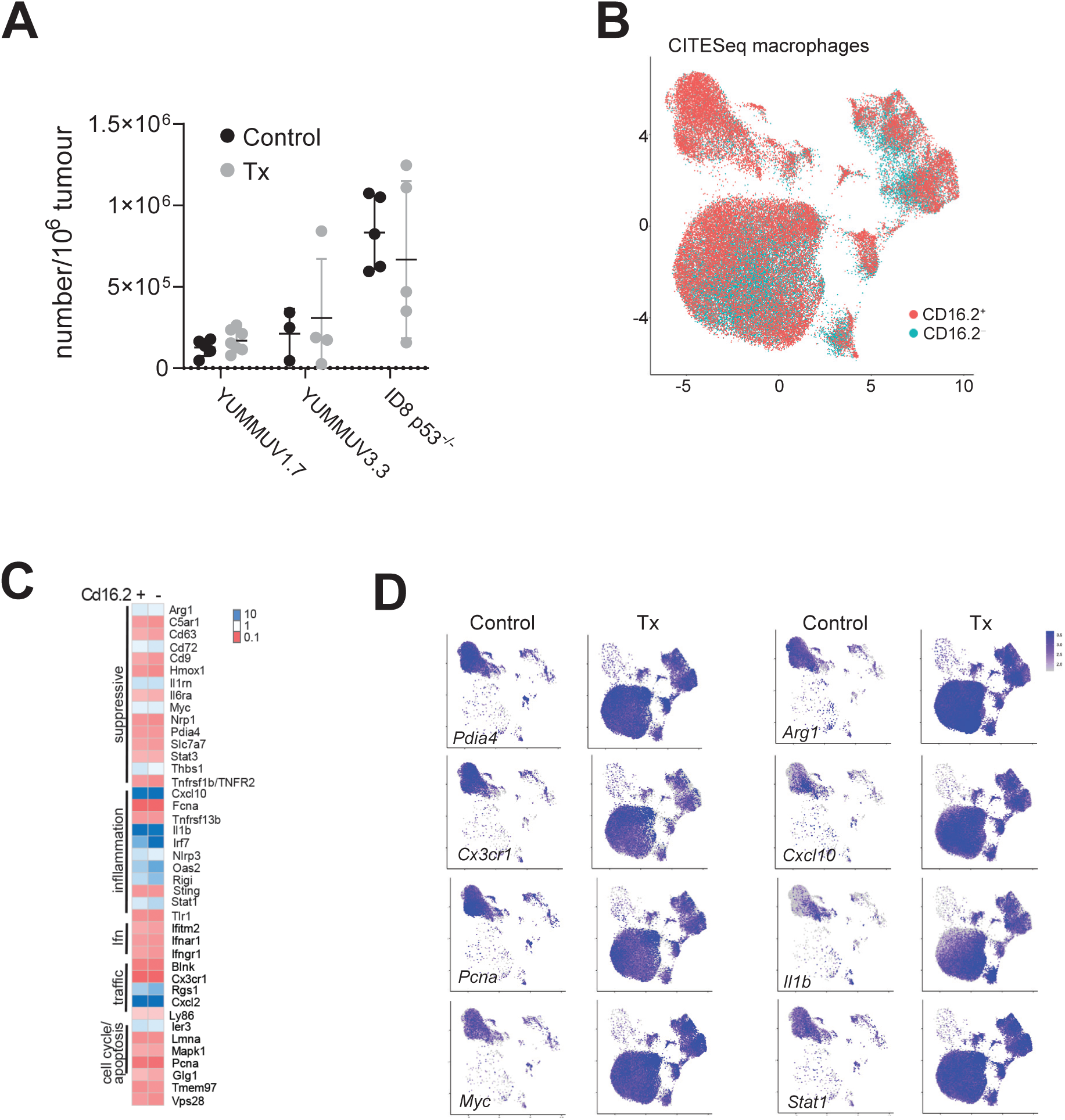
**A;** The numbers of FcRγ4^+^ macrophages (identified as CD45^+^ F4/80^+^ CD11b^+^ IgG2a^+^) from the indicated control and treated tumours. **B;** Overlay of the CITEseq data for CD16.2/FcRγ4 status on the macrophage UMAP clustering. **C;** Differentially expressed genes in control versus treated tumours (>2 fold change p>0.05 multiple 2 tailed t tests, mean of replicate tumours) for the FcRγ4^+^ and FcRγ4^-^ macrophages. **D;** Overlay of a subset of the differentially expressed genes onto the macrophage UMAP clustering. The control and treated (Tx) UMAPs are shown for each gene.

CHK1i+LDHU treatment had little effect on the numbers of Fcrg4^+^ macrophages across multiple tumour types (Figure 5A), and as in control mouse tumours, Fcrg4^+^ macrophages did not cluster selectively compared to Fcrg4^-^ (Figure 5B). When the differentially expressed genes between control and treated macrophages were assessed, there was little difference between the Fcrg4 populations (Figure 5C). Surprisingly, many of the strongly altered genes were not present as cluster markers, but visualization in the UMAP showed strong enrichment of treatment-upregulated genes e.g. *Arg1*, *Cxcl10*, *Myc* and *Stat1* across all treatment-specific clusters, and enrichment of down-regulated genes e.g. *Pdia4* and *Cx3cr1*. *Pcna* was down-regulated in most treatment-specific clusters except in the proliferative cluster 3, and *Il1b* had very low expression in parts of clusters 0 and 3 (Figure 5D). A number of genes that are commonly used as markers of macrophage phenotypes such as *C1qa* and *Spp1* were represented in the scRNAseq panel used here, but only *C1qa* was found selectively expressed in any of the clusters in this study (Supp. Table 4; clusters 0,3,4). Mapping expression of these genes on the UMAP revealed they are generally expressed, although there were modest increases in *C1qa* levels in its assigned clusters (Supp. Figure S6). This expression pattern was followed many genes including *Hmox1*, *Cd63*, *Cd86* and *Fcgr3*. *Cd163* and *Irf7* displayed expression profiles supporting their down and up-regulation, respectively, with treatment, the latter across all treatment related clusters. The MHC I gene *H2K1* also displayed general treatment-dependent up-regulation, whereas MHC II *H2Aa* displayed more selective high expression in multiple clusters (Supp. Figure S6). Together, this suggested that whereas macrophages retain much of their suppressive character in treated tumours, they have gained pro-inflammatory characteristics e.g. *Cxcl10*, *Il1b* expression, and downregulated pro-tumour signals e.g. *Cd63*, *Hmox1*.

To determine the contribution of the tumour-associated macrophages to the CHK1i+LDHU treatment response, we used a Csf1r neutralising antibody to depleted monocyte/macrophages prior to and during treatment to assess the effects on tumour growth and immune response (Figure 6A). Csf1r antibody treatment depleted >95% of F4/80^hi^ CD11b ^hi^ macrophages and F4/80^int^ CD11b^int^ Mo-MDSC in the YUMMUV1.7 model (Figure 6B box 1, 2; Figure 6C), but not F4/80^int^ CD11b^hi^ cells lacking Csf1r expression, possibly classical dendritic cells (cDCs) [32]. Csf1r depletion completely depleted F4/80^+^ NK1.1 antibody binding Fcrg4^+^ macrophages (Supp. Figure 7A box 1, Figure 6C) but had no effect on the F4/80^-^ NK1.1^+^ NK cell (box 2). The two-fold increase in NK cells is likely to simply reflect the 50% reduction in total tumour associated CD45^+^ cells with anti-Csf1r antibody treatment (Supp. Figure 7B). Csf1r depletion of macrophages had no significant effect on tumour growth but significantly enhanced the anti-tumour effects of treatment (Figure 6D). We have previously demonstrated that CHK1i+LDHU treatment triggers a CD8^+^ T cell dependent immune response [17, 18]. Analysis of the CD8^+^ T cell population from control and treated tumours showed that CHK1i+LDHU treatment increased the proportion of CD8^+^ T cells expressing PD-1, a marker of T cell activation, and the level of expression (Figure 6E). Depletion of Csf1r^+^ monocyte/macrophages enhanced the number of PD1^+^ CD8^+^ T cells, which increased further with treatment. The number of granzyme B (GrB) expressing CD8^+^ T cells, a marker of T cell effector function, was also significantly increased with CHK1i+LDHU treatment, and likewise increased with macrophage depletion, but increased further with treatment and macrophage depletion. Macrophage depletion did not augment the impact of CHK1i+LDHU treatment on the measured CD8 T cell parameters, indicating an alternative mechanism underlying the synergistic anti-tumour effect. Together the data indicate that the anti-proliferative effect of CHK1i+LDHU treatment was required in combination with T cell activation to inhibit tumour growth, and despite treatment reprogramming of the TAM population to a more anti-tumour phenotype, the dominant effect of macrophages remains immune suppressive.

**Figure 6:**
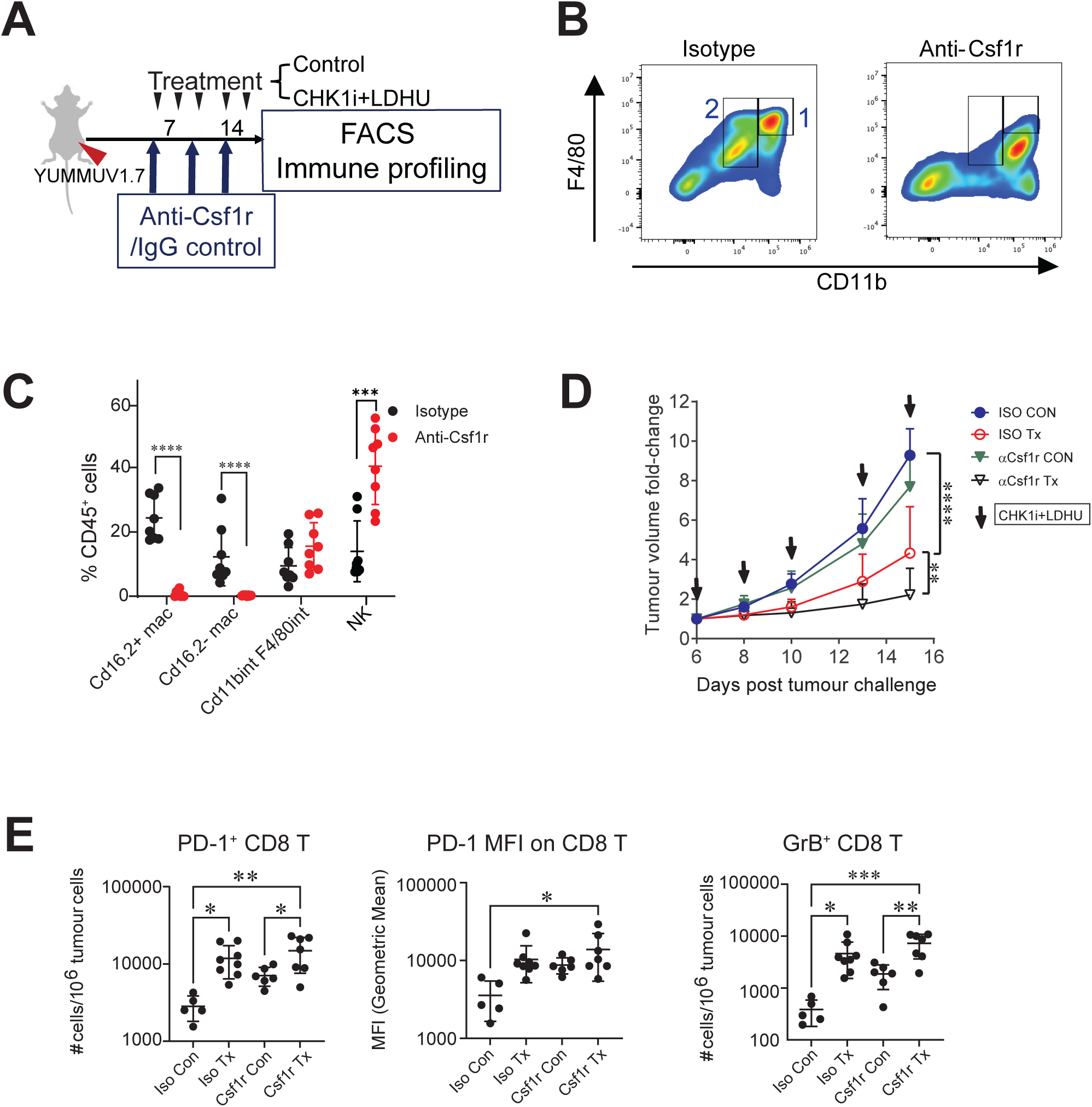
**A;** Diagram of the mouse experiment. Mice bearing YUMMUV1.7 tumours were treated with either isotype or anti-Csf1r antibodies to deplete Csf1r expressing macrophages and myeloid cells. Mice were also treated without or with CHK1i+LDHU and tumour growth measured. After five CHK1i+ LDHU treatments, tumours were excised and analysed for the tumour-associated immune cells. **B;** Flow cytometry of live CD45^+^ cells stained for F4/80 and CD11b. Gates 1 and 2 show the Csf1r antibody depleted F4/80^+^ CD11b^+^ population has been further subdivided into intermediate (int) and high (hi) staining. **C;** Numbers of CD11b^+^ F4/80^+^ cells in YUMMUV1.7 tumours from either isotype antibody or anti-Csf1r treated mice. The F4/80hi CD11bhi and CD11bint macrophages and myeloid cells were further divided on the IgG2a binding as CD16.2/FcRγ4^+^ and CD16.2/FcRγ4^-^. The F4/80^-^NK1.1^+^ NK cells were defined as in Supp. Figure S6A. The number cells are presented as cells/million tumour cells. The data are from 4 control and 4 CHK1i+LDHU treated for isotype and anti-Csf1r antibody treatments. **D;** Tumour growth of YUMMUV1.7 tumour either without (isotype contorl; ISO) or with CSF1R antibody depletion (αCsf1r) and without (Con) or with CHK1i+LDHU treatment (TX). This indicate treatments. The data are the mean and SD for 4 mice for each arm. Data was analysed by 2way ANOVA using Tukey’s multiple comparisons test. ** p<0.01, ****p<0.0001. **E;** The number of CD8^+^ T cells expressing PD-1, level of expression (mean fluorescence intensity MFI) of PD-1 on CD8^+^ T cells and number of CD8^+^ T cells expressing the markers of immune activity Granzyme B (GrB), for each treatment group. The data are the mean and SD for 5-7 mice. Data was analysed by 2way ANOVA using Tukey’s multiple comparisons test. * p<0.05, ** p<0.01, ***p<0.001.

## Discussion

TAM are a common feature of tumours and often major stromal component that has a major role in imposing an immunosuppressive tumour microenvironment. The question of what is responsible for the differing TAM content of tumours has not been unequivocally answered here. We have found that TAM content was significantly correlated with the level of tumour expressed Csf1, and tumours also variably expressed other chemokines that can act as attractants for macrophages suggesting that multiple factors likely contribute to the overall TAM content as suggested by the reduced TAM content in CCR2^-/-^ mice [33]. The alternative Csf1r ligands Il34 was not expressed and has no contribution to TAM levels as reported previously [22].

The discovery that a high proportion of TAM express Fcrg4 was surprising. This uncommon Fc receptor is not blocked by the commercial Fc blocking agents commonly used [25], but CD16.2/Fcgr4 is normally present on a minor population of lung interstitial monocytes [34] and its expression is strong increased on macrophages with cyclophosphamide treatment [35]. In TAM, the expression of Fcgr4 had only a minor effect on their phenotype, but the lack of these in other lymphoid tissues suggests that the increase expression is consequence of the tumour microenvironment. Fcrg4 has moderate affinity for IgG2a and IgG2b isotypes and is responsible for the therapeutic efficacy of IgG2a antibody treatment mediated through Antibody-Dependent Cellular Cytotoxicity (ADCC) and Antibody-Dependent Cellular Phagocytosis [36]. This is relevant not only for antibody-mediated therapies but may also explain additional contribution of these cells to the efficacy of the combination treatment. One potential consequence is that testing antibody therapies in mouse cancer models may be confounded by the high-level expression of this unexpected Fc receptor in tumours. The large numbers of CD16.2/Fcgr4^+^ macrophages found in tumours may also provide a critical mechanistic link to a potential humoral (antibody-mediated) antitumor response. CHK1i+LDHU has been demonstrated to trigger immunogenic cell death in all tumour models tested [17, 18], and the release of tumour antigens by the treatment may prime B cells to produce endogenous tumour-specific antibodies [37] that may then trigger a Fcrg4-dependent phagocytic response. However, the enhanced anti-tumour effect of macrophage depletion indicates that the main effect of TAM remains immunosuppressive even with treatment. The human orthologues of mouse Fcrg4, FCGR3A and FCGR3B, are commonly expressed on tumour associated NK cells and macrophages and have been shown to contribute to the efficacy of immunotherapies [38–40]. Natural variants of *FCGR3A* influence therapy responses to antibody treatments [41] demonstrating the potential importance of this Fc receptor in antibody therapeutic responses. Thus, while not previously recognised as being a major component of mouse tumour stroma, the presence of high levels of Cd16.2/Fcgr4 in mouse tumour expressed on macrophages mimics human tumours, although we did not find Fcrg4 expression on NK cells in mouse tumours.

Many groups have published different TAM clusters defined by expression of specific marker genes such as *C1qa* and *Spp1* [28–30, 42, 43]. We have not identified similar macrophage clusters, although on the basis of more generalised clusters suggested by Ma and colleagues [30], clusters 0 and 2 have regulatory -TAM (Reg-) like expression signatures with cluster 2 (control TAMs) also possessing some lipid associated-TAM) and monocyte characteristics such as expression of *Fcgr3*/Cd16, *F13a1*, *Maf* and *Mgst1.* The Reg-TAM resemble alternatively activated macrophages, pointing to their strongly suppressive properties. The monocyte-like properties of the control cluster 2 TAMs may reflect the possible circulating monocyte origin of this population [30]. Subtype markers common reported such as *Spp1* [30, 42, 43] were found generally expressed across all clusters in this study, whereas others such as *C1qa* showed some selective accumulation although all clusters had significant expression. One difference between this and published studies is the gene set interrogated here represented ∼400 immune-related genes rather than whole exomes used in the majority of published studies, possibly limiting the clusters that can be defined. Another important distinction is all the published studies were analysis of TAMs in untreated tumours and define subsets of immune suppressive TAM, whereas the TAM clustering in this study was primarily driven by the response to CHK1i+LDHU. While specific molecules were found to be strongly upregulated with treatment, for example MHC I molecules such as H2K1 were strongly upregulated with CHK1i+LDHU treatment across all treatment defined clusters whereas MHC II molecules such as H2Aa accumulated only in specific regions of control and several treatment-defined clusters. We have previously reported that CHK1i+LDHU promotes strong PD-1 expression on CD8^+^ T cells <u>in vivo</u>, suggestive of an exhausted state [17]. The overall effect of CHK1i+LDHU treatment was to drive macrophages from a pro-tumour immune suppressive (clusters 2) to a less suppressive (clusters 0, 1) and inflammatory (clusters 9, 10) with cluster 4 representing an intermediate state. It also promoted proliferation (cluster 3). Clusters with high level expression of markers of other immune cell types, T, NK and B cells (clusters 5, 6, 11) suggests that with treatment macrophages may be acting as antigen presenting cells (APCs). TAM were shown to be responsible for T cell exhaustion states through their MHC I expression and their high numbers in tumour compared to other tumour associated APCs [33]. Treatment also enhanced interactions with other stromal components (cluster 7). However, the overall effect of TAM remained immune suppressive, demonstrated by Csf1r antibody depletion increasing response to treatment. Macrophage depletion alone was sufficient to increase markers of CD8^+^ T cells activation, providing further functional evidence of immunosuppressive macrophage phenotype, but this was not sufficient to influence tumour growth. This matches previous studies that have demonstrated targeting TAM using either CSF1R inhibitor or CSF1R antibody depletion increases treatment efficacy in range of tumour types and treatments [44–47], and the failure of TAM targeted treatments as single agents in human clinical trials [15, 48].

The data presented here demonstrates that CHK1i+LDHU can promote reprogramming of TAM to a more inflammatory anti-tumour phenotype, however, this was not sufficient to fully reprogram TAMs and combination with another therapy that drives this reprogramming enhance the inflammatory phenotype is required to maximise therapeutic benefit.

## Acknowledgements

ID8-p53WT, ID8-p53^−/−^ were kindly provided by Professor Roby (Kansas University Medical Center).

## Author Contributions

Conceptualization, ZZ, BG.; methodology, ZZ, RB, AG, MP, JG, SM; validation, ZZ, AG, RB, MP; formal analysis, ZZ, AG, RB, JG, BG.; investigation, ZZ, AG, RB, MP, JG; resources, SW, KI; data curation, ZZ, AG,MP; writing—original draft preparation, ZZ, AG, RB, BG.; writing—review and editing, ZZ, AG, RB, MP, SM, SW, RD, KI, JW, JGC, BG.; visualization, ZZ, AG, JG, BG.; supervision, ZZ, RB, SM, BG.; project administration, BG.; funding acquisition, RD, JW, JGC, BG. All authors have read and agreed to the published version of the manuscript.

## Ethics

Animal ethics was obtained from The University of Queensland Molecular Biosciences Animal Ethics Committee - 2021/AE000025 and The University of Queensland Health Sciences Animal Ethics Committee – 2022/AE000652.

