## Supplementary Methods, Tables, Figures for "Reprogramming tumour-associated macrophages from immune suppressive to inflammatory state by Checkpoint kinase 1 inhibitor combination treatment"

### Supplementary Material

#### Supplementary Methods

##### *Spectral unmixing for Cytek flow cytometry*

Spectral unmixing was performed in SpectroFlo (RRID:SCR_025494) using the manufacturer's standard unmixing workflow. Single-stained compensation controls for every fluorochrome in the panel were used to generate reference emission spectra. An unstained cell control was included to capture and model the sample-specific autofluorescence profile, which was incorporated automatically by SpectroFlo's autofluorescence extraction algorithm during unmixing. This approach allowed subtraction of the autofluorescence signal across the full spectrum, improving resolution of dim markers and ensuring accurate separation of fluorochromes. All samples were unmixed using the same reference controls within the same acquisition session to maintain consistency and reproducibility. Acquired data were analysed using FlowJo software, RRID:SCR_008520.

Table S1: Flow cytometry antibodies

| Antibody | Supplier (Cat#) | RRID |
|---|---|---|
| ***Immune profiling panel*** | | |
| TruStain FcX™ (anti-mouse CD16/32) | Biolegend (101320) | AB_1574975 |
| CD11b-BV421 | Biolegend (101236) | AB_11203704 |
| CD11b-BV650 | Biolegend (101259) | AB_2566568 |
| CD19-APC-Cy7 | Biolegend (115529) | AB_830706 |
| CD19-BV785 | Biolegend (115543) | AB_11218994 |
| CD3-APC | biogems (05112-80) | AB_1272181 |
| CD3-BUV395 | BD Horizon™ (740268) | AB_2687927 |
| CD3-FITC | Biolegend (100204) | AB_312661 |
| CD4-Alexa Fluor 700 | Biolegend (100536) | AB_493701 |
| CD4-BUV395 | Invitrogen (363-0042-82) | AB_2920941 |
| CD45.2-PE/Dazzle 594 | Biolegend (109846) | AB_2564177 |
| CD8α-BV605 | Biolegend (100744) | AB_2562609 |
| F4/80- Alexa Fluor 488 | Biolegend (123120) | AB_893479 |
| F4/80-BV711 | Biolegend (123147) | AB_2564588 |
| FOXP3-Alexa Fluor 647 | Biolegend (126408) | AB_1089115 |
| Gr-1-PercpCy5.5 | Biolegend (108427) | AB_893561 |
| Ly6C-APC | Biolegend (128015) | AB_1732087 |
| Ly6G-Alexa Fluor 700 | Biolegend (127622) | AB_10643269 |
| MHCII-APC-Cy7 | Biolegend (107628) | AB_2069377 |
| Mouse IgG2a kappa Isotype- PE-Cy7 | Invitrogen (25-4724-81) | AB_470203 |
| NK1.1-PE-Cy7 | Biolegend (108714) | AB_389364 |
| NK1.1-PE-Cy7 | eBiosciences (25-5941-82) | AB_469665 |
| PD-1-BV785 | Biolegend (135225) | AB_2563680 |
| PD-L1-PE | Biolegend (155404) | AB_2728223 |
| TCRβ-PE | Biolegend (109207) | AB_313430 |
| TCRβ-Percp-Cy5.5 | Biolegend (109228) | AB_1575173 |
| ***Imaging flow cytometry panel*** | | |
| CD115-BV605 | Biolegend (135517) | AB_2562760 |
| CD3-PE/Dazzle 594 | Biolegend (100347) | AB_2564028 |
| F4/80- Alexa Fluor 647 | BD Pharmingen™ (565853) | AB_2744474 |
| NK1.1-BV421 | Biolegend (108732) | AB_2562218 |

Table S2: BD reagents

| Catalogue # | Description |
|---|---|
| 666262 | BD Rhapsody™ 8-Lane Cartridge |
| 667052 | BD Rhapsody™ Enhanced Cartridge Reagent Kit V3 |
| 633773 | BD Rhapsody™ cDNA Kit |
| 633774 | BD Rhapsody™ Targeted mRNA and AbSeq Amplification Kit |
| 633753 | BD Rhapsody™ Immune Response Panel Mm |
| 667059 | BD Rhapsody™ Mouse TCR/BCR Next Amplification Kit |
| 940111 | BD™ AbSeq Oligo Rat Anti-Mouse CD19, Clone: 1D3 |
| 940471 | BD™ AbSeq Oligo Rat Anti-Mouse CD4, Clone: GK1.5 |
| 940128 | BD™ AbSeq Oligo Hamster Anti-Mouse CD279 (PD-1), Clone: J43 |
| 940345 | BD™ AbSeq Oligo Rat Anti-Mouse CD8a, Clone: 53-6.7 |
| 940327 | BD™ AbSeq Oligo Hamster Anti-Mouse CD3e, Clone: 500A2 |
| 940131 | BD™ AbSeq Oligo Rat Anti-Mouse F4/80, Clone: T45-2342 |
| 940321 | BD™ AbSeq Oligo Hamster Anti-Mouse CD11c, Clone: N418 |
| 940356 | BD™ AbSeq Oligo Rat Anti-Mouse CD25, Clone: 3C7 |
| 940121 | BD™ AbSeq Oligo Mouse Anti-Mouse NK-1.1, Clone: PK136 |
| 940113 | BD™ AbSeq Oligo Rat Anti-Mouse Ly-6G, Clone: 1A8 |
| 940142 | BD™ AbSeq Oligo Rat Anti-Mouse CD274, Clone: MIH5 |
| 940119 | BD™ AbSeq Oligo Rat Anti-Mouse Ly-6G and Ly-6C, Clone: RB6-8C5 |
| 940125 | BD™ AbSeq Oligo Hamster Anti-Mouse TCR ß Chain, Clone: H57-597 |
| 940008 | BD™ AbSeq Oligo Rat Anti-CD11b, Clone: M1/70 |
| 940340 | BD™ AbSeq Oligo Rat Anti-Mouse CD370 (Clec9A), Clone: 10B4 |
| 940191 | BD™ AbSeq Oligo Mouse Anti-Mouse TIGIT, Clone: 1G9 |
| 940140 | BD™ AbSeq Oligo Rat Anti-Mouse CD335 (NKp46), Clone: 29A1.4 |
| 940109 | BD™ AbSeq Oligo Rat Anti-Mouse CD16/CD32, Clone: 2.4G2 |
| 940338 | BD™ AbSeq Oligo Hamster Anti-Mouse FcγRIV (CD16-2), Clone: 9E9 |
| 940198 | BD™ AbSeq Oligo Rat Anti-Mouse CD115 (CSF-1R), Clone: T38-320 |

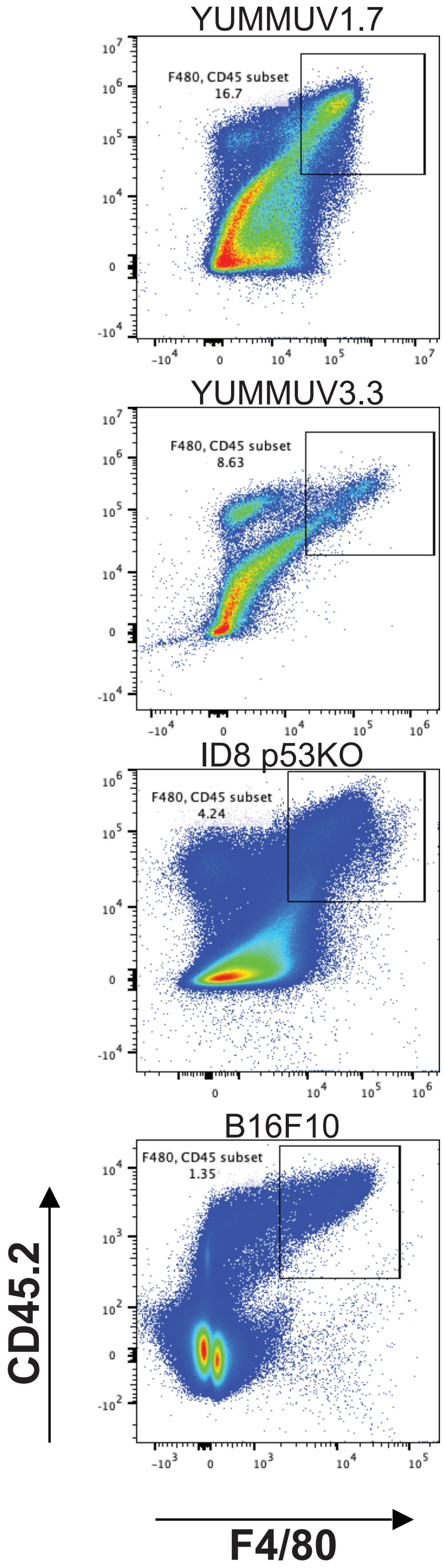


**Supplementary Figure S1:** Representative flow cytometry dot plots identifying the CD45.2⁺ F4/80⁺ cells in the indicated tumours.

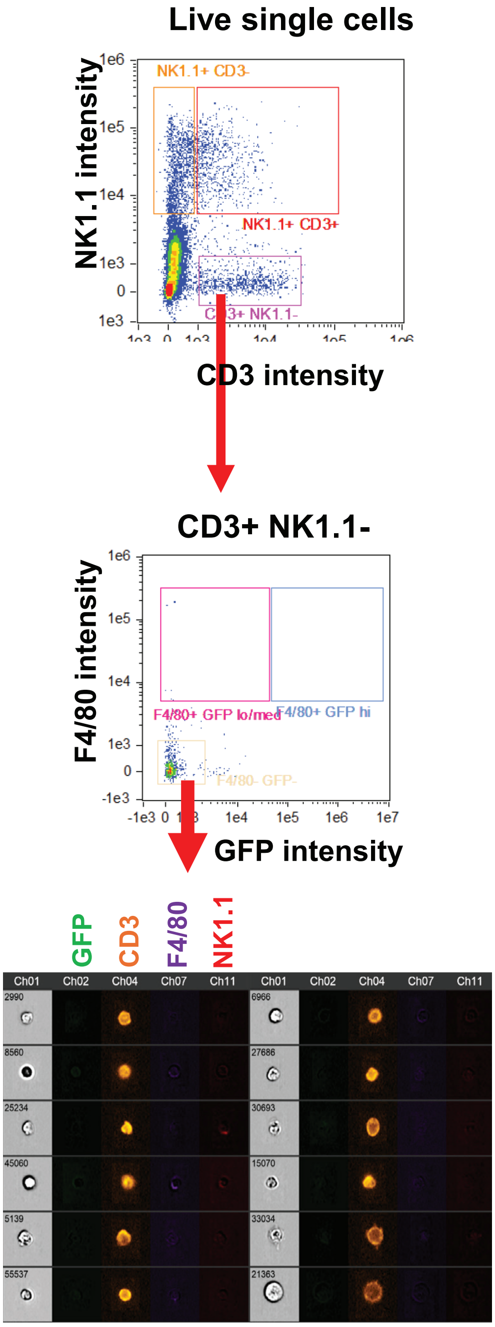


**Supplementary Figure S2:** Amnis imaging flow cytometry of CD45$^+$ tumour associated cells from MacGreen mice. YUMMUV1.7 tumours were grown in MacGreen mice that express EGFP from the Csf1r promoter. The CD45+ cells were stained for CD3, F4/80 and NK1.1 and GFP fluorescence was used to identify Csf1r+ cells. The images of cells from the CD45$^+$ CD3$^+$ NK1.1$^-$ F4/80$^-$ GFP$^-$ T cells are shown.

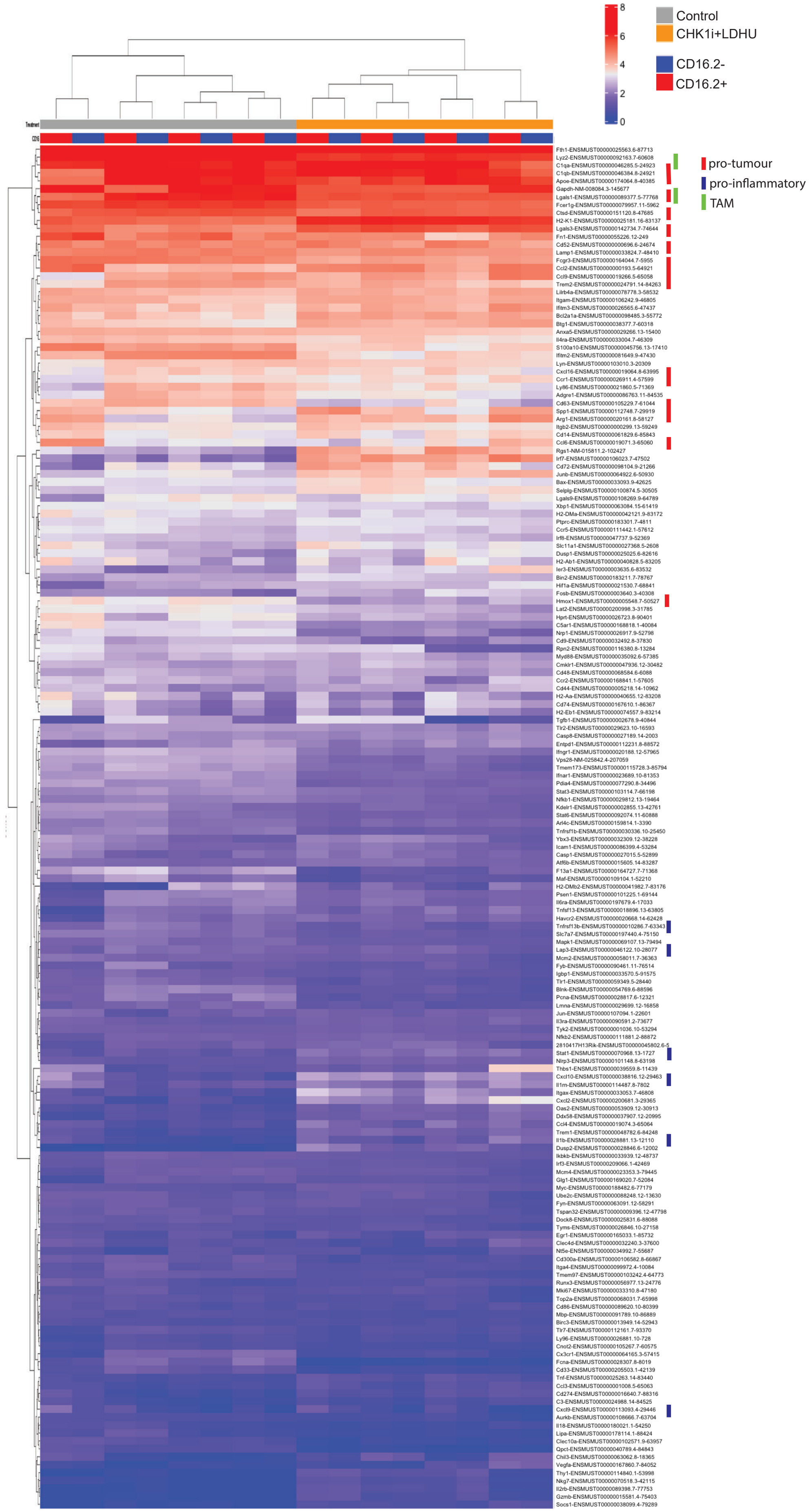


**Supplementary Figure S3:** **H**eat map of unsupervised clustering of 172 genes from scRNAseq of the F4/80⁺: CD11b⁺ population of tumour associated macrophages from untreated YUMMUV1.7 tumour bearing mice. These genes were present at > 2 transcripts per 10,000, The data are ln transcripts per 10,000+1). Cells were also classified on their CD16.2 levels from CITEseq data (see Figure 3B). Defined pro-tumoural and pro-inflammatory and TAM lineage genes are indicated defined by [30].

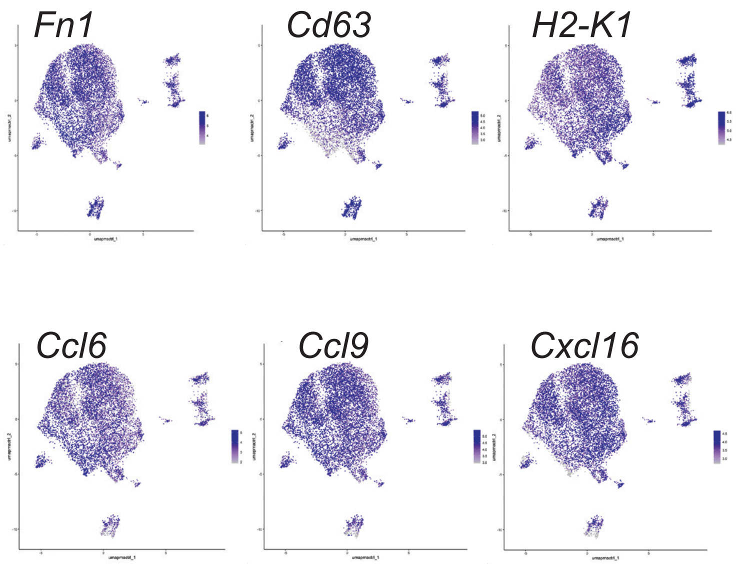


**Supplementary Figure S4:** Marker of the suppressive macrophage phenotype overlaid on the control CD11b$^+$ F4/80$^+$ macrophage UMAP clustering.

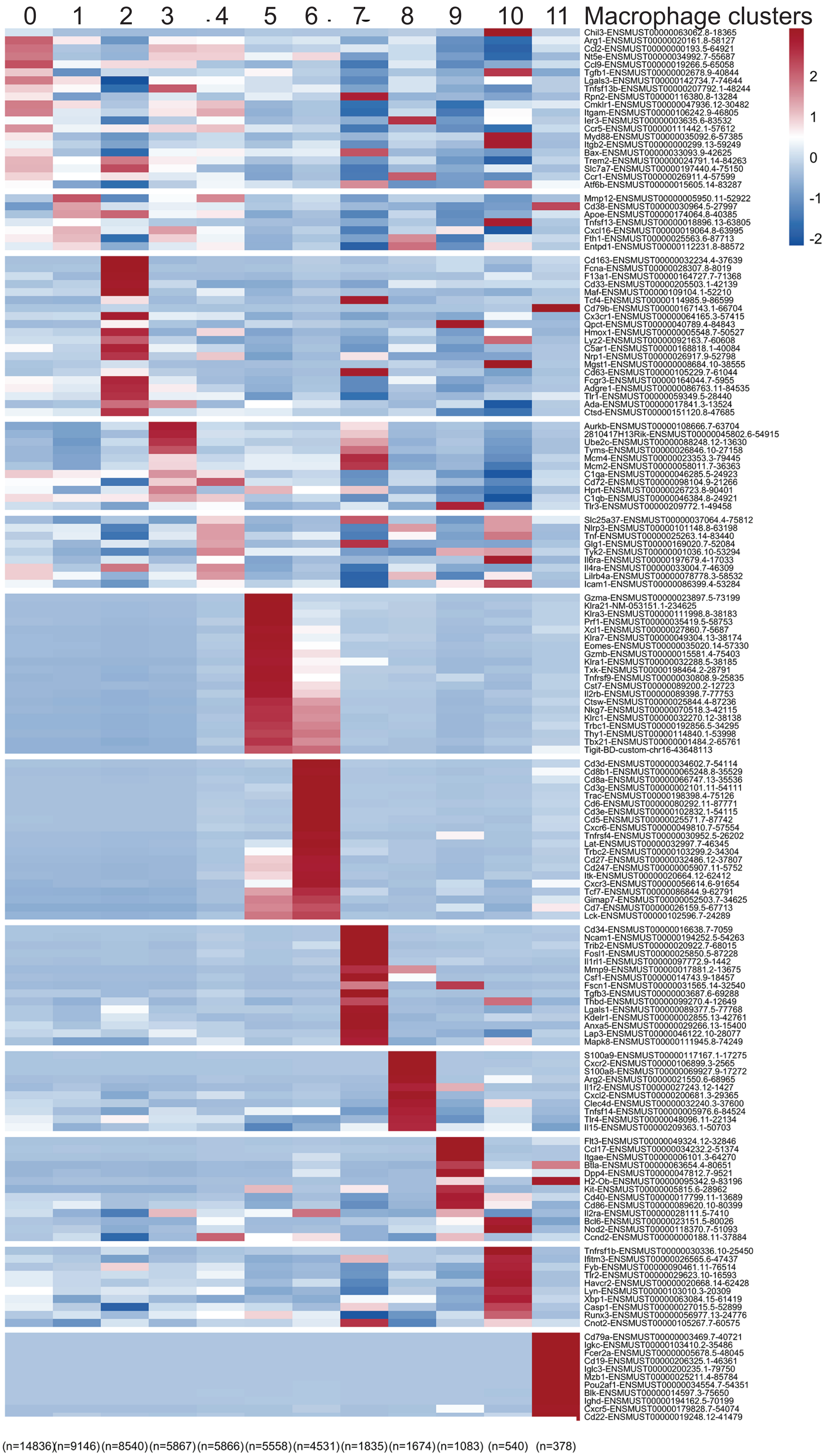


**Supplementary Figure S5:** Heat map of the top genes defining each of the clusters from scRNAseq of the F4/80$^+$: CD11b$^+$ population of tumour associated macrophages from YUMMUV1.7 tumour bearing mice without and with CHK1i+LDHU treatment for 2 weeks. The data are z score of the difference in expression in that cluster compared to all other clusters combined. The number of cells in each cluster is shown (bottom).

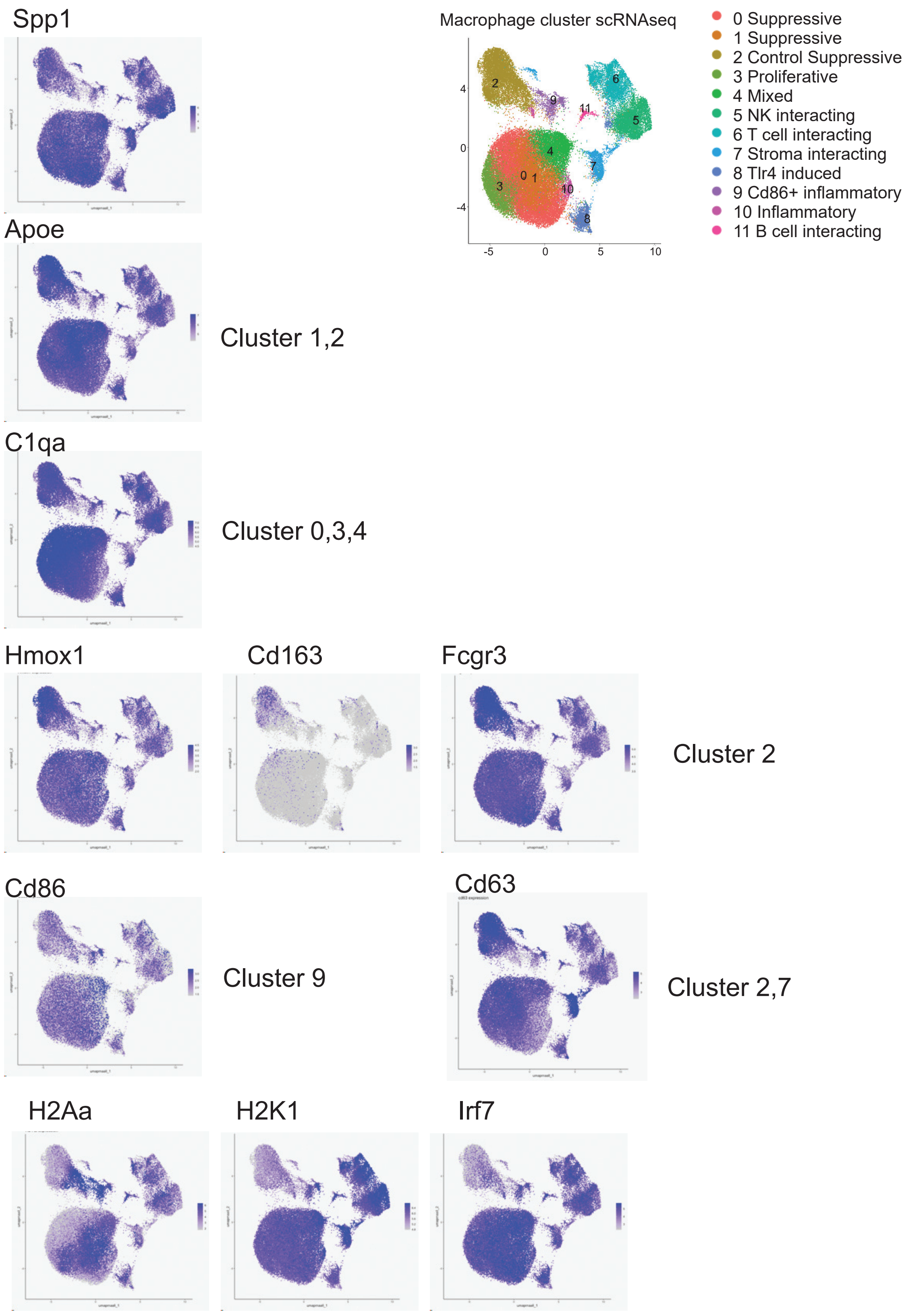


**Supplementary Figure S6:** Cluster defined markers overlaid on the control+treatment CD11b$^{+}$ F4/80$^{+}$ macrophage UMAP clustering.

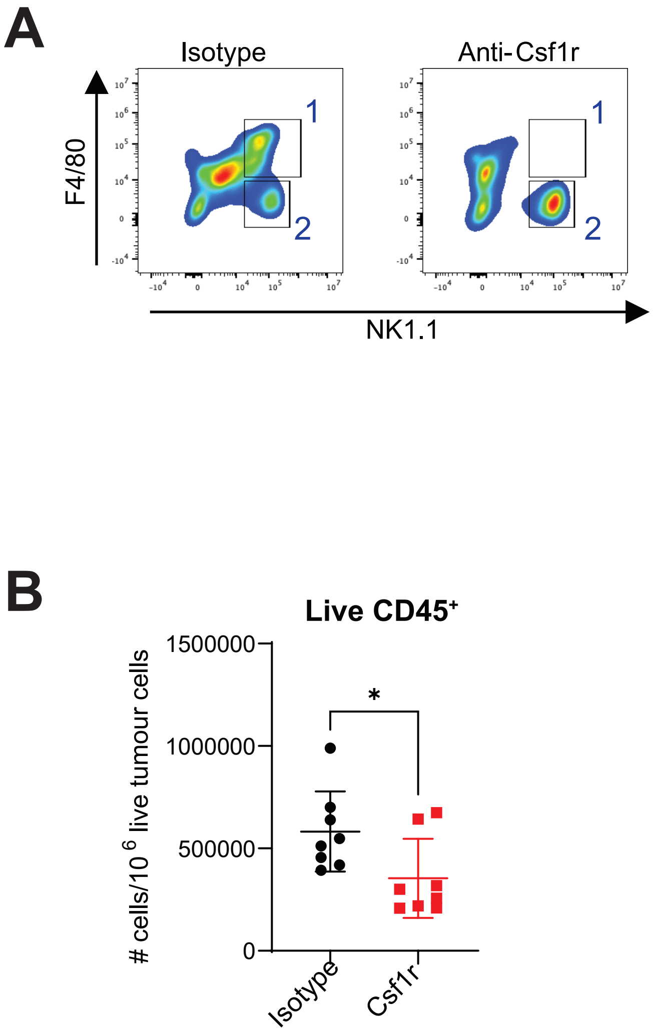


**Supplementary Figure S7: A;** Flow cytometry of live CD45$^+$ cells stained for F4/80 and NK1.1 in YUMMUV1.7 tumours from mice treated with either isotype or CSF1R antibody. Gates 1 and 2 show the Csf1r antibody depleted F4/80$^+$ NK1.1$^+$ CD16.2/FcRγ4$^+$ expressing cells, and gate 2 the F4/80$^-$ NK1.1$^+$ NK cells. **B;** Number of CD45$^+$ cells in YUMMUV1.7 tumours from control and CHK1i+ LDHU treated mice treated with either isotype or CSF1R antibody depletion.
