## Supplementary Table S3 for "Reprogramming tumour-associated macrophages from immune suppressive to inflammatory state by Checkpoint kinase 1 inhibitor combination treatment"

| | Con+Iso | | | | | | | | | TX+Iso | | | | | | | | |
|---|---|---|---|---|---|---|---|---|---|---|---|---|---|---|---|---|---|---|
| | 4-1_CD16+ | 4-1_CD16- | 4-2_CD16+ | 4-2_CD16- | 4-3_CD16+ | 4-3_CD16- | 4-5_CD16+ | 4-5_CD16- | av | 1-1-5_CD16+ | 1-1-5_CD16- | 1-2_CD16+ | 1-2_CD16- | 1-3_CD16+ | 1-3_CD16- | 1-4_CD16+ | 1-4_CD16- | av |
| Gapdh-NM-008084.3-145677 | 6.257145 | 6.309745 | 6.109092 | 6.169282 | 5.148314 | 5.161830 | 6.842799 | 6.832710 | | 6.012769 | 6.004748 | 5.373545 | 5.371568 | 5.442717 | 5.456443 | 5.285297 | 5.275550 | |
| Fth1-ENSMUST00000025563.6-87713 | 6.484415 | 6.410867 | 6.407407 | 6.333685 | 6.282705 | 6.243366 | 6.347776 | 6.356717 | **6.358367** | 6.946734 | 6.870525 | 6.740253 | 6.688931 | 6.792544 | 6.718052 | 6.898679 | 6.798894 | **6.806827** |
| C1qa-ENSMUST00000046285.5-24923 | 6.205419 | 6.038889 | 6.313252 | 5.890254 | 6.408065 | 6.302601 | 5.968282 | 5.988640 | **6.139425** | 5.609296 | 5.038439 | 6.324149 | 6.272661 | 6.294744 | 6.364753 | 6.475132 | 6.208213 | **6.073423** |
| C1qb-ENSMUST00000046384.8-24921 | 6.359377 | 6.189770 | 6.414727 | 5.979683 | 6.430444 | 6.278147 | 4.892030 | 4.847700 | **5.923985** | 5.839320 | 5.257890 | 6.165185 | 6.070790 | 6.344642 | 6.371107 | 6.394827 | 6.107197 | **6.068870** |
| H2-K1-ENSMUST00000025181.16-83137 | 5.130415 | 4.933350 | 5.091456 | 4.923931 | 5.302128 | 5.118339 | 5.467444 | 5.238438 | **5.150688** | 5.893521 | 5.808576 | 6.167089 | 6.086052 | 6.051608 | 5.994920 | 6.050587 | 6.076094 | **6.016056** |
| Lyz2-ENSMUST00000092163.7-60608 | 6.369312 | 6.270994 | 6.523624 | 6.318262 | 6.884418 | 6.898642 | 6.876285 | 6.867586 | **6.626141** | 5.871878 | 5.777538 | 5.954263 | 5.833727 | 5.894803 | 5.786054 | 6.078318 | 5.915296 | **5.888985** |
| Apoe-ENSMUST00000174064.8-40385 | 6.284718 | 6.046968 | 6.302471 | 5.869083 | 6.434336 | 6.311855 | 5.014481 | 5.012249 | **5.909520** | 6.081204 | 5.654074 | 5.880567 | 5.647756 | 5.722613 | 5.416818 | 6.449413 | 6.114712 | **5.870895** |
| Lgals1-ENSMUST00000089377.5-77768 | 6.065059 | 6.179313 | 6.100395 | 6.216267 | 6.103812 | 6.201348 | 5.017378 | 5.122859 | **5.875804** | 5.401147 | 5.412557 | 5.476345 | 5.607056 | 5.486806 | 5.604035 | 5.417841 | 5.484220 | **5.486251** |
| Lgals3-ENSMUST00000142734.7-74644 | 4.864106 | 4.781797 | 4.508609 | 4.524873 | 4.621412 | 4.578693 | 5.198328 | 5.122354 | **4.775022** | 5.607694 | 5.519648 | 5.709213 | 5.578618 | 5.467967 | 5.358654 | 5.394436 | 5.185736 | **5.477746** |
| Fcer1g-ENSMUST00000079957.11-5962 | 5.885855 | 5.793924 | 5.883664 | 5.744010 | 5.866040 | 5.768158 | 5.648647 | 5.601355 | **5.773957** | 5.243964 | 5.102516 | 5.351584 | 5.257059 | 5.376345 | 5.237568 | 5.285189 | 5.152958 | **5.250898** |
| Ctsd-ENSMUST00000151120.8-47685 | 5.517644 | 5.620511 | 5.530966 | 5.509279 | 5.437077 | 5.566543 | 5.127436 | 5.405611 | **5.464384** | 5.268399 | 5.198803 | 5.349917 | 5.373292 | 4.907047 | 4.919582 | 5.306977 | 5.189578 | **5.189199** |
| Ccl2-ENSMUST00000000193.5-64921 | 4.096157 | 4.142886 | 3.975345 | 3.796297 | 3.956438 | 3.814289 | 5.121513 | 5.335400 | **4.279791** | 4.980228 | 4.860425 | 4.826681 | 4.859023 | 4.703428 | 4.783469 | 4.355593 | 4.145438 | **4.689286** |
| Cd52-ENSMUST00000000696.6-24674 | 4.849531 | 4.596353 | 4.790193 | 4.608045 | 4.908877 | 4.648287 | 4.897845 | 4.628518 | **4.740956** | 4.402911 | 4.160338 | 4.504485 | 4.320238 | 4.984544 | 4.764587 | 4.906929 | 4.792224 | **4.604532** |
| Lamp1-ENSMUST00000033824.7-48410 | 4.760503 | 4.861174 | 4.729028 | 4.773067 | 4.852825 | 4.946092 | 4.716184 | 4.839789 | **4.809833** | 4.479745 | 4.425696 | 4.713112 | 4.721945 | 4.435095 | 4.431531 | 4.540693 | 4.465481 | **4.526662** |
| Irf7-ENSMUST00000106023.7-47502 | 2.589388 | 1.719006 | 1.933236 | 1.419509 | 2.368014 | 1.621101 | 2.221496 | 1.655400 | **1.940894** | 4.943833 | 4.605672 | 4.661701 | 4.362826 | 4.439397 | 4.120817 | 4.388669 | 4.035182 | **4.444762** |
| Fn1-ENSMUST00000055226.12-2435 | 5.064608 | 5.386301 | 4.642660 | 4.923747 | 4.617587 | 4.948677 | 5.555256 | 5.812741 | **5.118947** | 4.540890 | 4.724078 | 4.659519 | 4.700171 | 4.662260 | 4.740767 | 3.646572 | 3.745184 | **4.427430** |
| Ccl9-ENSMUST00000019266.5-65058 | 4.417249 | 4.344116 | 4.633224 | 4.509531 | 4.212492 | 4.197163 | 3.092006 | 3.111064 | **4.064606** | 4.984065 | 4.871742 | 4.482425 | 4.452038 | 4.083448 | 4.000850 | 4.205253 | 3.959978 | **4.379975** |
| Bcl2a1a-ENSMUST00000098485.3-55772 | 3.714039 | 3.423740 | 3.757759 | 3.400917 | 4.173149 | 3.796342 | 4.035964 | 3.616400 | **3.739789** | 4.374468 | 4.185421 | 4.411565 | 4.219790 | 4.442979 | 4.249749 | 4.413984 | 4.175769 | **4.309216** |
| Fcgr3-ENSMUST00000164044.7-5955 | 4.858720 | 4.720469 | 4.845665 | 4.654749 | 5.050943 | 4.956981 | 5.128729 | 5.036981 | **4.906655** | 4.446728 | 4.339554 | 4.438641 | 4.352070 | 4.238171 | 4.121919 | 4.303691 | 4.112868 | **4.294205** |
| Ifitm3-ENSMUST00000026565.6-47437 | 4.094845 | 3.661761 | 3.815257 | 3.435154 | 4.029703 | 3.585929 | 4.387796 | 3.989124 | **3.874946** | 4.689039 | 4.531720 | 4.316944 | 4.164121 | 4.137591 | 3.857723 | 4.091039 | 3.887139 | **4.209414** |
| Itgam-ENSMUST00000106242.9-46805 | 3.770359 | 3.732815 | 3.758103 | 3.718793 | 4.194118 | 4.182993 | 4.205315 | 4.224064 | **3.973320** | 4.242713 | 4.244453 | 4.284312 | 4.230836 | 4.263813 | 4.302036 | 4.047458 | 3.965506 | **4.197641** |
| Btg1-ENSMUST00000038377.7-60318 | 3.586703 | 3.501808 | 3.369070 | 3.362038 | 3.695098 | 3.649070 | 3.974293 | 3.838682 | **3.622095** | 4.387644 | 4.412318 | 4.063110 | 3.953960 | 4.286451 | 4.191046 | 4.064777 | 4.158192 | **4.189687** |
| Lilrb4a-ENSMUST00000078778.3-58532 | 4.099795 | 3.965519 | 3.905654 | 3.852258 | 4.079155 | 3.944376 | 4.323145 | 4.206119 | **4.047003** | 4.382274 | 4.311233 | 4.240555 | 4.052662 | 4.271687 | 4.142434 | 4.160181 | 3.934776 | **4.186975** |
| Arg1-ENSMUST00000020161.8-58127 | 3.401045 | 3.422283 | 2.769028 | 2.678697 | 2.834134 | 2.762075 | 4.419869 | 4.337818 | **3.328119** | 4.785060 | 4.617544 | 4.147326 | 3.995251 | 4.095584 | 3.863086 | 4.113653 | 3.608215 | **4.153215** |
| Rgs1-NM-015811.2-102427 | 2.823852 | 2.423878 | 2.310933 | 1.683367 | 2.616477 | 2.316615 | 3.124003 | 2.733696 | **2.504103** | 3.974809 | 3.669654 | 3.881986 | 3.546261 | 4.516787 | 4.263851 | 4.701760 | 4.479851 | **4.129370** |
| Spp1-ENSMUST00000112748.7-29919 | 3.299281 | 3.307811 | 2.911610 | 2.967420 | 3.627289 | 3.581913 | 4.161920 | 4.050701 | **3.488493** | 4.387356 | 4.341776 | 3.862122 | 3.989801 | 4.562681 | 4.775725 | 3.149305 | 3.206594 | **4.034420** |
| Trem2-ENSMUST00000024791.14-84263 | 4.518527 | 4.439996 | 4.481139 | 4.273678 | 4.470653 | 4.412268 | 3.453537 | 3.519356 | **4.196144** | 3.924215 | 3.640230 | 4.324473 | 4.217854 | 4.087072 | 4.006175 | 4.029254 | 3.677170 | **3.988305** |
| Junb-ENSMUST00000064922.6-50930 | 3.360987 | 3.368396 | 2.798480 | 2.748119 | 3.260549 | 3.201645 | 2.720688 | 2.685026 | **3.017986** | 4.222002 | 4.166774 | 3.948609 | 3.762328 | 3.913115 | 3.750231 | 3.891065 | 3.777073 | **3.928900** |
| Anxa5-ENSMUST00000029266.13-15400 | 4.143485 | 4.213733 | 4.076704 | 4.139627 | 4.128156 | 4.177435 | 4.165377 | 4.207982 | **4.156562** | 3.733146 | 3.662049 | 3.928270 | 3.929312 | 3.943750 | 3.944608 | 3.874275 | 3.805202 | **3.852577** |
| Cd72-ENSMUST00000098104.9-21266 | 3.438232 | 2.910574 | 3.173538 | 2.547521 | 3.430442 | 2.917240 | 1.970969 | 1.617378 | **2.750737** | 3.342933 | 2.765057 | 4.054249 | 3.679220 | 4.506229 | 4.319379 | 4.209974 | 3.764119 | **3.830145** |
| Lyn-ENSMUST00000103010.3-20309 | 3.757313 | 3.753212 | 3.897919 | 3.941571 | 3.878128 | 3.897053 | 3.400737 | 3.406356 | **3.741536** | 3.624198 | 3.557365 | 3.695211 | 3.678476 | 3.891696 | 3.802326 | 3.737934 | 3.654201 | **3.705176** |
| S100a10-ENSMUST00000045756.13-17410 | 4.424628 | 4.540681 | 4.313490 | 4.550804 | 4.324145 | 4.461587 | 4.871992 | 4.925139 | **4.551558** | 3.728851 | 3.771700 | 3.768803 | 3.862382 | 3.623080 | 3.655767 | 3.247479 | 3.351205 | **3.626159** |
| Il4ra-ENSMUST00000033004.7-46309 | 3.820171 | 3.863120 | 3.772253 | 3.788027 | 3.921317 | 3.995571 | 3.925518 | 4.053765 | **3.892468** | 3.514083 | 3.420973 | 3.899884 | 3.912683 | 3.544771 | 3.527196 | 3.522931 | 3.394641 | **3.592145** |
| Itgb2-ENSMUST00000000299.13-59249 | 3.448538 | 3.479598 | 3.399540 | 3.360959 | 3.475264 | 3.429376 | 4.055616 | 3.931927 | **3.572602** | 3.532706 | 3.448113 | 3.641393 | 3.483414 | 3.982596 | 3.838090 | 3.417257 | 3.300257 | **3.580478** |
| Ifitm2-ENSMUST00000081649.9-47430 | 4.772755 | 4.772394 | 4.809553 | 4.783579 | 4.711797 | 4.719979 | 4.051208 | 4.016930 | **4.579774** | 3.699737 | 3.528818 | 3.337458 | 3.181948 | 3.818106 | 3.535951 | 3.831728 | 3.669242 | **3.575374** |
| Selplg-ENSMUST00000100874.5-30505 | 3.032052 | 2.907506 | 2.961513 | 2.889592 | 3.082952 | 3.037817 | 2.771425 | 2.718622 | **2.925185** | 3.605396 | 3.540458 | 3.724579 | 3.647768 | 3.583144 | 3.581199 | 3.377660 | 3.444644 | **3.563106** |
| Cxcl16-ENSMUST00000019064.8-63995 | 3.931855 | 3.675304 | 3.797436 | 3.459668 | 3.863223 | 3.634700 | 3.389443 | 3.085962 | **3.604699** | 3.447701 | 3.018708 | 3.660906 | 3.387200 | 3.783973 | 3.552892 | 3.896883 | 3.540192 | **3.536057** |
| Cd14-ENSMUST00000061829.6-85843 | 3.420826 | 3.493850 | 3.169613 | 3.093157 | 3.223455 | 3.209065 | 3.716156 | 3.851601 | **3.397215** | 3.779400 | 3.668678 | 3.216781 | 2.925456 | 3.858613 | 3.689308 | 3.663774 | 3.433405 | **3.529427** |
| Ccr1-ENSMUST00000026911.4-57599 | 3.365182 | 3.396780 | 3.583300 | 3.536184 | 3.677260 | 3.609375 | 3.036193 | 3.037469 | **3.405218** | 3.769009 | 3.636909 | 3.610517 | 3.514078 | 3.405110 | 3.262313 | 3.489782 | 3.266085 | **3.494225** |
| Ly86-ENSMUST00000021860.5-71369 | 4.148301 | 3.898002 | 4.240924 | 3.891403 | 4.220368 | 3.976921 | 2.843485 | 2.507118 | **3.715815** | 3.377396 | 3.015913 | 3.553661 | 3.328903 | 3.664014 | 3.462558 | 3.770831 | 3.410714 | **3.447999** |
| Ccl6-ENSMUST00000019071.3-65060 | 3.296953 | 3.214103 | 3.360428 | 3.047613 | 3.185107 | 3.035140 | 4.696457 | 4.667063 | **3.562858** | 4.192592 | 3.968320 | 3.466820 | 3.151579 | 3.122449 | 2.571227 | 3.615614 | 3.155302 | **3.405488** |
| Bax-ENSMUST00000033093.9-42625 | 3.187406 | 3.170642 | 3.106724 | 3.082073 | 3.061665 | 3.019092 | 3.238791 | 3.232632 | **3.137378** | 3.196677 | 3.065565 | 3.723799 | 3.662096 | 3.565173 | 3.508417 | 3.273799 | 3.136095 | **3.391453** |
| Adgre1-ENSMUST00000086763.11-84535 | 3.868914 | 3.803097 | 3.964812 | 3.768221 | 4.219143 | 4.171838 | 3.061160 | 3.162636 | **3.752478** | 3.186667 | 2.894289 | 3.441140 | 3.431551 | 3.349781 | 3.319118 | 3.342819 | 3.113827 | **3.259899** |
| Slc11a1-ENSMUST00000027368.5-2608 | 2.554675 | 2.388407 | 2.595939 | 2.372080 | 2.715166 | 2.462070 | 3.622845 | 3.374747 | **2.760741** | 3.378524 | 3.198968 | 3.211378 | 2.951276 | 3.719494 | 3.452476 | 3.213822 | 2.944804 | **3.258843** |
| Lgals9-ENSMUST00000108269.9-64789 | 3.543568 | 3.350393 | 3.554068 | 3.444546 | 3.329814 | 3.267629 | 2.100433 | 2.008504 | **3.074869** | 3.448173 | 3.316079 | 3.367466 | 3.292258 | 2.788346 | 2.662733 | 3.223945 | 3.043871 | **3.142859** |
| Ier3-ENSMUST00000003635.6-83532 | 2.224613 | 2.295206 | 2.099362 | 1.983426 | 1.715007 | 1.617053 | 2.730335 | 2.720235 | **2.173155** | 3.782912 | 3.678560 | 3.112118 | 2.919783 | 2.913640 | 2.551191 | 3.148507 | 2.856695 | **3.120426** |
| Ptprc-ENSMUST00000183301.7-4811 | 2.944814 | 2.787005 | 2.799139 | 2.774071 | 3.230884 | 3.175973 | 3.057401 | 2.982536 | **2.968978** | 2.970770 | 2.860306 | 3.141550 | 3.135917 | 3.247742 | 3.231002 | 3.079560 | 3.140971 | **3.100977** |
| Dusp1-ENSMUST00000025025.6-82616 | 2.491938 | 2.431616 | 2.084938 | 1.905034 | 2.955906 | 2.855865 | 3.219112 | 3.198771 | **2.642897** | 3.329947 | 3.185665 | 2.802695 | 2.388867 | 3.275129 | 2.842068 | 3.380071 | 3.106579 | **3.038878** |
| Xbp1-ENSMUST00000063084.15-61419 | 3.206062 | 3.292843 | 3.219783 | 3.289783 | 3.173161 | 3.188760 | 3.242636 | 3.233721 | **3.230843** | 3.114240 | 3.081124 | 3.035329 | 3.028483 | 3.021213 | 2.960505 | 2.782807 | 2.678591 | **2.962786** |
| Ccr5-ENSMUST00000111442.1-57612 | 2.759109 | 2.690738 | 2.674916 | 2.483552 | 3.244052 | 3.194312 | 3.243631 | 3.141367 | **2.928960** | 2.998304 | 2.783116 | 3.180297 | 3.133636 | 3.026125 | 2.985623 | 2.874741 | 2.673832 | **2.956959** |
| H2-Ab1-ENSMUST00000040828.5-83205 | 2.629627 | 2.124313 | 2.873420 | 2.035159 | 3.503324 | 2.824154 | 3.765004 | 2.907106 | **2.832763** | 2.970942 | 2.559587 | 2.892367 | 2.252745 | 3.399577 | 2.520309 | 3.776684 | 3.279683 | **2.956487** |
| Cd63-ENSMUST00000105229.7-61044 | 3.952297 | 3.926038 | 4.121722 | 3.881947 | 4.290414 | 4.285426 | 3.806160 | 3.918244 | **4.022781** | 2.695383 | 2.253047 | 3.004118 | 2.902739 | 2.746448 | 2.594290 | 3.716692 | 3.293124 | **2.900730** |
| H2-DMa-ENSMUST00000042121.9-83172 | 3.148341 | 2.878018 | 3.060855 | 2.781552 | 3.399081 | 3.074114 | 3.611791 | 3.172367 | **3.140765** | 2.846275 | 2.607155 | 3.190000 | 2.865857 | 3.010841 | 2.669006 | 3.073130 | 2.817980 | **2.885030** |
| Bin2-ENSMUST00000183211.7-78767 | 2.542370 | 2.423000 | 2.449317 | 2.375694 | 2.630132 | 2.564788 | 2.258262 | 2.198381 | **2.430243** | 2.698969 | 2.570151 | 3.037956 | 2.946595 | 3.166000 | 3.080050 | 2.744735 | 2.725978 | **2.871304** |
| Hif1a-ENSMUST00000021530.7-68841 | 2.397032 | 2.557489 | 2.537476 | 2.685925 | 2.300593 | 2.448405 | 1.625624 | 1.636715 | **2.273658** | 3.166644 | 3.215479 | 2.768756 | 2.763533 | 2.713868 | 2.670006 | 2.640598 | 2.636052 | **2.821867** |

| | | | | | | | | | | | | | | | | | | |
|---|---|---|---|---|---|---|---|---|---|---|---|---|---|---|---|---|---|---|
| Irf8-ENSMUST00000047737.9-52369 | 2.789529 | 2.688896 | 2.740151 | 2.568056 | 2.795407 | 2.707996 | 3.052741 | 2.932691 | **2.784434** | 2.630662 | 2.461554 | 2.826699 | 2.805877 | 3.027339 | 2.975378 | 2.947973 | 2.872797 | **2.818535** |
| Fosb-ENSMUST00000003640.3-40308 | 2.180885 | 2.151956 | 1.674763 | 1.528909 | 2.244770 | 2.219864 | 2.116151 | 2.216703 | **2.041750** | 2.899668 | 2.740759 | 2.549930 | 2.317199 | 2.791830 | 2.611485 | 3.209279 | 3.049286 | **2.771179** |
| Cmklr1-ENSMUST00000047936.12-30482 | 2.237565 | 2.204941 | 2.372092 | 2.305141 | 2.590103 | 2.544178 | 2.706258 | 2.634168 | **2.449306** | 2.519443 | 2.374227 | 2.921044 | 2.804389 | 2.591155 | 2.469293 | 2.600776 | 2.335582 | **2.576989** |
| Lat2-ENSMUST00000200998.3-31785 | 3.420667 | 3.318810 | 3.179531 | 3.075285 | 3.152796 | 3.088049 | 3.344519 | 3.318771 | **3.237304** | 2.337317 | 2.081396 | 2.702561 | 2.567016 | 2.873379 | 2.686030 | 2.702032 | 2.492894 | **2.555328** |
| Hprt-ENSMUST00000026723.8-90401 | 3.351223 | 3.313237 | 3.383838 | 3.361189 | 2.500012 | 2.335642 | 3.701159 | 3.640875 | **3.198397** | 2.506573 | 2.357216 | 2.799683 | 2.741143 | 2.585028 | 2.464833 | 2.530512 | 2.335504 | **2.540061** |
| Cd74-ENSMUST00000167610.1-86367 | 2.124098 | 1.594922 | 2.372504 | 1.584239 | 2.854325 | 2.199475 | 3.336302 | 2.452716 | **2.314823** | 2.664400 | 2.304922 | 2.454737 | 1.816326 | 2.890932 | 2.051006 | 3.151119 | 2.735621 | **2.508633** |
| Cd44-ENSMUST00000005218.14-10962 | 1.916333 | 2.112052 | 1.943699 | 2.179685 | 2.409819 | 2.510337 | 2.780422 | 2.865142 | **2.339686** | 2.730138 | 2.844723 | 2.394572 | 2.455609 | 2.444306 | 2.503976 | 2.186911 | 2.328629 | **2.486108** |
| Myd88-ENSMUST00000035092.6-57385 | 2.754071 | 2.626099 | 2.854547 | 2.691777 | 2.492758 | 2.348310 | 3.069917 | 2.987424 | **2.728113** | 2.447350 | 2.235701 | 2.808264 | 2.723736 | 2.639325 | 2.464364 | 2.342887 | 2.161277 | **2.477863** |
| Cxcl2-ENSMUST00000200681.3-29365 | 1.099090 | 0.976250 | 0.533811 | 0.521523 | 0.531413 | 0.453413 | 1.265978 | 1.217178 | **0.824832** | 3.384902 | 3.297582 | 2.032050 | 1.738948 | 2.439712 | 2.068286 | 2.510681 | 2.278884 | **2.468881** |
| Cd48-ENSMUST00000068584.6-6088 | 2.449391 | 2.364045 | 2.495427 | 2.371400 | 2.792606 | 2.698027 | 2.393432 | 2.352584 | **2.489614** | 2.486318 | 2.181341 | 2.620744 | 2.472734 | 2.594245 | 2.444236 | 2.439428 | 2.187167 | **2.428277** |
| Ccr2-ENSMUST00000168841.1-57605 | 2.733570 | 2.557796 | 2.397016 | 2.220579 | 2.919832 | 2.639558 | 2.933678 | 2.723277 | **2.640663** | 2.937066 | 2.862126 | 2.706726 | 2.503341 | 2.561638 | 2.211075 | 1.683395 | 1.829895 | **2.411908** |
| Hmox1-ENSMUST00000005548.7-50527 | 3.579674 | 3.513105 | 3.446119 | 3.373597 | 3.224837 | 3.245106 | 3.475319 | 3.660186 | **3.439743** | 2.777452 | 2.584939 | 2.382638 | 2.280549 | 2.261780 | 2.063712 | 2.520664 | 2.190373 | **2.382764** |
| H2-Aa-ENSMUST00000040655.12-83208 | 2.477927 | 1.824540 | 2.741614 | 1.748981 | 3.416354 | 2.694449 | 3.630566 | 2.746766 | **2.660150** | 2.232098 | 1.836251 | 2.090890 | 1.479852 | 2.843402 | 1.900728 | 3.202991 | 2.671337 | **2.282194** |
| Cxcl10-ENSMUST00000038816.12-29463 | 0.982550 | 0.759398 | 0.975936 | 0.607818 | 1.224258 | 0.832772 | 2.373431 | 1.761951 | **1.189764** | 2.426708 | 1.998629 | 2.367380 | 2.030831 | 2.542492 | 2.252855 | 2.487611 | 2.124238 | **2.278843** |
| Itgax-ENSMUST00000033053.7-46808 | 1.114397 | 0.715661 | 0.939923 | 0.646879 | 1.176230 | 0.824014 | 1.310142 | 0.842187 | **0.946179** | 1.927306 | 1.633400 | 2.258181 | 1.878231 | 2.959651 | 2.603710 | 2.445222 | 2.169365 | **2.234383** |
| H2-Eb1-ENSMUST00000074557.9-83214 | 2.008289 | 1.516028 | 2.237627 | 1.433404 | 2.856066 | 2.165008 | 3.202765 | 2.330988 | **2.218772** | 2.147235 | 1.802548 | 2.130773 | 1.513019 | 2.649874 | 1.778530 | 2.978845 | 2.454358 | **2.181898** |
| Casp8-ENSMUST00000027189.14-2003 | 2.163041 | 2.120733 | 2.252782 | 2.105117 | 2.445285 | 2.364374 | 2.338150 | 2.261170 | **2.256331** | 2.267977 | 2.167969 | 2.238151 | 2.128353 | 2.153631 | 2.019371 | 2.214022 | 2.059591 | **2.156133** |
| Tlr2-ENSMUST00000029623.10-16593 | 2.345210 | 2.363617 | 2.416404 | 2.353168 | 2.397717 | 2.376976 | 2.275775 | 2.349105 | **2.359746** | 2.292715 | 2.054805 | 2.192677 | 2.053434 | 2.345159 | 2.173264 | 2.152956 | 1.894297 | **2.144913** |
| Rpn2-ENSMUST00000116380.8-13284 | 2.859317 | 2.891173 | 3.044661 | 3.026866 | 2.557752 | 2.569606 | 3.216568 | 3.173817 | **2.917470** | 1.280682 | 1.191697 | 3.010023 | 3.129226 | 2.878873 | 2.928551 | 1.354990 | 1.264901 | **2.129868** |
| Entpd1-ENSMUST00000112231.8-88572 | 2.196995 | 2.123837 | 2.495991 | 2.295518 | 1.919920 | 1.850595 | 1.534163 | 1.408412 | **1.978179** | 2.365460 | 2.162774 | 2.068598 | 1.933276 | 2.076634 | 1.780825 | 2.327864 | 2.014170 | **2.091200** |
| C5ar1-ENSMUST00000168818.1-40084 | 2.936763 | 3.115127 | 2.899601 | 2.928184 | 3.007356 | 3.009945 | 3.503449 | 3.622600 | **3.127878** | 2.118280 | 2.012722 | 2.113089 | 2.042569 | 2.051825 | 1.987898 | 2.204875 | 2.032884 | **2.070518** |
| Il1rn-ENSMUST00000114487.8-7802 | 1.430714 | 1.178568 | 1.500975 | 1.325280 | 1.104718 | 0.906867 | 2.127128 | 1.929651 | **1.437988** | 2.378901 | 2.117060 | 1.785416 | 1.514298 | 2.145748 | 1.895122 | 2.158858 | 1.793439 | **1.973605** |
| Tgfb1-ENSMUST00000002678.9-40844 | 2.366240 | 2.347269 | 1.672872 | 1.541983 | 3.011352 | 3.018911 | 0.854085 | 0.826557 | **1.954908** | 1.079980 | 1.028743 | 3.073952 | 3.072470 | 3.167182 | 3.149721 | 0.550500 | 0.500455 | **1.952875** |
| Cd9-ENSMUST00000032492.8-37830 | 2.749832 | 2.747231 | 2.639322 | 2.675014 | 2.299485 | 2.431128 | 3.018190 | 3.051733 | **2.701492** | 2.272874 | 2.159354 | 1.618032 | 1.447615 | 2.356213 | 2.229045 | 1.778405 | 1.648453 | **1.938749** |
| Thbs1-ENSMUST00000039559.8-11439 | 1.180391 | 1.082854 | 1.028986 | 1.103526 | 0.771826 | 0.854787 | 2.174728 | 2.156669 | **1.294221** | 3.607375 | 3.625246 | 1.462018 | 1.277692 | 1.443461 | 1.189015 | 1.449479 | 1.379827 | **1.929264** |
| Nrp1-ENSMUST00000026917.9-52798 | 3.033956 | 3.132850 | 3.189333 | 3.150015 | 2.929240 | 3.013550 | 2.863557 | 3.112179 | **3.053085** | 1.646938 | 1.439385 | 2.121699 | 2.091219 | 1.821120 | 1.820642 | 2.349753 | 2.130001 | **1.927595** |
| Ifngr1-ENSMUST00000020188.12-57965 | 2.478974 | 2.479143 | 2.540566 | 2.513680 | 2.809520 | 2.808390 | 2.601795 | 2.560302 | **2.599046** | 1.878785 | 1.739112 | 1.959786 | 1.807487 | 1.922146 | 1.751822 | 1.909777 | 1.905602 | **1.859315** |
| Ybx3-ENSMUST00000032309.12-38228 | 1.496405 | 1.543187 | 1.330198 | 1.310997 | 2.021711 | 1.902407 | 2.446370 | 2.255313 | **1.788323** | 1.325844 | 1.221849 | 2.196848 | 2.125721 | 2.062171 | 1.981933 | 1.970097 | 1.840752 | **1.840652** |
| Stat1-ENSMUST00000070968.13-1727 | 1.267800 | 0.971427 | 1.317299 | 1.024428 | 1.467170 | 1.178766 | 1.221663 | 0.917576 | **1.170766** | 1.933833 | 1.690385 | 1.823715 | 1.685025 | 1.908583 | 1.694518 | 1.964552 | 1.767288 | **1.808487** |
| Casp1-ENSMUST00000027015.5-52899 | 1.506196 | 1.381069 | 1.863232 | 1.707621 | 1.465278 | 1.335431 | 2.267117 | 2.005406 | **1.691419** | 1.695695 | 1.519585 | 1.983058 | 1.776837 | 2.008636 | 1.775969 | 1.796627 | 1.556803 | **1.764151** |
| Atf6b-ENSMUST00000015605.14-83287 | 1.537643 | 1.588742 | 1.568214 | 1.634212 | 1.722261 | 1.707585 | 1.835565 | 1.796592 | **1.673852** | 1.653016 | 1.519541 | 1.854312 | 1.887776 | 1.873786 | 1.831946 | 1.693812 | 1.581569 | **1.736970** |
| Stat3-ENSMUST00000103114.7-66198 | 2.125337 | 2.234374 | 2.275312 | 2.195705 | 2.274756 | 2.374612 | 2.082665 | 2.098565 | **2.207666** | 1.585551 | 1.524110 | 1.779995 | 1.783393 | 1.897467 | 1.876699 | 1.656247 | 1.695019 | **1.724810** |
| Icam1-ENSMUST00000086399.4-53284 | 1.604512 | 1.641143 | 1.658360 | 1.580464 | 1.926570 | 1.862523 | 2.314384 | 2.157217 | **1.843147** | 1.479544 | 1.280971 | 1.982606 | 1.877313 | 1.983024 | 1.871743 | 1.730821 | 1.583237 | **1.723657** |
| Tnfsf13-ENSMUST00000018896.13-63805 | 1.757811 | 1.707683 | 2.041941 | 1.903009 | 1.895495 | 1.824626 | 0.566644 | 0.560030 | **1.532155** | 1.981254 | 1.689986 | 1.747652 | 1.526687 | 1.600407 | 1.267883 | 1.950637 | 1.556822 | **1.665166** |
| Psen1-ENSMUST00000101225.1-69144 | 2.055076 | 2.115312 | 2.075496 | 2.169759 | 2.150115 | 2.154836 | 1.003920 | 0.965595 | **1.836264** | 1.655837 | 1.610860 | 1.828077 | 1.811544 | 1.668907 | 1.613581 | 1.611984 | 1.502970 | **1.662970** |
| Havcr2-ENSMUST00000020668.14-62428 | 1.713857 | 1.497657 | 1.821790 | 1.721136 | 1.957236 | 1.848896 | 0.893512 | 0.754163 | **1.526031** | 1.782641 | 1.563362 | 1.689862 | 1.537523 | 1.851139 | 1.615919 | 1.757515 | 1.464333 | **1.657787** |
| Ifnar1-ENSMUST00000023689.10-81353 | 2.418387 | 2.419386 | 2.168007 | 2.262084 | 2.477401 | 2.483235 | 2.258951 | 2.289575 | **2.347128** | 1.610350 | 1.490359 | 1.723421 | 1.753809 | 1.647990 | 1.627310 | 1.717533 | 1.602658 | **1.646679** |
| Vps28-NM-025842.4-207059 | 2.548948 | 2.418115 | 2.477076 | 2.431849 | 2.605809 | 2.565032 | 2.434473 | 2.389731 | **2.483879** | 1.484435 | 1.331388 | 1.687977 | 1.545396 | 1.841220 | 1.633608 | 1.910472 | 1.705354 | **1.642481** |
| Il6ra-ENSMUST00000197679.4-17033 | 1.940637 | 1.913075 | 2.114635 | 2.133529 | 2.147032 | 2.160577 | 1.140753 | 1.113397 | **1.832954** | 1.660290 | 1.593609 | 1.899351 | 1.838894 | 1.647288 | 1.604445 | 1.485521 | 1.398539 | **1.640992** |
| Nfkb1-ENSMUST00000029812.13-19464 | 1.985891 | 2.005961 | 2.117656 | 2.113450 | 2.076191 | 2.101485 | 1.933714 | 1.927698 | **2.032756** | 1.411635 | 1.383533 | 1.731183 | 1.759965 | 1.805440 | 1.787123 | 1.557106 | 1.624555 | **1.632568** |
| Arl4c-ENSMUST00000159814.1-3390 | 1.789601 | 1.822766 | 1.855758 | 1.860465 | 2.087516 | 2.035417 | 1.929593 | 1.917457 | **1.912321** | 1.689326 | 1.539950 | 1.759329 | 1.619859 | 1.566740 | 1.416899 | 1.753638 | 1.592915 | **1.617332** |
| Trem1-ENSMUST00000048782.6-84248 | 0.801169 | 0.676435 | 0.672049 | 0.568288 | 0.554024 | 0.453906 | 1.200555 | 1.039600 | **0.745753** | 1.745801 | 1.643214 | 1.749528 | 1.547044 | 1.741103 | 1.614183 | 1.518533 | 1.331797 | **1.611400** |
| Tmem173-ENSMUST00000115728.3-85794 | 2.463581 | 2.265795 | 2.435680 | 2.302826 | 2.751378 | 2.658952 | 2.398516 | 2.173365 | **2.431262** | 1.306467 | 1.183197 | 1.732509 | 1.650223 | 1.817694 | 1.724067 | 1.817856 | 1.645385 | **1.609675** |
| Ccl4-ENSMUST00000019074.3-65064 | 0.972069 | 0.855828 | 1.033545 | 0.656340 | 0.780667 | 0.627447 | 1.344221 | 1.331002 | **0.950140** | 1.838917 | 1.402480 | 1.404258 | 1.080912 | 2.005923 | 1.616908 | 1.952413 | 1.544194 | **1.605751** |
| Oas2-ENSMUST00000053909.12-30913 | 1.001616 | 0.690190 | 0.807129 | 0.650956 | 1.028759 | 0.804920 | 0.854196 | 0.727051 | **0.820602** | 1.953769 | 1.767615 | 1.691131 | 1.548079 | 1.501649 | 1.296615 | 1.677150 | 1.409780 | **1.605723** |
| Nlrp3-ENSMUST00000101148.8-63198 | 1.113877 | 1.182247 | 1.129303 | 1.097424 | 1.159766 | 1.094731 | 1.368586 | 1.339240 | **1.185647** | 1.653959 | 1.577021 | 1.671752 | 1.461352 | 1.718668 | 1.516485 | 1.692957 | 1.485840 | **1.597249** |
| Pdia4-ENSMUST00000077290.8-34496 | 2.164455 | 2.327262 | 2.156104 | 2.209852 | 2.117367 | 2.161253 | 2.438668 | 2.490868 | **2.258229** | 1.677781 | 1.651028 | 1.598753 | 1.700710 | 1.430737 | 1.501562 | 1.564415 | 1.492858 | **1.577231** |
| Kdelr1-ENSMUST00000002855.13-42761 | 1.934173 | 1.909319 | 1.672072 | 1.687329 | 2.324621 | 2.342164 | 2.283228 | 2.312352 | **2.058157** | 1.467075 | 1.321164 | 1.769026 | 1.748798 | 1.547704 | 1.496255 | 1.643202 | 1.553072 | **1.568287** |
| 2810417H13Rik-ENSMUST00000045802.6-54915 | 1.440979 | 1.310157 | 1.659074 | 1.451711 | 1.584988 | 1.450347 | 0.947493 | 0.890065 | **1.341852** | 1.145389 | 0.984937 | 1.697167 | 1.815159 | 1.679837 | 1.735574 | 1.825475 | 1.643845 | **1.565923** |
| Dusp2-ENSMUST00000028846.6-12002 | 0.379602 | 0.329115 | 0.298989 | 0.338771 | 0.409708 | 0.351930 | 0.253412 | 0.197975 | **0.319938** | 1.278092 | 1.059588 | 1.424954 | 1.294597 | 2.192395 | 1.942397 | 1.587194 | 1.708052 | **1.560909** |
| Nfkb2-ENSMUST00000111881.2-88872 | 1.528081 | 1.581326 | 1.467788 | 1.432596 | 1.614635 | 1.594013 | 1.172764 | 1.145327 | **1.442066** | 1.499642 | 1.420271 | 1.708114 | 1.683074 | 1.612503 | 1.569546 | 1.504638 | 1.424191 | **1.552747** |
| Ddx58-ENSMUST00000037907.12-20995 | 0.972598 | 0.742577 | 0.902138 | 0.772901 | 1.001473 | 0.863760 | 0.580549 | 0.509110 | **0.793138** | 1.834797 | 1.696604 | 1.611160 | 1.612116 | 1.409457 | 1.339263 | 1.508124 | 1.372762 | **1.548035** |
| Il3ra-ENSMUST00000090591.2-73677 | 1.364030 | 1.247073 | 1.280587 | 1.302468 | 1.418492 | 1.316232 | 1.640230 | 1.597383 | **1.395812** | 1.726774 | 1.525803 | 1.612508 | 1.465613 | 1.633958 | 1.471558 | 1.505662 | 1.394549 | **1.542053** |
| Jun-ENSMUST00000107094.1-22601 | 1.557458 | 1.380225 | 1.693031 | 1.443863 | 1.019181 | 0.900783 | 1.859498 | 1.776906 | **1.453868** | 1.573918 | 1.480091 | 1.490905 | 1.297141 | 1.664730 | 1.472818 | 1.604485 | 1.441587 | **1.503209** |
| Stat6-ENSMUST00000092074.11-60888 | 1.829604 | 1.892491 | 1.763542 | 1.721626 | 2.093595 | 2.097816 | 2.187770 | 2.226716 | **1.976645** | 1.476348 | 1.318770 | 1.554902 | 1.489744 | 1.524647 | 1.400170 | 1.609234 | 1.453188 | **1.478375** |
| Lap3-ENSMUST00000046122.10-28077 | 1.798745 | 1.670086 | 1.747243 | 1.599428 | 1.960896 | 1.733418 | 1.553303 | 1.325993 | **1.673639** | 1.233320 | 1.099422 | 1.687942 | 1.538679 | 1.609769 | 1.389287 | 1.775167 | 1.468482 | **1.475258** |
| H2-DMb2-ENSMUST00000041982.7-83176 | 2.757045 | 2.363142 | 2.687543 | 2.290958 | 0.586251 | 0.487344 | 0.897003 | 0.719079 | **1.598546** | 2.004416 | 1.737164 | 1.869634 | 1.485243 | 1.626372 | 1.234136 | 1.035902 | 0.794936 | **1.473475** |

| | | | | | | | | | | | | | | | | | | |
|---|---|---|---|---|---|---|---|---|---|---|---|---|---|---|---|---|---|---|
| Tyk2-ENSMUST00000001036.10-53294 | 1.439448 | 1.380545 | 1.386203 | 1.409623 | 1.544242 | 1.449712 | 1.582581 | 1.507494 | **1.462481** | 1.363365 | 1.207779 | 1.577449 | 1.417333 | 1.556486 | 1.484859 | 1.465949 | 1.371798 | **1.430627** |
| Il1b-ENSMUST00000028881.13-12110 | 0.474182 | 0.383952 | 0.600304 | 0.426054 | 0.689541 | 0.509627 | 1.338252 | 1.005558 | **0.678434** | 2.033747 | 1.703231 | 1.135452 | 0.771609 | 1.720183 | 1.120776 | 1.433921 | 1.117069 | **1.379499** |
| Tnfrsf1b-ENSMUST00000030336.10-25450 | 1.832396 | 1.802974 | 1.888155 | 1.965784 | 2.128206 | 2.233413 | 1.977042 | 2.066766 | **1.986842** | 1.409628 | 1.323132 | 1.373449 | 1.279509 | 1.410184 | 1.275481 | 1.410909 | 1.385894 | **1.358523** |
| Myc-ENSMUST00000188482.6-77179 | 1.068185 | 1.051700 | 0.985110 | 0.950992 | 1.174413 | 1.086497 | 1.274889 | 1.208735 | **1.100065** | 1.157901 | 1.051835 | 1.427318 | 1.408521 | 1.457895 | 1.411582 | 1.336735 | 1.280235 | **1.316503** |
| Mapk1-ENSMUST00000069107.13-79494 | 1.563168 | 1.604423 | 1.722705 | 1.769866 | 1.880469 | 1.874393 | 1.609929 | 1.607179 | **1.704016** | 1.263326 | 1.194607 | 1.344529 | 1.355783 | 1.335537 | 1.325982 | 1.311351 | 1.247540 | **1.297332** |
| Mcm2-ENSMUST00000058011.7-36363 | 1.668900 | 1.685978 | 1.629899 | 1.509168 | 1.666280 | 1.537268 | 1.886491 | 1.806040 | **1.673753** | 0.991639 | 0.827533 | 1.549356 | 1.481286 | 1.403783 | 1.373985 | 1.393309 | 1.318311 | **1.292400** |
| Tnfrsf13b-ENSMUST00000010286.7-63343 | 1.879591 | 1.657029 | 1.838126 | 1.665632 | 2.091529 | 1.990673 | 1.632189 | 1.572205 | **1.790872** | 1.358482 | 1.208460 | 1.630545 | 1.503329 | 1.246352 | 1.146138 | 1.166718 | 1.014465 | **1.284311** |
| Slc7a7-ENSMUST00000197440.4-75150 | 1.803061 | 1.691018 | 1.834417 | 1.578910 | 1.962570 | 1.899463 | 1.945438 | 1.937521 | **1.831550** | 1.236129 | 1.089071 | 1.411815 | 1.336133 | 1.332203 | 1.198403 | 1.385501 | 1.148242 | **1.267187** |
| Ikbkb-ENSMUST00000033939.12-48737 | 1.437478 | 1.380004 | 1.399227 | 1.439359 | 1.638474 | 1.607185 | 1.200081 | 1.215830 | **1.414705** | 1.223846 | 1.136797 | 1.276917 | 1.236488 | 1.372728 | 1.306129 | 1.221132 | 1.141222 | **1.239407** |
| F13a1-ENSMUST00000164727.7-71368 | 2.033771 | 2.091439 | 2.495152 | 2.581951 | 2.958221 | 3.051473 | 2.301779 | 2.631960 | **2.518218** | 1.675078 | 1.599767 | 1.365150 | 1.321864 | 0.917740 | 0.763527 | 1.148681 | 1.069176 | **1.232623** |
| Mcm4-ENSMUST00000023353.3-79445 | 1.514536 | 1.445586 | 1.607456 | 1.464896 | 1.503568 | 1.400585 | 0.919862 | 0.834575 | **1.336383** | 0.957370 | 0.859736 | 1.358672 | 1.349153 | 1.276739 | 1.229814 | 1.345478 | 1.281764 | **1.207341** |
| Irf3-ENSMUST0000209066.1-42469 | 1.356649 | 1.394405 | 1.351006 | 1.286994 | 1.464177 | 1.425219 | 1.168344 | 1.115670 | **1.320308** | 1.071537 | 0.932325 | 1.335535 | 1.305012 | 1.288801 | 1.207074 | 1.250887 | 1.110259 | **1.187679** |
| Fyb-ENSMUST00000090461.11-76514 | 1.253241 | 1.180066 | 1.448652 | 1.386360 | 2.160583 | 2.150647 | 1.267804 | 1.263866 | **1.513902** | 1.033783 | 0.897372 | 1.214092 | 1.113170 | 1.205104 | 1.077451 | 1.523401 | 1.277781 | **1.167770** |
| Ube2c-ENSMUST00000088248.12-13630 | 1.137334 | 1.091354 | 1.288998 | 1.077316 | 1.487167 | 1.458201 | 1.455438 | 1.462093 | **1.307238** | 0.767311 | 0.659551 | 1.292759 | 1.459891 | 1.289389 | 1.400888 | 1.276273 | 1.138329 | **1.160549** |
| Igbp1-ENSMUST00000033570.5-91575 | 1.633747 | 1.406095 | 1.449082 | 1.230768 | 1.824465 | 1.580625 | 1.508556 | 1.370168 | **1.500438** | 1.098066 | 0.902835 | 1.342386 | 1.198587 | 1.220221 | 1.113081 | 1.242287 | 1.099625 | **1.152136** |
| Egr1-ENSMUST00000165033.1-85732 | 1.260562 | 1.250213 | 0.839057 | 0.647345 | 0.746463 | 0.733433 | 1.264738 | 1.189051 | **0.991358** | 1.314150 | 1.201059 | 0.822123 | 0.685130 | 1.353001 | 1.132079 | 1.388974 | 1.253074 | **1.143699** |
| Fyn-ENSMUST00000063091.12-58291 | 1.201610 | 1.212639 | 1.285370 | 1.230178 | 1.492595 | 1.488755 | 1.426445 | 1.338584 | **1.334522** | 0.786708 | 0.741223 | 1.202132 | 1.141038 | 1.396578 | 1.384901 | 1.130755 | 1.137322 | **1.115082** |
| Blnk-ENSMUST00000054769.6-88596 | 2.174968 | 2.051815 | 2.021712 | 1.894377 | 2.060738 | 1.994431 | 1.440836 | 1.525389 | **1.895533** | 0.989987 | 0.761660 | 1.174769 | 1.152592 | 1.092545 | 1.126899 | 1.398052 | 1.070638 | **1.095893** |
| Glg1-ENSMUST00000169020.7-52084 | 1.315924 | 1.363383 | 1.296024 | 1.431943 | 1.442611 | 1.520191 | 0.560147 | 0.557184 | **1.185926** | 1.036975 | 0.934543 | 1.142211 | 1.196457 | 1.064229 | 1.084246 | 1.117693 | 1.016688 | **1.074130** |
| Lmna-ENSMUST00000029699.12-16858 | 1.476105 | 1.746945 | 1.612950 | 1.866088 | 1.581071 | 1.759491 | 1.238681 | 1.458982 | **1.592539** | 0.988315 | 1.030158 | 1.061417 | 1.270575 | 1.096005 | 1.279206 | 0.900860 | 0.946813 | **1.071669** |
| Tspan32-ENSMUST00000009396.12-47798 | 1.397592 | 1.179073 | 1.384740 | 1.093258 | 1.510158 | 1.341891 | 1.335028 | 1.230649 | **1.309049** | 0.927127 | 0.737410 | 1.173067 | 1.107646 | 1.207064 | 1.083131 | 1.251860 | 1.050344 | **1.067206** |
| Maf-ENSMUST00000109104.1-52210 | 1.538877 | 1.580290 | 1.915685 | 1.839194 | 2.480590 | 2.547758 | 1.805072 | 1.998311 | **1.963222** | 1.105233 | 1.021818 | 1.164453 | 1.171971 | 0.813685 | 0.752642 | 1.329529 | 1.146114 | **1.063181** |
| Tlr1-ENSMUST00000059349.5-28440 | 1.618383 | 1.462308 | 1.654790 | 1.458961 | 1.925825 | 1.802684 | 1.516477 | 1.423677 | **1.607888** | 1.004104 | 0.809926 | 1.323322 | 1.184719 | 1.002266 | 0.863758 | 1.124166 | 0.909359 | **1.027703** |
| Chil3-ENSMUST00000063062.8-18365 | 0.273119 | 0.252654 | 0.281376 | 0.344343 | 0.403396 | 0.400506 | 1.309494 | 1.147389 | **0.551535** | 1.513179 | 1.520629 | 1.443843 | 1.230617 | 0.846538 | 0.634453 | 0.369181 | 0.467418 | **1.003232** |
| Pcna-ENSMUST00000028817.6-12321 | 1.650395 | 1.594268 | 2.186636 | 2.065829 | 2.105576 | 2.010237 | 1.584255 | 1.489069 | **1.835783** | 0.946604 | 0.807791 | 1.126819 | 1.028756 | 1.050332 | 0.908611 | 1.154013 | 0.958442 | **0.997671** |
| Fcna-ENSMUST00000028307.8-8019 | 1.302758 | 1.305916 | 1.911451 | 1.699164 | 1.730863 | 1.770699 | 0.809979 | 0.966495 | **1.437166** | 0.368780 | 0.254103 | 0.293293 | 0.298781 | 0.213644 | 0.177290 | 0.361798 | 0.249263 | **0.277119** |
| Cx3cr1-ENSMUST00000064165.3-57415 | 1.651371 | 1.561834 | 1.825923 | 1.788655 | 1.677648 | 1.604153 | 0.549540 | 0.504209 | **1.395417** | 0.471621 | 0.390395 | 0.757096 | 0.746911 | 0.596346 | 0.606749 | 0.893331 | 0.736986 | **0.649929** |
| Cd300a-ENSMUST00000106582.8-66867 | 1.202089 | 1.075552 | 1.446711 | 1.232259 | 1.564810 | 1.479421 | 1.390471 | 1.276527 | **1.333480** | 0.927278 | 0.776484 | 0.918264 | 0.784552 | 0.790379 | 0.549513 | 0.959251 | 0.782728 | **0.811056** |
| Tmem97-ENSMUST00000103242.4-64773 | 1.337090 | 1.231805 | 1.244562 | 1.190822 | 1.467757 | 1.414653 | 1.095843 | 1.030472 | **1.251626** | 0.694457 | 0.624879 | 0.985865 | 1.004643 | 0.885421 | 0.893259 | 0.806033 | 0.729508 | **0.828008** |
| Dock8-ENSMUST00000025831.6-88088 | 1.356647 | 1.306741 | 1.069604 | 1.049566 | 1.222690 | 1.230728 | 1.362651 | 1.336554 | **1.241898** | 0.962438 | 0.856495 | 1.073141 | 1.084723 | 1.018748 | 0.990329 | 0.893328 | 0.899285 | **0.972311** |
| Itga4-ENSMUST00000099972.4-10084 | 0.934918 | 1.004058 | 1.291274 | 1.309501 | 1.626100 | 1.620281 | 1.064245 | 0.948602 | **1.224872** | 0.820666 | 0.780504 | 0.864904 | 0.864417 | 0.908664 | 0.862358 | 0.904662 | 0.926554 | **0.866591** |
| Cd33-ENSMUST00000205503.1-42139 | 1.160403 | 1.015123 | 1.350019 | 1.181153 | 1.616306 | 1.477876 | 0.815403 | 0.828554 | **1.180605** | 0.519568 | 0.432741 | 0.644176 | 0.554296 | 0.397838 | 0.327780 | 0.634764 | 0.500865 | **0.501503** |
| Tyms-ENSMUST00000026846.10-27158 | 1.162983 | 1.077736 | 1.085999 | 0.944996 | 1.139376 | 1.069102 | 1.145062 | 1.089411 | **1.089333** | 0.736398 | 0.636840 | 1.223862 | 1.200848 | 1.027160 | 0.991089 | 1.046945 | 0.903960 | **0.970888** |
| Clec4d-ENSMUST00000032240.3-37600 | 1.149591 | 0.939385 | 1.095964 | 0.962153 | 0.849035 | 0.852566 | 1.468853 | 1.343740 | **1.082661** | 1.707630 | 1.518071 | 0.761352 | 0.600046 | 0.900746 | 0.652166 | 1.019831 | 0.781576 | **0.992677** |
| Top2a-ENSMUST00000068031.7-65998 | 1.078469 | 0.889867 | 1.046256 | 0.801417 | 1.189226 | 1.093279 | 1.133710 | 1.034138 | **1.033295** | 0.551911 | 0.425070 | 0.849685 | 0.845525 | 0.816067 | 0.792525 | 0.939110 | 0.856766 | **0.759582** |
| Runx3-ENSMUST00000056977.13-24776 | 0.885476 | 0.725437 | 0.658311 | 0.706810 | 1.228753 | 1.161828 | 1.506460 | 1.322202 | **1.024409** | 0.884918 | 0.798051 | 0.901120 | 0.848038 | 0.915063 | 0.842402 | 1.172501 | 1.077571 | **0.929958** |
| Tlr7-ENSMUST00000112161.7-93370 | 1.086229 | 1.040945 | 1.114062 | 0.982256 | 1.337877 | 1.341061 | 0.571891 | 0.649034 | **1.015420** | 0.943192 | 0.779065 | 0.976357 | 0.870161 | 0.741262 | 0.637798 | 0.860317 | 0.697957 | **0.813264** |
| Mki67-ENSMUST00000033310.8-47180 | 0.877049 | 0.827911 | 0.944245 | 0.847034 | 1.339396 | 1.352090 | 0.962424 | 0.960431 | **1.013823** | 0.557128 | 0.476553 | 0.987070 | 1.137857 | 0.948167 | 1.042727 | 1.016564 | 0.915878 | **0.885243** |
| Clec10a-ENSMUST00000102571.9-63957 | 0.755535 | 0.814907 | 0.893489 | 0.724032 | 1.060950 | 0.997665 | 1.194004 | 1.259573 | **0.962519** | 0.462654 | 0.421443 | 0.529962 | 0.447249 | 0.315422 | 0.268105 | 0.427145 | 0.308103 | **0.397510** |
| Cd86-ENSMUST00000089620.10-80399 | 0.898971 | 0.716926 | 0.906513 | 0.778399 | 1.197993 | 1.030412 | 1.121894 | 0.946434 | **0.949693** | 1.024017 | 0.815867 | 0.861574 | 0.740724 | 0.997620 | 0.812456 | 0.947160 | 0.763882 | **0.870413** |
| Mbp-ENSMUST00000091789.10-86889 | 1.015270 | 0.962937 | 0.898606 | 0.837249 | 0.796287 | 0.883571 | 1.029567 | 1.038115 | **0.932700** | 0.757454 | 0.720988 | 0.741707 | 0.700810 | 0.776691 | 0.708834 | 0.660496 | 0.708819 | **0.721975** |
| Qpct-ENSMUST00000040789.4-84843 | 0.787901 | 0.783741 | 0.890510 | 0.882279 | 0.762503 | 0.790640 | 1.139337 | 1.069155 | **0.888258** | 0.328809 | 0.279089 | 0.373557 | 0.318735 | 0.292539 | 0.256660 | 0.221078 | 0.223096 | **0.286695** |
| Nt5e-ENSMUST00000034992.7-55687 | 1.188692 | 1.148785 | 0.845649 | 0.998052 | 0.469866 | 0.483379 | 0.996951 | 0.953903 | **0.885660** | 1.227475 | 1.175243 | 0.637880 | 0.634617 | 0.997394 | 1.043226 | 0.581787 | 0.555857 | **0.856685** |
| Cnot2-ENSMUST00000105267.7-60575 | 0.942587 | 0.771979 | 0.916467 | 0.887126 | 1.077936 | 0.999005 | 0.710357 | 0.617272 | **0.865341** | 0.614069 | 0.518633 | 0.695643 | 0.665526 | 0.663648 | 0.636647 | 0.638397 | 0.568955 | **0.625190** |
| Birc3-ENSMUST00000013949.14-52943 | 0.749749 | 0.728739 | 0.807137 | 0.767226 | 0.931354 | 0.913820 | 0.981434 | 0.836350 | **0.839476** | 0.806408 | 0.689752 | 0.795472 | 0.731104 | 0.844408 | 0.754219 | 0.832475 | 0.736730 | **0.773821** |
| Lipa-ENSMUST00000178114.1-88424 | 0.665521 | 0.494872 | 0.566989 | 0.520366 | 1.366188 | 1.288117 | 0.835141 | 0.763349 | **0.812568** | 0.403938 | 0.350939 | 0.443453 | 0.412455 | 0.442939 | 0.402947 | 0.522476 | 0.441938 | **0.427635** |
| Ly96-ENSMUST00000026881.10-728 | 0.822157 | 0.752305 | 0.981773 | 0.868481 | 1.030776 | 0.951889 | 0.497722 | 0.504098 | **0.801150** | 0.850819 | 0.685687 | 0.837525 | 0.756205 | 0.749502 | 0.623940 | 0.773289 | 0.551901 | **0.728608** |
| Aurkb-ENSMUST00000108666.7-63704 | 0.755591 | 0.731594 | 0.743759 | 0.655612 | 0.832234 | 0.772858 | 0.916967 | 0.859178 | **0.783474** | 0.384028 | 0.291286 | 0.658885 | 0.636055 | 0.570535 | 0.565312 | 0.745482 | 0.610712 | **0.557787** |
| Il18-ENSMUST00000180021.1-54250 | 0.638183 | 0.565939 | 0.561476 | 0.468044 | 1.078144 | 0.943953 | 0.928925 | 0.915865 | **0.762566** | 0.704761 | 0.584423 | 0.528089 | 0.448278 | 0.451171 | 0.366258 | 0.526821 | 0.404859 | **0.501833** |
| Cxcl9-ENSMUST00000113093.4-29446 | 0.431273 | 0.230730 | 0.673493 | 0.166905 | 1.219563 | 0.512451 | 1.845865 | 0.838402 | **0.739835** | 0.255035 | 0.150986 | 0.596347 | 0.316719 | 1.470823 | 0.827448 | 1.041206 | 0.650289 | **0.663606** |
| Tnf-ENSMUST00000025263.14-83440 | 0.765267 | 0.720282 | 0.680238 | 0.600482 | 0.760899 | 0.633274 | 0.882694 | 0.729937 | **0.721634** | 0.867521 | 0.663168 | 0.937104 | 0.732061 | 1.285031 | 1.031460 | 1.216116 | 0.938300 | **0.958845** |
| Ticam1-ENSMUST00000058136.8-84443 | 0.695011 | 0.684023 | 0.680194 | 0.690604 | 0.668332 | 0.653377 | 0.797497 | 0.782471 | **0.706439** | 0.490989 | 0.447751 | 0.465395 | 0.436767 | 0.479156 | 0.451430 | 0.495504 | 0.443773 | **0.463845** |
| Ccl3-ENSMUST00000001008.5-65063 | 0.711060 | 0.676938 | 0.772192 | 0.568055 | 0.637718 | 0.504607 | 0.892762 | 0.868017 | **0.703919** | 1.145021 | 0.966968 | 0.699506 | 0.567798 | 1.288784 | 1.009244 | 1.085136 | 0.887418 | **0.956234** |
| Cd274-ENSMUST00000016640.7-88316 | 0.533624 | 0.373969 | 0.550374 | 0.357099 | 0.503761 | 0.338177 | 1.332131 | 0.917466 | **0.613325** | 1.170063 | 1.060727 | 0.880319 | 0.660901 | 1.077744 | 0.803602 | 1.160011 | 0.958518 | **0.971486** |
| Thy1-ENSMUST00000114840.1-53998 | 0.201527 | 0.178275 | 0.113165 | 0.224042 | 0.338011 | 0.413680 | 0.100842 | 0.113402 | **0.210368** | 0.497888 | 0.505409 | 0.933723 | 0.975909 | 1.253901 | 1.310981 | 0.968303 | 1.294552 | **0.967583** |
| Vegfa-ENSMUST00000167860.7-84052 | 0.450057 | 0.426507 | 0.337225 | 0.329724 | 0.129982 | 0.134377 | 0.576482 | 0.492868 | **0.359653** | 1.695482 | 1.551880 | 0.442609 | 0.354134 | 0.659665 | 0.471932 | 1.024320 | 0.840337 | **0.880045** |
| Il2rb-ENSMUST00000089398.7-77753 | 0.163935 | 0.119775 | 0.084396 | 0.168015 | 0.268000 | 0.337431 | 0.173682 | 0.191671 | **0.188363** | 0.470913 | 0.482514 | 0.835539 | 0.876705 | 1.113597 | 1.209423 | 0.781192 | 1.122916 | **0.861600** |

| | | | | | | | | | | | | | | | | | | |
|---|---|---|---|---|---|---|---|---|---|---|---|---|---|---|---|---|---|---|
| Gzmb-ENSMUST00000015581.4-75403 | 0.172159 | 0.138123 | 0.070226 | 0.175291 | 0.285458 | 0.317778 | 0.239909 | 0.240511 | **0.204932** | 0.518558 | 0.501430 | 0.686663 | 0.729400 | 1.148615 | 1.205986 | 0.732261 | 1.044541 | **0.820932** |
| Socs1-ENSMUST00000038099.4-79289 | 0.310096 | 0.332112 | 0.203385 | 0.159531 | 0.394040 | 0.256516 | 0.136928 | 0.097859 | **0.236308** | 1.024506 | 0.893459 | 0.786329 | 0.719279 | 0.893985 | 0.755952 | 0.727521 | 0.635335 | **0.804546** |
| Nkg7-ENSMUST00000070518.3-42115 | 0.186968 | 0.134731 | 0.097017 | 0.208834 | 0.318199 | 0.372702 | 0.485687 | 0.449743 | **0.281735** | 0.446563 | 0.438390 | 0.727439 | 0.774905 | 1.030960 | 1.088501 | 0.758478 | 1.076573 | **0.792726** |
| C3-ENSMUST0000024988.14-84525 | 0.382475 | 0.298365 | 0.487470 | 0.399194 | 0.639732 | 0.437552 | 1.086252 | 0.860388 | **0.573929** | 0.874217 | 0.840585 | 0.662057 | 0.455971 | 0.831585 | 0.550181 | 0.857782 | 0.677318 | **0.718712** |
| Clec4e-ENSMUST00000032239.10-37602 | 0.525797 | 0.467035 | 0.546032 | 0.591977 | 0.422314 | 0.394846 | 0.918601 | 0.841369 | **0.588496** | 0.915057 | 0.878227 | 0.555212 | 0.453130 | 0.855264 | 0.714242 | 0.508057 | 0.467632 | **0.668353** |
| Cd38-ENSMUST00000030964.5-27997 | 0.431408 | 0.430891 | 0.969173 | 0.624965 | 0.931678 | 0.791237 | 0.446494 | 0.404596 | **0.628805** | 0.534416 | 0.319911 | 0.640830 | 0.582933 | 0.556191 | 0.399970 | 1.260095 | 0.986919 | **0.660158** |
| Cd69-ENSMUST00000032259.5-38083 | 0.184446 | 0.139005 | 0.187733 | 0.119811 | 0.260643 | 0.187156 | 0.249336 | 0.173220 | **0.187669** | 0.616973 | 0.507276 | 0.640853 | 0.551181 | 0.793325 | 0.679706 | 0.732574 | 0.720407 | **0.655287** |
| Trbc1-ENSMUST00000192856.5-34295 | 0.150514 | 0.097739 | 0.046211 | 0.127115 | 0.219235 | 0.263754 | 0.112351 | 0.103524 | **0.140055** | 0.343260 | 0.318064 | 0.596068 | 0.645471 | 0.882401 | 0.923935 | 0.627803 | 0.874464 | **0.651433** |
| Mmp12-ENSMUST00000005950.11-52922 | 0.463980 | 0.311008 | 0.235267 | 0.226124 | 0.132514 | 0.092007 | 0.540217 | 0.488325 | **0.311180** | 1.047164 | 0.855033 | 0.484199 | 0.380429 | 0.444766 | 0.325170 | 0.941719 | 0.699010 | **0.647186** |
| Ccl5-ENSMUST00000035938.2-65053 | 0.558109 | 0.344995 | 0.600509 | 0.281811 | 0.697144 | 0.523784 | 0.748911 | 0.455830 | **0.526387** | 0.609824 | 0.459039 | 0.620299 | 0.509645 | 0.804695 | 0.669842 | 0.750458 | 0.726517 | **0.643790** |
| Tnfsf13b-ENSMUST00000207792.1-48244 | 0.274382 | 0.147491 | 0.158266 | 0.095757 | 0.351558 | 0.195457 | 0.380398 | 0.189963 | **0.224159** | 0.263121 | 0.156929 | 0.756841 | 0.617241 | 0.882882 | 0.796699 | 0.826967 | 0.597024 | **0.612213** |
| Tnfrsf9-ENSMUST00000030808.9-25835 | 0.239366 | 0.198629 | 0.156844 | 0.151818 | 0.189481 | 0.225411 | 0.290894 | 0.233280 | **0.210715** | 0.599763 | 0.541092 | 0.454969 | 0.499143 | 0.670090 | 0.688867 | 0.554751 | 0.719872 | **0.591068** |
| Jak2-ENSMUST00000065796.9-88304 | 0.610721 | 0.546144 | 0.643466 | 0.630813 | 0.693889 | 0.682929 | 0.667871 | 0.538266 | **0.626762** | 0.704932 | 0.656498 | 0.626684 | 0.575988 | 0.600898 | 0.531227 | 0.495777 | 0.478777 | **0.583848** |
| Trbc2-ENSMUST00000103299.2-34304 | 0.167334 | 0.166190 | 0.147061 | 0.180948 | 0.284258 | 0.348292 | 0.174485 | 0.164286 | **0.204107** | 0.284052 | 0.249933 | 0.455393 | 0.457203 | 0.695346 | 0.719310 | 0.585877 | 0.835200 | **0.535289** |
| Gzma-ENSMUST00000023897.5-73199 | 0.110652 | 0.074775 | 0.053492 | 0.104592 | 0.231857 | 0.223614 | 0.186016 | 0.181296 | **0.145787** | 0.343135 | 0.316191 | 0.580594 | 0.583183 | 0.750472 | 0.751967 | 0.389114 | 0.529229 | **0.530486** |
| Ighm-ENSMUST00000177715.7-70202 | 0.467775 | 0.347444 | 0.517412 | 0.381240 | 0.720117 | 0.550453 | 0.352964 | 0.303512 | **0.455115** | 0.343019 | 0.324698 | 0.494264 | 0.528111 | 0.649613 | 0.650141 | 0.530879 | 0.712342 | **0.529133** |
| Ctsw-ENSMUST00000025844.4-87236 | 0.153962 | 0.111568 | 0.065875 | 0.150103 | 0.242828 | 0.299730 | 0.141502 | 0.139696 | **0.163158** | 0.241000 | 0.252983 | 0.526750 | 0.568618 | 0.603650 | 0.675621 | 0.545471 | 0.815203 | **0.528662** |
| Slc25a37-ENSMUST00000037064.4-75812 | 0.757008 | 0.584068 | 0.686756 | 0.799798 | 0.781113 | 0.779693 | 0.540901 | 0.523378 | **0.681589** | 0.574373 | 0.515885 | 0.494908 | 0.536773 | 0.592490 | 0.544727 | 0.482944 | 0.486052 | **0.528519** |
| Lck-ENSMUST00000102596.7-24289 | 0.133952 | 0.131510 | 0.084728 | 0.141140 | 0.246908 | 0.287369 | 0.151076 | 0.139171 | **0.164482** | 0.268496 | 0.275384 | 0.472737 | 0.521960 | 0.646634 | 0.696125 | 0.508236 | 0.770400 | **0.519996** |
| Tlr4-ENSMUST00000048096.11-22134 | 0.472747 | 0.515522 | 0.615631 | 0.517616 | 0.764048 | 0.690152 | 0.611369 | 0.504875 | **0.586495** | 0.723102 | 0.660408 | 0.520492 | 0.505889 | 0.427190 | 0.373152 | 0.506759 | 0.417344 | **0.516792** |
| Il12rb1-ENSMUST00000000808.7-50283 | 0.109279 | 0.064010 | 0.096725 | 0.085728 | 0.202339 | 0.117880 | 0.486282 | 0.217774 | **0.172502** | 0.323725 | 0.209135 | 0.554142 | 0.460898 | 0.674234 | 0.513966 | 0.672604 | 0.622397 | **0.503888** |
| Nod1-ENSMUST00000168172.3-34966 | 0.365392 | 0.319640 | 0.451080 | 0.324706 | 0.465706 | 0.370750 | 0.344408 | 0.275960 | **0.364705** | 0.559210 | 0.470411 | 0.474464 | 0.403356 | 0.607395 | 0.441946 | 0.532243 | 0.453828 | **0.492857** |
| Il1r2-ENSMUST00000027243.12-1427 | 0.371505 | 0.233353 | 0.293895 | 0.239918 | 0.234483 | 0.211248 | 0.327889 | 0.251619 | **0.270489** | 0.948402 | 0.796735 | 0.482098 | 0.308966 | 0.435640 | 0.258533 | 0.384041 | 0.305832 | **0.490031** |
| Sell-ENSMUST00000192047.5-5643 | 0.124714 | 0.128569 | 0.130564 | 0.172030 | 0.156949 | 0.196181 | 0.275636 | 0.301062 | **0.185713** | 0.646148 | 0.634933 | 0.412259 | 0.379599 | 0.452923 | 0.433888 | 0.382681 | 0.564746 | **0.488397** |
| Stat5a-ENSMUST00000107356.7-66194 | 0.541520 | 0.553822 | 0.673954 | 0.575587 | 0.753761 | 0.699182 | 0.722352 | 0.699680 | **0.652482** | 0.356600 | 0.333368 | 0.533879 | 0.555809 | 0.522163 | 0.505992 | 0.547301 | 0.525428 | **0.485068** |
| Il15-ENSMUST00000209363.1-50703 | 0.599343 | 0.540752 | 0.585683 | 0.525168 | 0.613248 | 0.524438 | 0.632671 | 0.586282 | **0.575948** | 0.606249 | 0.516530 | 0.543693 | 0.461487 | 0.425186 | 0.358280 | 0.507892 | 0.380661 | **0.474997** |
| Xcl1-ENSMUST00000027860.7-5687 | 0.207207 | 0.104905 | 0.059792 | 0.167960 | 0.240300 | 0.273394 | 0.081431 | 0.064424 | **0.149927** | 0.286733 | 0.290702 | 0.456911 | 0.487411 | 0.478651 | 0.500366 | 0.534534 | 0.745301 | **0.472576** |
| Cd37-ENSMUST00000209779.1-42572 | 0.632670 | 0.745407 | 0.676145 | 0.647295 | 0.857030 | 0.868101 | 0.486078 | 0.554725 | **0.683432** | 0.548617 | 0.495059 | 0.481182 | 0.453361 | 0.507523 | 0.482588 | 0.369572 | 0.388610 | **0.465814** |
| Ccnd2-ENSMUST00000000188.11-37884 | 0.216149 | 0.197756 | 0.210578 | 0.225994 | 0.447448 | 0.359027 | 0.235176 | 0.189406 | **0.260192** | 0.450930 | 0.388919 | 0.512836 | 0.464348 | 0.410523 | 0.391112 | 0.570147 | 0.527215 | **0.464504** |
| Tlr3-ENSMUST00000209772.1-49458 | 0.297374 | 0.273408 | 0.271367 | 0.253761 | 0.428018 | 0.396800 | 0.196362 | 0.151683 | **0.283597** | 0.499821 | 0.369007 | 0.550260 | 0.482614 | 0.410182 | 0.385072 | 0.588612 | 0.427151 | **0.464090** |
| Tnfsf8-ENSMUST00000030047.2-22108 | 0.296649 | 0.194379 | 0.201963 | 0.180568 | 0.243014 | 0.197127 | 0.194956 | 0.132975 | **0.205204** | 0.487026 | 0.413151 | 0.580809 | 0.529948 | 0.427428 | 0.392210 | 0.469505 | 0.407973 | **0.463506** |
| Il15ra-ENSMUST00000138349.7-7415 | 0.292681 | 0.235548 | 0.303587 | 0.238918 | 0.425756 | 0.318917 | 0.326470 | 0.240371 | **0.297781** | 0.472146 | 0.395929 | 0.533844 | 0.490710 | 0.490009 | 0.435264 | 0.459000 | 0.422491 | **0.462424** |
| Klrk1-ENSMUST00000168919.7-38128 | 0.202750 | 0.155428 | 0.145196 | 0.137774 | 0.252575 | 0.200161 | 0.289128 | 0.156886 | **0.192487** | 0.390085 | 0.330904 | 0.445200 | 0.383133 | 0.484035 | 0.414283 | 0.419322 | 0.521957 | **0.423615** |
| Tfrc-ENSMUST00000023486.14-80222 | 0.612057 | 0.695541 | 0.592186 | 0.704128 | 0.817961 | 0.858117 | 0.512328 | 0.526800 | **0.664890** | 0.322927 | 0.335044 | 0.476477 | 0.481190 | 0.442279 | 0.426092 | 0.446297 | 0.458248 | **0.423569** |
| Nod2-ENSMUST00000118370.7-51093 | 0.164614 | 0.137147 | 0.162182 | 0.223456 | 0.273015 | 0.262682 | 0.513475 | 0.411190 | **0.268470** | 0.508516 | 0.452761 | 0.386024 | 0.313425 | 0.411149 | 0.296240 | 0.401656 | 0.338218 | **0.388499** |
| S100a8-ENSMUST00000069927.9-17272 | 0.302033 | 0.277206 | 0.273380 | 0.225087 | 0.202277 | 0.180691 | 0.153889 | 0.150160 | **0.220590** | 0.779560 | 0.663791 | 0.323587 | 0.218152 | 0.356514 | 0.234171 | 0.271114 | 0.261048 | **0.388492** |
| Cd40-ENSMUST00000017799.11-13689 | 0.539546 | 0.486133 | 0.451583 | 0.384332 | 0.638956 | 0.517097 | 0.642774 | 0.449264 | **0.513711** | 0.350044 | 0.274020 | 0.444024 | 0.344137 | 0.495364 | 0.359401 | 0.449475 | 0.325563 | **0.380253** |
| Tnfrsf18-ENSMUST00000103173.9-26194 | 0.148521 | 0.164163 | 0.136689 | 0.145156 | 0.189239 | 0.232471 | 0.160627 | 0.146676 | **0.165443** | 0.286498 | 0.296207 | 0.341990 | 0.363550 | 0.432154 | 0.442901 | 0.324998 | 0.482335 | **0.371329** |
| Zap70-ENSMUST00000027291.6-1234 | 0.076567 | 0.064889 | 0.042772 | 0.097495 | 0.113407 | 0.169965 | 0.064545 | 0.074913 | **0.088069** | 0.178273 | 0.191653 | 0.343758 | 0.387826 | 0.444926 | 0.501162 | 0.349666 | 0.548349 | **0.368201** |
| Stat4-ENSMUST00000027277.6-1721 | 0.231815 | 0.262876 | 0.374255 | 0.358467 | 0.415310 | 0.383278 | 0.298752 | 0.294859 | **0.327451** | 0.378074 | 0.364438 | 0.267147 | 0.282691 | 0.370244 | 0.368842 | 0.398299 | 0.473836 | **0.362946** |
| Bcl6-ENSMUST00000023151.5-80026 | 0.525876 | 0.404475 | 0.485748 | 0.515043 | 0.486997 | 0.413328 | 0.322991 | 0.286731 | **0.430149** | 0.364391 | 0.310341 | 0.312618 | 0.269498 | 0.504662 | 0.416244 | 0.394718 | 0.325476 | **0.362244** |
| Mapk8-ENSMUST00000111945.8-74249 | 0.424831 | 0.392717 | 0.420295 | 0.442648 | 0.415852 | 0.376900 | 0.129041 | 0.123283 | **0.340696** | 0.352656 | 0.323023 | 0.435770 | 0.380748 | 0.371377 | 0.346006 | 0.335418 | 0.335827 | **0.360103** |
| Fscn1-ENSMUST0000031565.14-32540 | 0.433852 | 0.375191 | 0.348153 | 0.351992 | 0.484702 | 0.513540 | 0.448491 | 0.444709 | **0.425079** | 0.284032 | 0.297566 | 0.342363 | 0.377296 | 0.368058 | 0.396864 | 0.296098 | 0.372756 | **0.341879** |
| Tnfsf14-ENSMUST00000005976.6-84524 | 0.264548 | 0.216883 | 0.171368 | 0.224363 | 0.323358 | 0.263158 | 0.401448 | 0.359962 | **0.278136** | 0.402720 | 0.374208 | 0.402280 | 0.336877 | 0.313824 | 0.283745 | 0.292420 | 0.271939 | **0.334752** |
| Klra7-ENSMUST00000049304.13-38174 | 0.073766 | 0.040209 | 0.029587 | 0.096899 | 0.148080 | 0.131618 | 0.059596 | 0.048787 | **0.078568** | 0.188412 | 0.198723 | 0.289114 | 0.331336 | 0.446692 | 0.485392 | 0.286395 | 0.446855 | **0.334115** |
| Cst7-ENSMUST00000089200.2-12723 | 0.244855 | 0.159237 | 0.122650 | 0.183587 | 0.294971 | 0.282758 | 0.166889 | 0.124147 | **0.197387** | 0.180298 | 0.188406 | 0.305649 | 0.334301 | 0.359116 | 0.382634 | 0.345604 | 0.515832 | **0.326480** |
| Mgst1-ENSMUST00000008684.10-38555 | 0.486814 | 0.324561 | 0.549575 | 0.411687 | 0.830529 | 0.694724 | 0.876279 | 0.696102 | **0.608784** | 0.483290 | 0.456367 | 0.311833 | 0.245104 | 0.252694 | 0.154036 | 0.392386 | 0.287968 | **0.322960** |
| Il2ra-ENSMUST00000028111.5-7410 | 0.096748 | 0.076260 | 0.075713 | 0.055848 | 0.084145 | 0.095885 | 0.043219 | 0.027245 | **0.069383** | 0.257486 | 0.229144 | 0.399771 | 0.384924 | 0.403184 | 0.451209 | 0.224218 | 0.232444 | **0.322798** |
| Klrc1-ENSMUST00000032270.12-38138 | 0.053654 | 0.037378 | 0.027563 | 0.091857 | 0.099096 | 0.123220 | 0.058926 | 0.076585 | **0.071035** | 0.164820 | 0.184672 | 0.298727 | 0.318621 | 0.368443 | 0.414787 | 0.298585 | 0.478175 | **0.315854** |
| Ltb-ENSMUST00000025262.5-83420 | 0.130389 | 0.122117 | 0.139466 | 0.110431 | 0.164410 | 0.200919 | 0.075604 | 0.067232 | **0.126321** | 0.317373 | 0.312490 | 0.251069 | 0.247484 | 0.344183 | 0.338147 | 0.291148 | 0.412004 | **0.314237** |
| Cd2-ENSMUST00000029456.4-18064 | 0.139446 | 0.138216 | 0.122724 | 0.153584 | 0.183128 | 0.204561 | 0.105689 | 0.107465 | **0.144352** | 0.208261 | 0.224029 | 0.267096 | 0.297345 | 0.386960 | 0.390165 | 0.294736 | 0.443510 | **0.314013** |
| Il18rap-ENSMUST00000027237.11-1453 | 0.244936 | 0.232980 | 0.244792 | 0.274326 | 0.288203 | 0.266503 | 0.475429 | 0.369830 | **0.299625** | 0.257866 | 0.309703 | 0.321070 | 0.343684 | 0.330985 | 0.348735 | 0.236854 | 0.336445 | **0.310668** |
| Tgfb3-ENSMUST00000003687.6-69288 | 0.174652 | 0.195369 | 0.152988 | 0.179894 | 0.154214 | 0.181640 | 0.240944 | 0.209003 | **0.186088** | 0.326717 | 0.268015 | 0.199049 | 0.187039 | 0.370944 | 0.362975 | 0.340884 | 0.296474 | **0.294012** |
| Lat-ENSMUST00000032997.7-46345 | 0.100561 | 0.103764 | 0.062321 | 0.107124 | 0.145488 | 0.205544 | 0.099620 | 0.109587 | **0.116751** | 0.147836 | 0.139232 | 0.230360 | 0.276038 | 0.405020 | 0.418863 | 0.299894 | 0.427403 | **0.293081** |
| Trac-ENSMUST00000198398.4-75126 | 0.088809 | 0.105422 | 0.099067 | 0.109445 | 0.218062 | 0.279384 | 0.127596 | 0.138844 | **0.145829** | 0.114773 | 0.116959 | 0.178372 | 0.222391 | 0.359929 | 0.390122 | 0.304060 | 0.438816 | **0.265678** |

| | | | | | | | | | | | | | | | | | | |
|---|---|---|---|---|---|---|---|---|---|---|---|---|---|---|---|---|---|---|
| Mmp9-ENSMUST00000017881.2-13675 | 0.167969 | 0.198980 | 0.157018 | 0.177572 | 0.140584 | 0.195257 | 0.244887 | 0.223839 | **0.188263** | 0.314896 | 0.324364 | 0.220632 | 0.213325 | 0.306872 | 0.295866 | 0.192899 | 0.241695 | **0.263819** |
| Tbx21-ENSMUST00000001484.2-65761 | 0.034030 | 0.049075 | 0.027517 | 0.076583 | 0.083228 | 0.090648 | 0.056651 | 0.052731 | **0.058808** | 0.130163 | 0.161101 | 0.257531 | 0.269279 | 0.308671 | 0.341930 | 0.233422 | 0.400245 | **0.262793** |
| Ada-ENSMUST00000017841.3-13524 | 0.509150 | 0.462654 | 0.529447 | 0.387133 | 0.634389 | 0.551526 | 0.439865 | 0.404871 | **0.489879** | 0.182512 | 0.142150 | 0.361599 | 0.314232 | 0.238970 | 0.198370 | 0.343140 | 0.249954 | **0.253866** |
| Prf1-ENSMUST00000035419.5-58753 | 0.032878 | 0.023775 | 0.003227 | 0.022370 | 0.076295 | 0.100602 | 0.062418 | 0.069538 | **0.048888** | 0.131485 | 0.128859 | 0.211642 | 0.250801 | 0.335571 | 0.369493 | 0.193924 | 0.315300 | **0.242134** |
| Tnfsf10-ENSMUST00000046383.11-15025 | 0.117921 | 0.086300 | 0.083750 | 0.102721 | 0.146102 | 0.118822 | 0.099795 | 0.054762 | **0.101272** | 0.258376 | 0.223718 | 0.271008 | 0.251178 | 0.197868 | 0.196328 | 0.245810 | 0.244099 | **0.236048** |
| Tcf7-ENSMUST00000086844.9-62791 | 0.076374 | 0.065758 | 0.065070 | 0.081521 | 0.103155 | 0.152157 | 0.093472 | 0.080429 | **0.089742** | 0.114508 | 0.144479 | 0.195673 | 0.225429 | 0.261978 | 0.301175 | 0.237087 | 0.402066 | **0.235299** |
| Cd247-ENSMUST00000005907.11-5752 | 0.052005 | 0.062887 | 0.028208 | 0.080137 | 0.102687 | 0.139225 | 0.076943 | 0.081746 | **0.077980** | 0.104005 | 0.127812 | 0.200527 | 0.212615 | 0.316423 | 0.342520 | 0.217705 | 0.344164 | **0.233221** |
| Bcl2-ENSMUST00000112751.1-3862 | 0.238924 | 0.218820 | 0.163896 | 0.197916 | 0.235686 | 0.265112 | 0.216369 | 0.231701 | **0.221053** | 0.127033 | 0.134164 | 0.215373 | 0.224185 | 0.256220 | 0.280218 | 0.218551 | 0.326391 | **0.222767** |
| Cd244-ENSMUST00000004829.12-6071 | 0.319622 | 0.206539 | 0.214008 | 0.294997 | 0.342256 | 0.301999 | 0.249081 | 0.200870 | **0.266171** | 0.209568 | 0.196737 | 0.229598 | 0.211019 | 0.252854 | 0.234432 | 0.209569 | 0.220729 | **0.220563** |
| Cblb-ENSMUST00000114471.1-80770 | 0.151234 | 0.167502 | 0.185350 | 0.171682 | 0.225205 | 0.262320 | 0.186464 | 0.209350 | **0.194888** | 0.159315 | 0.160685 | 0.206621 | 0.229576 | 0.229442 | 0.253543 | 0.217849 | 0.296868 | **0.219238** |
| Cd3g-ENSMUST00000002101.11-54111 | 0.087397 | 0.075478 | 0.069078 | 0.084218 | 0.138864 | 0.183680 | 0.095895 | 0.088616 | **0.102903** | 0.095023 | 0.105444 | 0.148961 | 0.170752 | 0.285556 | 0.293238 | 0.257615 | 0.376386 | **0.216622** |
| Il1a-ENSMUST00000028882.1-12106 | 0.035326 | 0.026950 | 0.022580 | 0.039372 | 0.016710 | 0.012864 | 0.062158 | 0.049389 | **0.033169** | 0.283110 | 0.218786 | 0.138117 | 0.104050 | 0.199389 | 0.132996 | 0.359449 | 0.269524 | **0.213178** |
| Cd36-ENSMUST00000170051.7-26684 | 0.270133 | 0.268994 | 0.242841 | 0.220407 | 0.103982 | 0.112481 | 0.255562 | 0.228380 | **0.212848** | 0.165697 | 0.158056 | 0.111564 | 0.092855 | 0.452304 | 0.389504 | 0.151556 | 0.132062 | **0.206700** |
| Il7r-ENSMUST00000003981.4-76570 | 0.078275 | 0.109291 | 0.093477 | 0.082465 | 0.071307 | 0.078924 | 0.128314 | 0.099972 | **0.092753** | 0.301496 | 0.264636 | 0.115739 | 0.099505 | 0.245440 | 0.207917 | 0.199961 | 0.190119 | **0.203102** |
| Cxcr3-ENSMUST00000056614.6-91654 | 0.179228 | 0.121643 | 0.208012 | 0.142862 | 0.280106 | 0.276144 | 0.142103 | 0.139995 | **0.186262** | 0.110167 | 0.103597 | 0.205313 | 0.194909 | 0.233689 | 0.230361 | 0.250542 | 0.290970 | **0.202444** |
| Csf1-ENSMUST00000014743.9-18457 | 0.108700 | 0.147195 | 0.150832 | 0.184546 | 0.140313 | 0.194850 | 0.174394 | 0.145863 | **0.155837** | 0.226677 | 0.228908 | 0.193252 | 0.219525 | 0.191070 | 0.210461 | 0.152041 | 0.188005 | **0.201242** |
| Txk-ENSMUST0000198464.2-28791 | 0.054759 | 0.025256 | 0.016709 | 0.051951 | 0.058915 | 0.066698 | 0.045466 | 0.044928 | **0.045585** | 0.118854 | 0.102061 | 0.196493 | 0.224381 | 0.252960 | 0.286780 | 0.154173 | 0.273904 | **0.201201** |
| Gimap5-ENSMUST00000055558.5-34636 | 0.022333 | 0.042163 | 0.021779 | 0.039793 | 0.058863 | 0.093371 | 0.034678 | 0.044709 | **0.044711** | 0.098515 | 0.104048 | 0.164746 | 0.207102 | 0.242541 | 0.263900 | 0.179235 | 0.310153 | **0.196280** |
| Tigit-BD-custom-chr16-43648113 | 0.064445 | 0.049973 | 0.020592 | 0.054156 | 0.109883 | 0.153631 | 0.055243 | 0.050452 | **0.069797** | 0.102137 | 0.104741 | 0.169535 | 0.174582 | 0.224349 | 0.218967 | 0.201415 | 0.335423 | **0.191394** |
| Eomes-ENSMUST00000035020.14-57330 | 0.027014 | 0.018151 | 0.022157 | 0.046509 | 0.067019 | 0.082037 | 0.033772 | 0.028731 | **0.040674** | 0.084745 | 0.094306 | 0.168305 | 0.219417 | 0.220426 | 0.252769 | 0.168428 | 0.298906 | **0.188413** |
| Fas-ENSMUST00000025691.11-88416 | 0.181332 | 0.195769 | 0.293871 | 0.212785 | 0.336736 | 0.309395 | 0.248760 | 0.235387 | **0.251754** | 0.183349 | 0.164591 | 0.209160 | 0.153120 | 0.219730 | 0.192100 | 0.194581 | 0.152550 | **0.183648** |
| Itk-ENSMUST00000020664.12-62412 | 0.043359 | 0.041766 | 0.035618 | 0.072749 | 0.088604 | 0.102101 | 0.055780 | 0.065271 | **0.063156** | 0.080467 | 0.084901 | 0.119480 | 0.179932 | 0.241002 | 0.274652 | 0.185655 | 0.296741 | **0.182854** |
| Thbd-ENSMUST00000099270.4-12649 | 0.292861 | 0.272973 | 0.363739 | 0.382169 | 0.343199 | 0.368642 | 0.337532 | 0.342657 | **0.337971** | 0.311954 | 0.271525 | 0.169865 | 0.162778 | 0.142577 | 0.134094 | 0.119857 | 0.137244 | **0.181237** |
| Aqp9-ENSMUST00000074465.8-55103 | 0.145661 | 0.160397 | 0.130531 | 0.143280 | 0.076772 | 0.073666 | 0.196961 | 0.169828 | **0.137137** | 0.224636 | 0.217030 | 0.198013 | 0.172963 | 0.207269 | 0.166056 | 0.134929 | 0.126585 | **0.180935** |
| Ccr7-ENSMUST00000103134.3-65951 | 0.084294 | 0.062991 | 0.079819 | 0.052109 | 0.057481 | 0.070520 | 0.046577 | 0.050614 | **0.063051** | 0.159853 | 0.162642 | 0.110695 | 0.123791 | 0.233015 | 0.237087 | 0.136316 | 0.233199 | **0.174575** |
| Cd7-ENSMUST00000026159.5-67713 | 0.058838 | 0.048314 | 0.036699 | 0.076520 | 0.083663 | 0.102689 | 0.030831 | 0.032318 | **0.058734** | 0.079027 | 0.076379 | 0.130851 | 0.132041 | 0.174963 | 0.193228 | 0.201522 | 0.319267 | **0.163410** |
| Gimap7-ENSMUST00000052503.7-34625 | 0.048972 | 0.045487 | 0.021081 | 0.045664 | 0.076892 | 0.103914 | 0.079042 | 0.083895 | **0.063119** | 0.113569 | 0.109735 | 0.123031 | 0.129554 | 0.189104 | 0.203337 | 0.161921 | 0.251983 | **0.160279** |
| Cd3d-ENSMUST00000034602.7-54114 | 0.072619 | 0.067436 | 0.062937 | 0.071519 | 0.111712 | 0.159518 | 0.076770 | 0.079372 | **0.087735** | 0.083706 | 0.080135 | 0.113591 | 0.132095 | 0.201284 | 0.216543 | 0.185934 | 0.268478 | **0.160221** |
| Cd1d1-ENSMUST00000029717.3-16694 | 0.156479 | 0.122963 | 0.091315 | 0.123477 | 0.073439 | 0.106629 | 0.129913 | 0.108777 | **0.114124** | 0.169473 | 0.170053 | 0.187140 | 0.166441 | 0.155561 | 0.123444 | 0.129101 | 0.142911 | **0.155515** |
| Fosl1-ENSMUST00000025850.5-87228 | 0.125558 | 0.117981 | 0.177115 | 0.118657 | 0.183246 | 0.170763 | 0.204945 | 0.192188 | **0.161307** | 0.193096 | 0.180868 | 0.119914 | 0.143868 | 0.149239 | 0.161725 | 0.145736 | 0.149413 | **0.155482** |
| Traf6-ENSMUST00000004949.7-10898 | 0.240797 | 0.225899 | 0.237298 | 0.263803 | 0.207086 | 0.229062 | 0.136057 | 0.133538 | **0.209193** | 0.162989 | 0.141025 | 0.180935 | 0.178164 | 0.137763 | 0.116171 | 0.158736 | 0.144896 | **0.152585** |
| Prdm1-ENSMUST00000039174.10-57501 | 0.169369 | 0.090238 | 0.122993 | 0.109179 | 0.175961 | 0.157494 | 0.157516 | 0.124023 | **0.138347** | 0.171814 | 0.167449 | 0.135353 | 0.112258 | 0.149595 | 0.134809 | 0.187483 | 0.154717 | **0.151685** |
| Tlr9-ENSMUST00000062241.10-56431 | 0.178082 | 0.141017 | 0.182094 | 0.159362 | 0.134340 | 0.105819 | 0.064648 | 0.042884 | **0.126031** | 0.224338 | 0.152802 | 0.185085 | 0.147034 | 0.135473 | 0.091307 | 0.137058 | 0.103451 | **0.147068** |
| Fut4-ENSMUST00000061498.5-53030 | 0.243693 | 0.161699 | 0.160591 | 0.117351 | 0.249805 | 0.178643 | 0.222653 | 0.169547 | **0.187998** | 0.126849 | 0.124985 | 0.155991 | 0.137290 | 0.160539 | 0.135632 | 0.171095 | 0.141878 | **0.144282** |
| Cd34-ENSMUST00000016638.7-7059 | 0.191593 | 0.169317 | 0.121914 | 0.178672 | 0.181035 | 0.228487 | 0.145944 | 0.122597 | **0.167445** | 0.087256 | 0.129589 | 0.158425 | 0.173123 | 0.144664 | 0.168653 | 0.107954 | 0.154234 | **0.140487** |
| Cd79b-ENSMUST00000167143.1-66704 | 0.624166 | 0.485938 | 0.577837 | 0.389400 | 0.594392 | 0.455711 | 0.297239 | 0.282772 | **0.463432** | 0.167737 | 0.107327 | 0.143427 | 0.111023 | 0.153559 | 0.116368 | 0.173172 | 0.144746 | **0.139670** |
| Lag3-ENSMUST00000032217.1-37728 | 0.090717 | 0.049228 | 0.126418 | 0.080246 | 0.103446 | 0.117367 | 0.020852 | 0.027519 | **0.076974** | 0.073404 | 0.059843 | 0.097396 | 0.112098 | 0.164741 | 0.169457 | 0.186404 | 0.251067 | **0.139301** |
| Cd28-ENSMUST00000027165.2-2133 | 0.024278 | 0.040651 | 0.032322 | 0.029214 | 0.056364 | 0.080494 | 0.018569 | 0.022778 | **0.038084** | 0.082822 | 0.081322 | 0.118039 | 0.141641 | 0.147644 | 0.180806 | 0.125640 | 0.214174 | **0.136511** |
| Cd8a-ENSMUST00000066747.13-35536 | 0.019049 | 0.013258 | 0.019656 | 0.042445 | 0.046244 | 0.082064 | 0.054346 | 0.056058 | **0.041640** | 0.033169 | 0.036382 | 0.069341 | 0.088535 | 0.176666 | 0.192158 | 0.152632 | 0.235268 | **0.123019** |
| Cd8b1-ENSMUST00000065248.8-35529 | 0.010929 | 0.022761 | 0.015242 | 0.033479 | 0.062002 | 0.088076 | 0.047992 | 0.057676 | **0.042269** | 0.032187 | 0.036251 | 0.083866 | 0.087286 | 0.161245 | 0.176868 | 0.145649 | 0.230201 | **0.119194** |
| Tnfrsf4-ENSMUST00000030952.5-26202 | 0.091059 | 0.097810 | 0.097342 | 0.052165 | 0.159566 | 0.143572 | 0.071021 | 0.078269 | **0.098850** | 0.095578 | 0.079357 | 0.103401 | 0.084979 | 0.176389 | 0.153555 | 0.115920 | 0.137583 | **0.118345** |
| Cd6-ENSMUST00000080292.11-87771 | 0.052483 | 0.050277 | 0.038601 | 0.037366 | 0.075021 | 0.110699 | 0.059870 | 0.065427 | **0.061218** | 0.050160 | 0.056608 | 0.075508 | 0.083498 | 0.166598 | 0.168754 | 0.134306 | 0.208314 | **0.117968** |
| Klra21-NM-053151.1-234625 | 0.023112 | 0.012430 | 0.010861 | 0.047883 | 0.049804 | 0.049644 | 0.011585 | 0.007304 | **0.026578** | 0.075837 | 0.071377 | 0.127674 | 0.128999 | 0.129776 | 0.145539 | 0.092856 | 0.163338 | **0.116925** |
| Cd27-ENSMUST00000032486.12-37807 | 0.019719 | 0.044962 | 0.024809 | 0.041826 | 0.054019 | 0.078775 | 0.009924 | 0.020432 | **0.036808** | 0.056617 | 0.064580 | 0.099661 | 0.103608 | 0.138887 | 0.145995 | 0.105668 | 0.185021 | **0.112505** |
| Klra1-ENSMUST00000032288.5-38185 | 0.032126 | 0.018682 | 0.008675 | 0.040926 | 0.081476 | 0.087189 | 0.024586 | 0.021133 | **0.039349** | 0.062822 | 0.056975 | 0.093397 | 0.115869 | 0.134896 | 0.150584 | 0.094790 | 0.144794 | **0.106766** |
| Tcf4-ENSMUST00000114985.9-86599 | 0.161072 | 0.142991 | 0.175153 | 0.240695 | 0.566472 | 0.667755 | 0.254829 | 0.266152 | **0.309390** | 0.081109 | 0.089241 | 0.080795 | 0.103418 | 0.089438 | 0.098662 | 0.146274 | 0.157422 | **0.105795** |
| Ifng-ENSMUST00000068592.3-60655 | 0.021173 | 0.020763 | 0.008678 | 0.038192 | 0.040257 | 0.062816 | 0.056362 | 0.056299 | **0.038067** | 0.060711 | 0.062239 | 0.086214 | 0.080415 | 0.136798 | 0.149285 | 0.099523 | 0.163947 | **0.104891** |
| Cd5-ENSMUST00000025571.7-87742 | 0.052361 | 0.040037 | 0.048183 | 0.044106 | 0.081893 | 0.109690 | 0.044710 | 0.045844 | **0.058353** | 0.050669 | 0.058659 | 0.061934 | 0.080606 | 0.150356 | 0.159200 | 0.095514 | 0.150266 | **0.100900** |
| Cd3e-ENSMUST00000102832.1-54115 | 0.045447 | 0.041481 | 0.059356 | 0.057639 | 0.066913 | 0.099592 | 0.040486 | 0.041322 | **0.056530** | 0.045048 | 0.056928 | 0.073655 | 0.080726 | 0.149742 | 0.159231 | 0.087588 | 0.146568 | **0.099936** |
| Cxcr6-ENSMUST00000049810.7-57554 | 0.026542 | 0.051113 | 0.025820 | 0.040395 | 0.059306 | 0.094296 | 0.032790 | 0.028222 | **0.044810** | 0.040584 | 0.050764 | 0.058759 | 0.059261 | 0.127318 | 0.146321 | 0.124113 | 0.171840 | **0.097370** |
| Fasl-ENSMUST00000000834.3-5532 | 0.032341 | 0.022907 | 0.013818 | 0.037755 | 0.023089 | 0.029965 | 0.003276 | 0.008854 | **0.021501** | 0.071637 | 0.051908 | 0.089088 | 0.108104 | 0.118463 | 0.134703 | 0.078044 | 0.122356 | **0.096788** |
| Klra3-ENSMUST00000111998.8-38183 | 0.007294 | 0.007509 | 0.000000 | 0.021542 | 0.033469 | 0.030337 | 0.006160 | 0.005980 | **0.014036** | 0.052128 | 0.050333 | 0.094815 | 0.106192 | 0.103959 | 0.119336 | 0.096909 | 0.143254 | **0.095866** |
| Ctla4-ENSMUST00000027164.8-2142 | 0.044110 | 0.047444 | 0.041437 | 0.017186 | 0.097670 | 0.113774 | 0.028581 | 0.033123 | **0.052916** | 0.067985 | 0.074251 | 0.066912 | 0.083416 | 0.111754 | 0.121648 | 0.091983 | 0.146112 | **0.095508** |
| Pask-ENSMUST00000027493.3-3606 | 0.209950 | 0.200025 | 0.211809 | 0.148778 | 0.150577 | 0.158474 | 0.161495 | 0.179517 | **0.177578** | 0.059210 | 0.054005 | 0.120626 | 0.097192 | 0.098861 | 0.091295 | 0.110965 | 0.104557 | **0.092089** |
| Klrg1-ENSMUST00000032207.8-37485 | 0.025353 | 0.023862 | 0.012661 | 0.025728 | 0.045384 | 0.045754 | 0.013054 | 0.011386 | **0.025398** | 0.049536 | 0.049975 | 0.084957 | 0.112847 | 0.124595 | 0.143924 | 0.059692 | 0.092768 | **0.089787** |
| Cxcl1-ENSMUST00000031327.8-29363 | 0.026615 | 0.020947 | 0.009998 | 0.019173 | 0.013322 | 0.009150 | 0.011942 | 0.018272 | **0.016177** | 0.150366 | 0.119100 | 0.054013 | 0.050746 | 0.070513 | 0.061091 | 0.113732 | 0.095441 | **0.089375** |
| Cd160-ENSMUST00000107074.7-17893 | 0.030815 | 0.025416 | 0.007396 | 0.031990 | 0.041944 | 0.049873 | 0.011087 | 0.015983 | **0.026813** | 0.037355 | 0.029856 | 0.084640 | 0.092333 | 0.101119 | 0.096539 | 0.104192 | 0.157948 | **0.087998** |

| | | | | | | | | | | | | | | | | | | |
|---|---|---|---|---|---|---|---|---|---|---|---|---|---|---|---|---|---|---|
| Icos-ENSMUST00000102827.3-2147 | 0.034960 | 0.043984 | 0.033714 | 0.019315 | 0.072467 | 0.099392 | 0.020931 | 0.028300 | **0.044133** | 0.060986 | 0.055318 | 0.061068 | 0.075298 | 0.119982 | 0.135136 | 0.081236 | 0.112075 | **0.087637** |
| Lamp3-ENSMUST00000081880.5-79778 | 0.326828 | 0.358459 | 0.366026 | 0.350295 | 0.304431 | 0.308030 | 0.250102 | 0.239230 | **0.312925** | 0.089247 | 0.086016 | 0.062253 | 0.061579 | 0.083019 | 0.096735 | 0.102350 | 0.101705 | **0.085363** |
| Trdc-ENSMUST00000196323.1-75099 | 0.010695 | 0.021465 | 0.009184 | 0.034038 | 0.025944 | 0.043429 | 0.023837 | 0.021459 | **0.023757** | 0.052980 | 0.058212 | 0.079370 | 0.092315 | 0.078464 | 0.092961 | 0.081397 | 0.132737 | **0.083555** |
| Ncam1-ENSMUST00000194252.5-54263 | 0.094804 | 0.115136 | 0.085131 | 0.119684 | 0.100481 | 0.176682 | 0.066851 | 0.062872 | **0.102705** | 0.029088 | 0.055897 | 0.080844 | 0.118988 | 0.095548 | 0.121520 | 0.054503 | 0.105949 | **0.082792** |
| Tlr8-ENSMUST00000112170.1-93367 | 0.081260 | 0.103222 | 0.199820 | 0.131422 | 0.177950 | 0.162248 | 0.045918 | 0.055843 | **0.119710** | 0.123212 | 0.101771 | 0.069677 | 0.065519 | 0.069084 | 0.058328 | 0.089823 | 0.079865 | **0.082160** |
| S100a9-ENSMUST00000117167.1-17275 | 0.070990 | 0.073439 | 0.085010 | 0.079982 | 0.031229 | 0.025799 | 0.028744 | 0.034159 | **0.053669** | 0.177135 | 0.166796 | 0.054796 | 0.036834 | 0.068765 | 0.040483 | 0.057720 | 0.054711 | **0.082155** |
| Pik3ip1-ENSMUST00000045153.10-61217 | 0.048538 | 0.058775 | 0.026014 | 0.046880 | 0.064658 | 0.072753 | 0.075954 | 0.048980 | **0.055319** | 0.050299 | 0.053106 | 0.092910 | 0.079160 | 0.089136 | 0.079747 | 0.072195 | 0.106326 | **0.077860** |
| Pdcd1-ENSMUST00000027507.7-3686 | 0.022185 | 0.036765 | 0.082337 | 0.022995 | 0.060471 | 0.059224 | 0.029517 | 0.029333 | **0.042853** | 0.057816 | 0.051476 | 0.045777 | 0.046629 | 0.117239 | 0.110525 | 0.071493 | 0.107674 | **0.076079** |
| Cd22-ENSMUST00000019248.12-41479 | 0.118387 | 0.091517 | 0.089794 | 0.073901 | 0.134515 | 0.096899 | 0.076047 | 0.072421 | **0.094185** | 0.054184 | 0.036705 | 0.075989 | 0.059083 | 0.053703 | 0.027527 | 0.185233 | 0.105111 | **0.074692** |
| Il12rb2-ENSMUST00000117441.7-35319 | 0.019286 | 0.017791 | 0.025842 | 0.043144 | 0.026215 | 0.049534 | 0.047064 | 0.030523 | **0.032425** | 0.072527 | 0.073129 | 0.057836 | 0.062701 | 0.070831 | 0.073549 | 0.061844 | 0.105732 | **0.072269** |
| Dpp4-ENSMUST00000047812.7-9521 | 0.102420 | 0.073607 | 0.089387 | 0.082958 | 0.139844 | 0.114889 | 0.165321 | 0.107461 | **0.109486** | 0.080161 | 0.080607 | 0.066113 | 0.048047 | 0.072500 | 0.060793 | 0.072335 | 0.081926 | **0.070310** |
| Arg2-ENSMUST00000021550.6-68965 | 0.039304 | 0.053908 | 0.062906 | 0.060287 | 0.030968 | 0.032584 | 0.083615 | 0.092087 | **0.056957** | 0.154389 | 0.162557 | 0.033163 | 0.024099 | 0.047271 | 0.038933 | 0.050415 | 0.040535 | **0.068920** |
| Ccl17-ENSMUST00000034232.2-51374 | 0.067704 | 0.074591 | 0.068815 | 0.041869 | 0.073986 | 0.076634 | 0.043609 | 0.040849 | **0.061007** | 0.114655 | 0.119103 | 0.078792 | 0.064934 | 0.030795 | 0.029404 | 0.046920 | 0.040151 | **0.065594** |
| Tigit-ENSMUST00000096065.4-80560 | 0.042165 | 0.046804 | 0.031394 | 0.038004 | 0.065476 | 0.088425 | 0.021113 | 0.024025 | **0.044676** | 0.046726 | 0.051278 | 0.045007 | 0.039785 | 0.060138 | 0.068081 | 0.067895 | 0.134205 | **0.064140** |
| Flt3-ENSMUST00000049324.12-32846 | 0.094755 | 0.085747 | 0.082605 | 0.076615 | 0.085353 | 0.118387 | 0.087285 | 0.089938 | **0.090086** | 0.061115 | 0.076747 | 0.062549 | 0.056583 | 0.064630 | 0.071828 | 0.049800 | 0.068044 | **0.063912** |
| Trib2-ENSMUST00000020922.7-68015 | 0.109050 | 0.118218 | 0.077300 | 0.149317 | 0.107817 | 0.141743 | 0.082991 | 0.068145 | **0.106823** | 0.037563 | 0.053142 | 0.057111 | 0.075854 | 0.071596 | 0.091923 | 0.048730 | 0.073403 | **0.063665** |
| Igha-ENSMUST00000178282.2-70166 | 0.013767 | 0.004600 | 0.000000 | 0.027741 | 0.007082 | 0.011215 | 0.000000 | 0.001479 | **0.008236** | 0.013256 | 0.011603 | 0.067421 | 0.071396 | 0.074774 | 0.091578 | 0.052279 | 0.092028 | **0.059292** |
| Kit-ENSMUST00000005815.6-28962 | 0.026379 | 0.039425 | 0.040827 | 0.043891 | 0.063277 | 0.067586 | 0.044629 | 0.050561 | **0.047072** | 0.042334 | 0.050232 | 0.045675 | 0.052515 | 0.060622 | 0.077799 | 0.048721 | 0.090937 | **0.058605** |
| H2-Ob-ENSMUST00000095342.9-83196 | 0.150133 | 0.158996 | 0.157865 | 0.127859 | 0.099303 | 0.114650 | 0.079373 | 0.079083 | **0.120908** | 0.065040 | 0.059128 | 0.036922 | 0.033604 | 0.065067 | 0.056492 | 0.071762 | 0.065397 | **0.056676** |
| Igkc-ENSMUST00000103410.2-35486 | 0.034396 | 0.049466 | 0.027056 | 0.022051 | 0.019057 | 0.031859 | 0.051842 | 0.047540 | **0.035408** | 0.053919 | 0.058049 | 0.054547 | 0.057195 | 0.029469 | 0.036961 | 0.062808 | 0.088209 | **0.055145** |
| Il1rl1-ENSMUST00000097772.9-1442 | 0.073812 | 0.058505 | 0.058857 | 0.088382 | 0.082465 | 0.099243 | 0.057990 | 0.072377 | **0.073954** | 0.034758 | 0.057274 | 0.048503 | 0.051115 | 0.064045 | 0.077707 | 0.044163 | 0.062667 | **0.055029** |
| Cd4-ENSMUST00000024044.6-37727 | 0.055654 | 0.062083 | 0.098232 | 0.046350 | 0.102116 | 0.112273 | 0.040058 | 0.050569 | **0.070917** | 0.033641 | 0.034788 | 0.030996 | 0.033728 | 0.080259 | 0.074747 | 0.076944 | 0.066019 | **0.053890** |
| Il18r1-ENSMUST00000108044.3-1452 | 0.002991 | 0.012316 | 0.027753 | 0.015399 | 0.042993 | 0.057241 | 0.020005 | 0.017853 | **0.024569** | 0.029136 | 0.039524 | 0.044345 | 0.055392 | 0.052758 | 0.072172 | 0.049210 | 0.086379 | **0.053615** |
| Lef1-ENSMUST00000029611.13-19323 | 0.022510 | 0.021857 | 0.002911 | 0.033955 | 0.023139 | 0.029623 | 0.019421 | 0.021465 | **0.021860** | 0.026763 | 0.036606 | 0.051910 | 0.046296 | 0.061004 | 0.071448 | 0.047354 | 0.083847 | **0.053153** |
| Foxo1-ENSMUST00000053764.6-15810 | 0.040406 | 0.061819 | 0.044609 | 0.034303 | 0.108863 | 0.122129 | 0.068774 | 0.077394 | **0.069787** | 0.040459 | 0.034213 | 0.050710 | 0.055784 | 0.047492 | 0.050921 | 0.056387 | 0.066730 | **0.050337** |
| Gata3-ENSMUST00000102976.3-7307 | 0.006846 | 0.011796 | 0.007212 | 0.009073 | 0.006096 | 0.012202 | 0.000000 | 0.001845 | **0.006884** | 0.003451 | 0.003349 | 0.082407 | 0.077316 | 0.098702 | 0.116701 | 0.004882 | 0.007323 | **0.049266** |
| Adgrg3-ENSMUST00000051259.9-51409 | 0.010427 | 0.015685 | 0.002978 | 0.016302 | 0.007487 | 0.014416 | 0.013463 | 0.015373 | **0.012016** | 0.045333 | 0.050700 | 0.051680 | 0.048702 | 0.045201 | 0.053781 | 0.032459 | 0.052569 | **0.047553** |
| Itgae-ENSMUST00000006101.3-64270 | 0.051783 | 0.031873 | 0.030050 | 0.031604 | 0.061233 | 0.078479 | 0.032724 | 0.037934 | **0.044460** | 0.041010 | 0.033637 | 0.045456 | 0.045137 | 0.050831 | 0.052324 | 0.048002 | 0.063153 | **0.047444** |
| Irak1-ENSMUST00000114352.7-90942 | 0.243621 | 0.200852 | 0.183791 | 0.130040 | 0.274414 | 0.265045 | 0.580732 | 0.577187 | **0.306960** | 0.050891 | 0.047977 | 0.041811 | 0.041745 | 0.041212 | 0.035999 | 0.054399 | 0.053825 | **0.045982** |
| Il10-ENSMUST00000016673.5-4339 | 0.068377 | 0.056124 | 0.050569 | 0.050058 | 0.097847 | 0.069393 | 0.033154 | 0.045473 | **0.058874** | 0.056852 | 0.041999 | 0.049272 | 0.032821 | 0.055464 | 0.037538 | 0.045000 | 0.037388 | **0.044542** |
| Cxcr2-ENSMUST00000106899.3-2565 | 0.030856 | 0.041499 | 0.035589 | 0.021817 | 0.013581 | 0.015823 | 0.016921 | 0.015890 | **0.023997** | 0.097605 | 0.065349 | 0.036871 | 0.018924 | 0.044746 | 0.026742 | 0.035588 | 0.030273 | **0.044512** |
| Cd163-ENSMUST00000032234.4-37639 | 0.319372 | 0.333621 | 0.690840 | 0.587520 | 0.402792 | 0.443928 | 0.111689 | 0.166667 | **0.382054** | 0.069627 | 0.033518 | 0.072270 | 0.067384 | 0.012574 | 0.012088 | 0.053625 | 0.032727 | **0.044227** |
| Ighd-ENSMUST00000194162.5-70199 | 0.043477 | 0.033819 | 0.008471 | 0.030288 | 0.031136 | 0.032136 | 0.016017 | 0.015248 | **0.026324** | 0.030060 | 0.035448 | 0.043141 | 0.055808 | 0.025954 | 0.029143 | 0.050735 | 0.069120 | **0.042426** |
| Klra5-ENSMUST00000118060.7-38157 | 0.000000 | 0.006955 | 0.000000 | 0.013244 | 0.005284 | 0.007797 | 0.000000 | 0.000714 | **0.004249** | 0.015615 | 0.011834 | 0.044575 | 0.049427 | 0.067057 | 0.067572 | 0.028091 | 0.053674 | **0.042231** |
| Il6-ENSMUST00000199183.4-27154 | 0.003834 | 0.008652 | 0.021361 | 0.009390 | 0.001197 | 0.006042 | 0.005380 | 0.009062 | **0.008115** | 0.094374 | 0.090586 | 0.023975 | 0.026136 | 0.020787 | 0.018029 | 0.028777 | 0.020378 | **0.040380** |
| Fam65b-ENSMUST00000091694.9-71060 | 0.062404 | 0.059113 | 0.106601 | 0.070258 | 0.114576 | 0.099759 | 0.042162 | 0.044696 | **0.074946** | 0.034078 | 0.039544 | 0.035094 | 0.038956 | 0.046300 | 0.038338 | 0.035542 | 0.045136 | **0.039123** |
| Ikzf2-ENSMUST00000027146.8-2426 | 0.019352 | 0.023283 | 0.022185 | 0.024812 | 0.043047 | 0.059521 | 0.009665 | 0.017513 | **0.027422** | 0.026091 | 0.029708 | 0.024220 | 0.029191 | 0.040702 | 0.042345 | 0.039077 | 0.058577 | **0.036239** |
| Cd79a-ENSMUST00000003469.7-40721 | 0.040832 | 0.030310 | 0.006885 | 0.022869 | 0.013040 | 0.014015 | 0.025104 | 0.025863 | **0.022365** | 0.029939 | 0.039388 | 0.028999 | 0.042023 | 0.018503 | 0.018592 | 0.038345 | 0.053832 | **0.033702** |
| Fam129c-ENSMUST00000143662.7-50383 | 0.023662 | 0.032051 | 0.022016 | 0.031747 | 0.015930 | 0.024888 | 0.034524 | 0.025818 | **0.026329** | 0.022313 | 0.019033 | 0.029493 | 0.031436 | 0.036320 | 0.045119 | 0.032441 | 0.031086 | **0.030905** |
| Bach2-ENSMUST00000108180.8-20833 | 0.065689 | 0.098977 | 0.113561 | 0.104284 | 0.037708 | 0.044696 | 0.019767 | 0.023456 | **0.063517** | 0.027709 | 0.033217 | 0.030591 | 0.019034 | 0.041670 | 0.046364 | 0.022977 | 0.024277 | **0.030730** |
| Gzmm-ENSMUST00000020549.3-59427 | 0.012399 | 0.007562 | 0.000000 | 0.022293 | 0.004534 | 0.015265 | 0.006004 | 0.008853 | **0.009614** | 0.018640 | 0.011442 | 0.023329 | 0.032860 | 0.037613 | 0.041760 | 0.022268 | 0.046628 | **0.029317** |
| Iglc3-ENSMUST00000200235.1-79750 | 0.024339 | 0.020571 | 0.008639 | 0.015187 | 0.008162 | 0.009492 | 0.011743 | 0.009006 | **0.013392** | 0.020041 | 0.025234 | 0.031950 | 0.031566 | 0.015537 | 0.016825 | 0.032608 | 0.045984 | **0.027468** |
| Zbtb16-ENSMUST00000093852.3-54243 | 0.025785 | 0.032824 | 0.042991 | 0.028600 | 0.029177 | 0.025255 | 0.009567 | 0.006195 | **0.025049** | 0.026812 | 0.034306 | 0.018702 | 0.018478 | 0.040128 | 0.029541 | 0.024047 | 0.022178 | **0.026774** |
| Btla-ENSMUST00000063654.4-80651 | 0.026156 | 0.028931 | 0.034204 | 0.028686 | 0.045065 | 0.056573 | 0.024493 | 0.022047 | **0.033269** | 0.013696 | 0.019425 | 0.024785 | 0.023721 | 0.025364 | 0.031890 | 0.029144 | 0.036781 | **0.025601** |
| Lta-ENSMUST00000025266.5-83442 | 0.007582 | 0.010807 | 0.011113 | 0.021941 | 0.009577 | 0.014218 | 0.042590 | 0.041597 | **0.019928** | 0.018714 | 0.017549 | 0.026599 | 0.015099 | 0.026685 | 0.032827 | 0.020520 | 0.031438 | **0.023679** |
| Foxp3-ENSMUST00000115740.7-89338 | 0.018556 | 0.019065 | 0.021144 | 0.009175 | 0.048698 | 0.069698 | 0.014578 | 0.019283 | **0.027525** | 0.019217 | 0.021926 | 0.012309 | 0.016454 | 0.031978 | 0.032830 | 0.015539 | 0.027462 | **0.022214** |
| Cxcr5-ENSMUST00000179828.7-54074 | 0.015135 | 0.009748 | 0.024565 | 0.007140 | 0.005895 | 0.012168 | 0.015870 | 0.015088 | **0.013201** | 0.019598 | 0.019026 | 0.014260 | 0.019240 | 0.026355 | 0.028298 | 0.018350 | 0.028156 | **0.021660** |
| Tnfrsf8-ENSMUST00000030339.12-25454 | 0.026487 | 0.016495 | 0.031701 | 0.026732 | 0.027772 | 0.037313 | 0.029021 | 0.027934 | **0.027932** | 0.024976 | 0.024208 | 0.015998 | 0.012872 | 0.026961 | 0.020418 | 0.014494 | 0.019373 | **0.019912** |
| Il12b-ENSMUST00000102796.9-62286 | 0.022302 | 0.016241 | 0.052098 | 0.014257 | 0.034771 | 0.024927 | 0.007686 | 0.012088 | **0.023046** | 0.012076 | 0.018864 | 0.015513 | 0.011681 | 0.020329 | 0.020242 | 0.022219 | 0.036208 | **0.019641** |
| Klra17-ENSMUST00000122219.1-38152 | 0.020096 | 0.012261 | 0.045739 | 0.012562 | 0.028505 | 0.017007 | 0.009337 | 0.006641 | **0.019018** | 0.039394 | 0.018109 | 0.014961 | 0.008777 | 0.020649 | 0.010196 | 0.027405 | 0.010726 | **0.018777** |
| Fcrla-ENSMUST00000046322.13-5936 | 0.015222 | 0.018912 | 0.018146 | 0.007730 | 0.015082 | 0.021647 | 0.028107 | 0.027669 | **0.019064** | 0.026477 | 0.019526 | 0.018932 | 0.012751 | 0.013319 | 0.011242 | 0.025259 | 0.021409 | **0.018614** |
| F5-ENSMUST00000086040.5-5653 | 0.014826 | 0.025550 | 0.023881 | 0.030459 | 0.014231 | 0.029694 | 0.050175 | 0.065277 | **0.031762** | 0.040064 | 0.043488 | 0.010424 | 0.008385 | 0.012122 | 0.007307 | 0.008754 | 0.016141 | **0.018336** |
| Csf2-ENSMUST00000019060.5-62907 | 0.012267 | 0.003510 | 0.005193 | 0.007184 | 0.004501 | 0.008212 | 0.003706 | 0.003247 | **0.005977** | 0.006859 | 0.012477 | 0.016213 | 0.011185 | 0.013563 | 0.020044 | 0.019850 | 0.031068 | **0.016407** |
| Klra6-ENSMUST00000074056.2-38161 | 0.011232 | 0.005165 | 0.002911 | 0.011708 | 0.001052 | 0.008241 | 0.006305 | 0.006859 | **0.006684** | 0.013088 | 0.009671 | 0.007329 | 0.006153 | 0.015350 | 0.016837 | 0.022626 | 0.037925 | **0.016122** |
| Cd19-ENSMUST00000206325.1-46361 | 0.012150 | 0.013143 | 0.003956 | 0.010973 | 0.004412 | 0.005272 | 0.015980 | 0.013508 | **0.009924** | 0.014020 | 0.017156 | 0.014621 | 0.016270 | 0.006221 | 0.007796 | 0.014281 | 0.033061 | **0.015428** |
| Il12a-ENSMUST00000107816.3-16325 | 0.018275 | 0.018229 | 0.013961 | 0.012943 | 0.001697 | 0.004459 | 0.018018 | 0.016459 | **0.013005** | 0.011456 | 0.016971 | 0.013851 | 0.024893 | 0.013375 | 0.016380 | 0.008049 | 0.014581 | **0.014944** |
| Mzb1-ENSMUST00000025211.4-85784 | 0.014378 | 0.011764 | 0.000000 | 0.015401 | 0.003364 | 0.004144 | 0.007360 | 0.009859 | **0.008284** | 0.008761 | 0.015011 | 0.012711 | 0.013215 | 0.011387 | 0.011836 | 0.014678 | 0.020526 | **0.013516** |

| | | | | | | | | | | | | | | | | | | |
|---|---|---|---|---|---|---|---|---|---|---|---|---|---|---|---|---|---|---|
| Cpa3-ENSMUST00000001921.2-14937 | 0.010568 | 0.012797 | 0.000000 | 0.023485 | 0.018395 | 0.021298 | 0.005277 | 0.009253 | **0.012634** | 0.000441 | 0.000000 | 0.012658 | 0.010094 | 0.021724 | 0.022355 | 0.009814 | 0.024133 | **0.012652** |
| Gzmk-ENSMUST00000122399.7-73201 | 0.002991 | 0.003399 | 0.003227 | 0.007620 | 0.018220 | 0.025725 | 0.023547 | 0.021224 | **0.013244** | 0.003759 | 0.001887 | 0.006656 | 0.006274 | 0.022972 | 0.024116 | 0.016924 | 0.016823 | **0.012426** |
| Pou2af1-ENSMUST00000034554.7-54351 | 0.007515 | 0.006999 | 0.000000 | 0.003023 | 0.002909 | 0.005535 | 0.003122 | 0.005678 | **0.004348** | 0.009883 | 0.010619 | 0.012297 | 0.014966 | 0.006625 | 0.004321 | 0.012982 | 0.026094 | **0.012223** |
| Tnfrsf25-ENSMUST00000035275.7-25878 | 0.005306 | 0.018236 | 0.005156 | 0.009783 | 0.018117 | 0.008635 | 0.009349 | 0.008064 | **0.010331** | 0.005757 | 0.008319 | 0.008918 | 0.011884 | 0.014283 | 0.020397 | 0.010621 | 0.017082 | **0.012158** |
| Fcer2a-ENSMUST00000005678.5-48045 | 0.012553 | 0.009036 | 0.000000 | 0.005255 | 0.002858 | 0.005221 | 0.009504 | 0.010209 | **0.006829** | 0.013167 | 0.011513 | 0.012078 | 0.015784 | 0.004676 | 0.004838 | 0.011792 | 0.017056 | **0.011363** |
| Bcl11a-ENSMUST00000109514.7-61965 | 0.026853 | 0.015414 | 0.008041 | 0.015013 | 0.012633 | 0.025539 | 0.023676 | 0.022280 | **0.018681** | 0.015013 | 0.015655 | 0.013244 | 0.006725 | 0.007843 | 0.012140 | 0.008192 | 0.008352 | **0.010895** |
| Cd200-ENSMUST00000163230.7-80654 | 0.005447 | 0.010818 | 0.010581 | 0.014249 | 0.022658 | 0.033962 | 0.014461 | 0.014540 | **0.015840** | 0.009274 | 0.012316 | 0.007019 | 0.005327 | 0.011935 | 0.011186 | 0.009641 | 0.020194 | **0.010861** |
| Kcne3-ENSMUST00000208260.1-44727 | 0.060357 | 0.056861 | 0.061257 | 0.039682 | 0.048458 | 0.045058 | 0.052421 | 0.056004 | **0.052512** | 0.011696 | 0.008192 | 0.018308 | 0.012918 | 0.011500 | 0.007289 | 0.009083 | 0.004690 | **0.010459** |
| Ccr6-ENSMUST00000180103.1-81759 | 0.022670 | 0.025849 | 0.017709 | 0.014719 | 0.013350 | 0.018922 | 0.019683 | 0.019196 | **0.019012** | 0.007809 | 0.018236 | 0.008652 | 0.007447 | 0.006411 | 0.008123 | 0.006837 | 0.014869 | **0.009798** |
| Trat1-ENSMUST00000170861.1-80711 | 0.003491 | 0.003031 | 0.007386 | 0.006795 | 0.013108 | 0.015789 | 0.007724 | 0.005201 | **0.007816** | 0.005673 | 0.010481 | 0.005271 | 0.006989 | 0.010696 | 0.013425 | 0.008175 | 0.013819 | **0.009316** |
| Rora-ENSMUST00000034766.13-55012 | 0.000000 | 0.015103 | 0.010797 | 0.010971 | 0.018280 | 0.021751 | 0.009715 | 0.009473 | **0.012011** | 0.009529 | 0.009062 | 0.004799 | 0.005487 | 0.010052 | 0.009921 | 0.008439 | 0.011353 | **0.008580** |
| Blk-ENSMUST00000014597.3-75650 | 0.008363 | 0.007796 | 0.000000 | 0.011840 | 0.001702 | 0.003894 | 0.007457 | 0.008495 | **0.006193** | 0.006116 | 0.009345 | 0.004610 | 0.008596 | 0.003560 | 0.006964 | 0.008746 | 0.018548 | **0.008311** |
| Cxcr4-ENSMUST00000052172.6-4256 | 0.092028 | 0.078154 | 0.126628 | 0.079866 | 0.015878 | 0.009808 | 0.221680 | 0.192557 | **0.102075** | 0.009596 | 0.011575 | 0.009192 | 0.006420 | 0.004109 | 0.003769 | 0.010327 | 0.007839 | **0.007853** |
| Lrrc32-ENSMUST00000205956.1-44593 | 0.003218 | 0.004052 | 0.006577 | 0.007111 | 0.011702 | 0.015477 | 0.007247 | 0.010039 | **0.008178** | 0.007162 | 0.009992 | 0.003990 | 0.006084 | 0.007846 | 0.009060 | 0.003892 | 0.009880 | **0.007238** |
| Irf4-ENSMUST00000021784.8-71193 | 0.023829 | 0.012970 | 0.028780 | 0.045465 | 0.022439 | 0.028910 | 0.005337 | 0.009749 | **0.022185** | 0.013655 | 0.006014 | 0.006800 | 0.003711 | 0.005273 | 0.005154 | 0.004077 | 0.009840 | **0.006815** |
| Il33-ENSMUST00000120388.8-88332 | 0.003627 | 0.007847 | 0.007801 | 0.002180 | 0.002376 | 0.003068 | 0.007799 | 0.004461 | **0.004895** | 0.009548 | 0.009590 | 0.001914 | 0.003630 | 0.003921 | 0.003624 | 0.006555 | 0.004772 | **0.005444** |
| Fcer1a-ENSMUST00000049706.10-6210 | 0.000000 | 0.002471 | 0.003460 | 0.004230 | 0.005493 | 0.007155 | 0.003740 | 0.001838 | **0.003548** | 0.000916 | 0.000000 | 0.008198 | 0.003849 | 0.008887 | 0.008564 | 0.004009 | 0.008483 | **0.005363** |
| Slamf1-ENSMUST00000015460.4-6096 | 0.005632 | 0.009584 | 0.009819 | 0.005340 | 0.013478 | 0.017420 | 0.008377 | 0.010139 | **0.009974** | 0.002830 | 0.001505 | 0.002320 | 0.005122 | 0.010759 | 0.007718 | 0.007124 | 0.005384 | **0.005345** |
| Klrc3-ENSMUST00000071149.6-38132 | 0.000000 | 0.001954 | 0.000000 | 0.002471 | 0.003126 | 0.005717 | 0.000000 | 0.000885 | **0.001769** | 0.004589 | 0.006254 | 0.002893 | 0.003212 | 0.003754 | 0.005759 | 0.003568 | 0.012303 | **0.005292** |
| Ccr9-ENSMUST00000163559.7-57573 | 0.002530 | 0.001633 | 0.000000 | 0.001823 | 0.002078 | 0.002846 | 0.001012 | 0.003307 | **0.001904** | 0.003229 | 0.003641 | 0.006038 | 0.004289 | 0.004268 | 0.005056 | 0.005131 | 0.009581 | **0.005154** |
| Tnfrsf13c-ENSMUST00000089161.8-78067 | 0.000000 | 0.006615 | 0.000000 | 0.011629 | 0.006976 | 0.006940 | 0.009995 | 0.008021 | **0.006272** | 0.002257 | 0.005693 | 0.005838 | 0.008239 | 0.003129 | 0.002974 | 0.004531 | 0.005696 | **0.004795** |
| Ccr8-ENSMUST00000048777.2-57425 | 0.004640 | 0.002900 | 0.011568 | 0.008921 | 0.013913 | 0.029733 | 0.002700 | 0.003877 | **0.009782** | 0.003325 | 0.003917 | 0.002645 | 0.003044 | 0.004567 | 0.003850 | 0.004468 | 0.008099 | **0.004239** |
| Ms4a1-ENSMUST00000169159.2-87812 | 0.005395 | 0.000000 | 0.015592 | 0.022728 | 0.001637 | 0.005471 | 0.006824 | 0.003956 | **0.007700** | 0.003267 | 0.004197 | 0.002573 | 0.006511 | 0.001958 | 0.003505 | 0.004314 | 0.006853 | **0.004147** |
| Cr2-ENSMUST00000195120.5-7084 | 0.002912 | 0.002097 | 0.000000 | 0.009963 | 0.001272 | 0.000765 | 0.001974 | 0.002034 | **0.002627** | 0.002983 | 0.003797 | 0.002644 | 0.005014 | 0.001133 | 0.000670 | 0.005384 | 0.007811 | **0.003679** |
| Ccr3-ENSMUST00000039171.7-57601 | 0.008254 | 0.006168 | 0.014525 | 0.006187 | 0.011019 | 0.014207 | 0.008806 | 0.005935 | **0.009388** | 0.002786 | 0.002119 | 0.006164 | 0.002601 | 0.002444 | 0.001839 | 0.001950 | 0.003362 | **0.002908** |
| Cxcl11-ENSMUST00000077820.5-29466 | 0.003275 | 0.001175 | 0.003504 | 0.005483 | 0.009130 | 0.001283 | 0.004622 | 0.003091 | **0.003945** | 0.002297 | 0.001538 | 0.003598 | 0.000858 | 0.002857 | 0.003622 | 0.004565 | 0.003333 | **0.002834** |
| Cxcr1-ENSMUST00000190313.1-2566 | 0.019359 | 0.006903 | 0.014686 | 0.010630 | 0.002626 | 0.002107 | 0.003892 | 0.007091 | **0.008412** | 0.007553 | 0.002377 | 0.002088 | 0.001186 | 0.001491 | 0.001350 | 0.002559 | 0.003226 | **0.002729** |
| Ccr10-ENSMUST00000062759.3-66240 | 0.008648 | 0.007202 | 0.023410 | 0.011833 | 0.006103 | 0.005296 | 0.005133 | 0.003710 | **0.008917** | 0.004571 | 0.001367 | 0.003173 | 0.002827 | 0.002147 | 0.001036 | 0.003618 | 0.002517 | **0.002657** |
| Vpreb3-ENSMUST00000000926.2-59172 | 0.004667 | 0.002786 | 0.000000 | 0.003377 | 0.000860 | 0.001387 | 0.003741 | 0.002870 | **0.002461** | 0.004795 | 0.003552 | 0.000471 | 0.004102 | 0.000363 | 0.000198 | 0.003451 | 0.004252 | **0.002648** |
| Ebf1-ENSMUST00000081265.11-62371 | 0.000000 | 0.000000 | 0.006869 | 0.001375 | 0.002468 | 0.003615 | 0.002528 | 0.003054 | **0.002489** | 0.001954 | 0.001280 | 0.001905 | 0.004915 | 0.000770 | 0.000227 | 0.002835 | 0.005878 | **0.002470** |
| Klrb1-ENSMUST00000112110.3-38019 | 0.006651 | 0.000000 | 0.000000 | 0.011044 | 0.000796 | 0.000947 | 0.000000 | 0.000095 | **0.002442** | 0.000000 | 0.000781 | 0.003180 | 0.002658 | 0.000543 | 0.001898 | 0.001918 | 0.006393 | **0.002171** |
| Elane-ENSMUST00000046091.5-59482 | 0.000000 | 0.000792 | 0.000000 | 0.001793 | 0.001032 | 0.002103 | 0.003881 | 0.004859 | **0.001808** | 0.004956 | 0.001428 | 0.001882 | 0.001354 | 0.000414 | 0.001288 | 0.001313 | 0.001916 | **0.001819** |
| Cd1d2-ENSMUST00000194208.5-16688 | 0.000000 | 0.000000 | 0.010142 | 0.002471 | 0.000000 | 0.001893 | 0.000889 | 0.001269 | **0.002083** | 0.002044 | 0.001409 | 0.000687 | 0.000535 | 0.002117 | 0.001676 | 0.000579 | 0.001721 | **0.001346** |
| Pax5-ENSMUST00000014174.13-21312 | 0.000000 | 0.001555 | 0.002849 | 0.003597 | 0.000000 | 0.000149 | 0.000000 | 0.002332 | **0.001310** | 0.001403 | 0.000000 | 0.001712 | 0.002117 | 0.000371 | 0.000389 | 0.001397 | 0.002991 | **0.001298** |
| Il5-ENSMUST00000048605.2-62861 | 0.018184 | 0.001330 | 0.006860 | 0.002573 | 0.003053 | 0.000914 | 0.000000 | 0.000092 | **0.004126** | 0.002147 | 0.002898 | 0.001390 | 0.000855 | 0.000743 | 0.000617 | 0.000207 | 0.000000 | **0.001107** |
| Ighg2b-ENSMUST00000103418.2-70177 | 0.000000 | 0.000000 | 0.000000 | 0.000000 | 0.000000 | 0.000927 | 0.000000 | 0.000983 | **0.000239** | 0.001323 | 0.000000 | 0.002029 | 0.000000 | 0.001127 | 0.001460 | 0.000761 | 0.001477 | **0.001022** |
| Jchain-ENSMUST00000087033.5-29277 | 0.000000 | 0.001599 | 0.000000 | 0.000000 | 0.000943 | 0.004590 | 0.000000 | 0.000649 | **0.000973** | 0.002400 | 0.001464 | 0.000000 | 0.001309 | 0.001052 | 0.000942 | 0.000546 | 0.000000 | **0.000964** |
| Ighg3-ENSMUST00000103423.2-70185 | 0.000000 | 0.003525 | 0.000000 | 0.000000 | 0.001161 | 0.001678 | 0.000000 | 0.000252 | **0.000827** | 0.001007 | 0.000000 | 0.000000 | 0.000317 | 0.001047 | 0.001608 | 0.000821 | 0.002219 | **0.000877** |
| Il17a-ENSMUST00000027061.4-791 | 0.000000 | 0.003249 | 0.000000 | 0.000000 | 0.000000 | 0.000000 | 0.000000 | 0.000667 | **0.000489** | 0.000000 | 0.001162 | 0.000000 | 0.000997 | 0.001578 | 0.001152 | 0.001994 | 0.000000 | **0.000860** |
| Rorc-ENSMUST00000029795.9-17465 | 0.000000 | 0.000895 | 0.000000 | 0.002296 | 0.000000 | 0.000000 | 0.002591 | 0.000914 | **0.000837** | 0.000487 | 0.000000 | 0.000722 | 0.000920 | 0.001019 | 0.001137 | 0.001310 | 0.000565 | **0.000770** |
| Alas2-ENSMUST00000066337.12-92827 | 0.000000 | 0.001266 | 0.000000 | 0.000000 | 0.000000 | 0.001015 | 0.000706 | 0.002893 | **0.000735** | 0.000000 | 0.000672 | 0.000000 | 0.000295 | 0.000802 | 0.001341 | 0.002208 | 0.000407 | **0.000716** |
| Il23r-ENSMUST00000118364.1-35326 | 0.000000 | 0.001554 | 0.004425 | 0.005357 | 0.000000 | 0.000000 | 0.000660 | 0.000000 | **0.001499** | 0.001589 | 0.002236 | 0.000000 | 0.000000 | 0.000000 | 0.000647 | 0.000415 | 0.000791 | **0.000710** |
| Il17f-ENSMUST00000039046.9-792 | 0.003810 | 0.004879 | 0.000000 | 0.000000 | 0.000000 | 0.002443 | 0.000000 | 0.000119 | **0.001407** | 0.000718 | 0.000000 | 0.000456 | 0.000542 | 0.001242 | 0.001539 | 0.000659 | 0.000405 | **0.000695** |
| Cxcl13-ENSMUST00000023840.6-29610 | 0.000000 | 0.000000 | 0.000000 | 0.000000 | 0.000000 | 0.001198 | 0.000000 | 0.000392 | **0.000199** | 0.001533 | 0.000000 | 0.000511 | 0.000000 | 0.000150 | 0.000392 | 0.000000 | 0.001369 | **0.000494** |
| Il2-ENSMUST00000029275.5-15438 | 0.000000 | 0.000000 | 0.000000 | 0.000000 | 0.000873 | 0.000000 | 0.000000 | 0.000424 | **0.000162** | 0.001525 | 0.000000 | 0.000000 | 0.001104 | 0.000000 | 0.000237 | 0.000189 | 0.000789 | **0.000481** |
| Ighg2c-ENSMUST00000195192.1-70175 | 0.000000 | 0.000000 | 0.000000 | 0.000000 | 0.001101 | 0.000663 | 0.000000 | 0.000315 | **0.000260** | 0.000851 | 0.000000 | 0.000000 | 0.000000 | 0.000114 | 0.001703 | 0.000298 | 0.000743 | **0.000464** |
| Ccl22-ENSMUST00000034231.3-51368 | 0.000000 | 0.002374 | 0.000000 | 0.001697 | 0.000978 | 0.002273 | 0.001594 | 0.000754 | **0.001209** | 0.000515 | 0.000447 | 0.000637 | 0.000672 | 0.000448 | 0.000164 | 0.000484 | 0.000000 | **0.000421** |
| Il22-ENSMUST00000096691.4-60647 | 0.000000 | 0.000000 | 0.000000 | 0.000000 | 0.000000 | 0.000000 | 0.000000 | 0.000000 | **0.000000** | 0.000000 | 0.000000 | 0.000000 | 0.000887 | 0.000945 | 0.000673 | 0.000250 | 0.000369 | **0.000390** |
| Il4-ENSMUST00000000889.6-62853 | 0.000000 | 0.000000 | 0.000000 | 0.000000 | 0.000000 | 0.000392 | 0.000000 | 0.000000 | **0.000049** | 0.000744 | 0.000687 | 0.000000 | 0.000234 | 0.000394 | 0.000532 | 0.000302 | 0.000000 | **0.000362** |
| Il13-ENSMUST00000020650.1-62855 | 0.000000 | 0.000000 | 0.002909 | 0.000000 | 0.001211 | 0.001551 | 0.000000 | 0.000794 | **0.000808** | 0.000000 | 0.001335 | 0.000000 | 0.000000 | 0.000873 | 0.000205 | 0.000207 | 0.000000 | **0.000328** |
| Ccl19-ENSMUST00000102957.5-21191 | 0.003089 | 0.000000 | 0.000000 | 0.000000 | 0.004198 | 0.001102 | 0.000000 | 0.000268 | **0.001082** | 0.000000 | 0.000706 | 0.000985 | 0.000260 | 0.000104 | 0.000454 | 0.000000 | 0.000000 | **0.000314** |
| Il21-ENSMUST00000029273.7-15440 | 0.009299 | 0.000000 | 0.000000 | 0.000000 | 0.001217 | 0.001310 | 0.000000 | 0.000000 | **0.001478** | 0.000000 | 0.000000 | 0.000000 | 0.000000 | 0.000656 | 0.000457 | 0.000000 | 0.001023 | **0.000267** |
| Ifna1-ENSMUST00000094972.1-22502 | 0.002486 | 0.000000 | 0.000000 | 0.000000 | 0.000000 | 0.000533 | 0.000000 | 0.000000 | **0.000377** | 0.000000 | 0.001184 | 0.000000 | 0.000177 | 0.000395 | 0.000307 | 0.000000 | 0.000000 | **0.000258** |
| Tnfrsf17-ENSMUST00000023140.5-79339 | 0.000000 | 0.000000 | 0.000000 | 0.000000 | 0.000000 | 0.001279 | 0.000764 | 0.000000 | **0.000255** | 0.000971 | 0.000000 | 0.000000 | 0.000000 | 0.000088 | 0.000000 | 0.000320 | 0.000428 | **0.000226** |
| Ighe-ENSMUST00000137336.2-70173 | 0.000000 | 0.000000 | 0.000000 | 0.001907 | 0.000000 | 0.000296 | 0.000000 | 0.000000 | **0.000275** | 0.000000 | 0.000000 | 0.000517 | 0.000321 | 0.000196 | 0.000198 | 0.000000 | 0.000000 | **0.000154** |
| Ccl20-ENSMUST00000113437.8-3018 | 0.000000 | 0.000000 | 0.000000 | 0.000000 | 0.000000 | 0.000000 | 0.000000 | 0.000000 | **0.000000** | 0.000000 | 0.000000 | 0.000000 | 0.000175 | 0.000000 | 0.000389 | 0.000160 | 0.000000 | **0.000090** |
| Il9-ENSMUST00000022019.3-71893 | 0.000000 | 0.000000 | 0.000000 | 0.000000 | 0.000000 | 0.000000 | 0.000000 | 0.000000 | **0.000000** | 0.000000 | 0.000000 | 0.000000 | 0.000000 | 0.000000 | 0.000000 | 0.000000 | 0.000570 | **0.000071** |

| | | | | | | | | | | | | | | | | | | |
|---|---|---|---|---|---|---|---|---|---|---|---|---|---|---|---|---|---|---|
| Il25-ENSMUST00000037863.5-75244 | 0.000000 | 0.000000 | 0.000000 | 0.000000 | 0.000000 | 0.000000 | 0.000000 | 0.000000 | **0.000000** | 0.000000 | 0.000000 | 0.000000 | 0.000000 | 0.000112 | 0.000000 | 0.000000 | 0.000000 | **0.000014** |
| Ighg1-ENSMUST00000103420.2-70184 | 0.000000 | 0.000000 | 0.000000 | 0.000000 | 0.000000 | 0.000000 | 0.000000 | 0.000000 | **0.000000** | 0.000000 | 0.000000 | 0.000000 | 0.000000 | 0.000000 | 0.000000 | 0.000000 | 0.000000 | **0.000000** |
