## Supplementary Table S4 for "Reprogramming tumour-associated macrophages from immune suppressive to inflammatory state by Checkpoint kinase 1 inhibitor combination treatment"

**Info**

INFO: Top 20 cluster-specific macrophage markers (genes in ≤2 clusters)

Top20Markers table columns:
cluster : Cluster ID (0-11) from RNA UMAP

| cluster | gene | avg_log2FC | FC | pct.1 | pct.2 | p_val | p_val_adj | |
|---|---|---|---|---|---|---|---|---|
| 0 | Chil3-ENSMUST00000063062.8-18365 | 1.47 | 2.78 | 0.33 | 0.201 | 5.8626E-278 | 2.3274E-275 | |
| 0 | Arg1-ENSMUST00000020161.8-58127 | 0.85 | 1.80 | 0.943 | 0.872 | 0 | 0 | 0 suppressive |
| 0 | Ccl2-ENSMUST00000000193.5-64921 | 0.68 | 1.60 | 0.988 | 0.949 | 0 | 0 | |
| 0 | Nt5e-ENSMUST00000034992.7-55687 | 0.66 | 1.58 | 0.406 | 0.31 | 1.5017E-164 | 5.9619E-162 | |
| 1 | Mmp12-ENSMUST00000005950.11-52922 | 1.28 | 2.43 | 0.287 | 0.142 | 5.3498E-267 | 2.1239E-264 | 1 suppressive |
| 1 | Cd38-ENSMUST00000030964.5-27997 | 0.80 | 1.75 | 0.453 | 0.26 | 2.1981E-305 | 8.7265E-303 | |
| 2 | Cd163-ENSMUST00000032234.4-37639 | 3.50 | 11.33 | 0.252 | 0.02 | 0 | 0 | |
| 2 | Fcna-ENSMUST00000028307.8-8019 | 3.38 | 10.39 | 0.644 | 0.124 | 0 | 0 | |
| 2 | F13a1-ENSMUST00000164727.7-71368 | 2.25 | 4.75 | 0.847 | 0.389 | 0 | 0 | |
| 2 | Cd33-ENSMUST00000205503.1-42139 | 1.74 | 3.34 | 0.652 | 0.23 | 0 | 0 | |
| 2 | Maf-ENSMUST00000109104.1-52210 | 1.60 | 3.03 | 0.874 | 0.419 | 0 | 0 | |
| 2 | Tcf4-ENSMUST00000114985.9-86599 | 1.55 | 2.93 | 0.266 | 0.059 | 0 | 0 | |
| 2 | Cd79b-ENSMUST00000167143.1-66704 | 1.41 | 2.66 | 0.254 | 0.071 | 0 | 0 | |
| 2 | Cx3cr1-ENSMUST00000064165.3-57415 | 1.41 | 2.66 | 0.744 | 0.312 | 0 | 0 | |
| 2 | Qpct-ENSMUST00000040789.4-84843 | 1.26 | 2.40 | 0.386 | 0.133 | 0 | 0 | |
| 2 | Hmox1-ENSMUST00000005548.7-50527 | 1.23 | 2.34 | 0.953 | 0.736 | 0 | 0 | |
| 2 | Lyz2-ENSMUST00000092163.7-60608 | 1.22 | 2.33 | 1 | 0.996 | 0 | 0 | |
| 2 | C5ar1-ENSMUST00000168818.1-40084 | 1.06 | 2.08 | 0.958 | 0.72 | 0 | 0 | |
| 2 | Nrp1-ENSMUST00000026917.9-52798 | 1.02 | 2.03 | 0.939 | 0.677 | 0 | 0 | |
| 2 | Mgst1-ENSMUST00000008684.10-38555 | 1.00 | 2.00 | 0.317 | 0.128 | 0 | 0 | |
| 2 | Cd63-ENSMUST00000105229.7-61044 | 0.99 | 1.99 | 0.964 | 0.786 | 0 | 0 | |
| 2 | Fcgr3-ENSMUST00000164044.7-5955 | 0.88 | 1.84 | 1 | 0.966 | 0 | 0 | Control |
| 2 | Adgre1-ENSMUST00000086763.11-84535 | 0.87 | 1.83 | 0.994 | 0.889 | 0 | 0 | suppressive |
| 2 | Tlr1-ENSMUST00000059349.5-28440 | 0.80 | 1.74 | 0.767 | 0.441 | 0 | 0 | |
| 2 | Ada-ENSMUST00000017841.3-13524 | 0.79 | 1.73 | 0.309 | 0.136 | 0 | 0 | |
| 2 | Ctsd-ENSMUST00000151120.8-47685 | 0.77 | 1.70 | 1 | 0.995 | 0 | 0 | |
| 2 | Irf7-ENSMUST00000106023.7-47502 | -2.66 | -6.34 | 0.584 | 0.962 | 0 | 0 | |
| 2 | H2-K1-ENSMUST00000025181.16-83137 | -1.44 | -2.72 | 1 | 1 | 0 | 0 | |
| 2 | Stat1-ENSMUST00000070968.13-1727 | -1.30 | -2.46 | 0.527 | 0.651 | 0 | 0 | |
| 2 | Jun-ENSMUST00000107094.1-22601 | -1.08 | -2.11 | 0.512 | 0.578 | 1.0929E-194 | 4.3388E-192 | |
| 2 | Lgals3-ENSMUST00000142734.7-74644 | -1.01 | -2.01 | 0.993 | 0.995 | 0 | 0 | |
| 2 | Btg1-ENSMUST00000038377.7-60318 | -0.98 | -1.97 | 0.98 | 0.979 | 0 | 0 | |
| 2 | Junb-ENSMUST00000064922.6-50930 | -0.96 | -1.94 | 0.942 | 0.959 | 0 | 0 | |
| 2 | Casp1-ENSMUST00000027015.5-52899 | -0.92 | -1.90 | 0.657 | 0.677 | 3.3034E-260 | 1.3115E-257 | |
| 2 | Fosb-ENSMUST00000003640.3-40308 | -0.91 | -1.88 | 0.765 | 0.812 | 0 | 0 | |
| 2 | Bcl2a1a-ENSMUST00000098485.3-55772 | -0.73 | -1.66 | 0.983 | 0.967 | 0 | 0 | |
| 2 | Hif1a-ENSMUST00000021530.7-68841 | -0.65 | -1.57 | 0.885 | 0.847 | 4.1445E-258 | 1.6454E-255 | |
| 2 | Dusp1-ENSMUST00000025025.6-82616 | -0.59 | -1.50 | 0.841 | 0.833 | 3.5524E-176 | 1.4103E-173 | |
| 3 | Aurkb-ENSMUST00000108666.7-63704 | 2.46 | 5.48 | 0.845 | 0.195 | 0 | 0 | |
| 3 | Pclaf-ENSMUST00000045802.6-54915 | 2.16 | 4.48 | 0.992 | 0.49 | 0 | 0 | |
| 3 | Ube2c-ENSMUST00000088248.12-13630 | 1.94 | 3.84 | 0.843 | 0.406 | 0 | 0 | |
| 3 | Tyms-ENSMUST00000026846.10-27158 | 1.79 | 3.45 | 0.898 | 0.366 | 0 | 0 | proliferative |
| 3 | Tnfsf13b-ENSMUST00000207792.1-48244 | 0.93 | 1.90 | 0.532 | 0.267 | 0 | 0 | |
| 3 | Mcm4-ENSMUST00000023353.3-79445 | 0.60 | 1.52 | 0.732 | 0.478 | 3.1827E-274 | 1.2635E-271 | |
| 3 | Ada-ENSMUST00000017841.3-13524 | 0.59 | 1.51 | 0.259 | 0.15 | 9.11009E-98 | 3.61671E-95 | |
| 3 | Mcm2-ENSMUST00000058011.7-36363 | 0.59 | 1.51 | 0.768 | 0.503 | 9.1175E-281 | 3.6196E-278 | |
| 4 | Cd72-ENSMUST00000098104.9-21266 | 0.82 | 1.76 | 0.932 | 0.944 | 0 | 0 | |
| 4 | Slc25a37-ENSMUST00000037064.4-75812 | 0.77 | 1.71 | 0.135 | 0.275 | 2.25543E-69 | 8.95407E-67 | |
| 4 | Nlrp3-ENSMUST00000101148.8-63198 | 0.74 | 1.68 | 0.381 | 0.615 | 6.75573E-15 | 2.68203E-12 | inactivating |
| 4 | Tnf-ENSMUST00000025263.14-83440 | 0.71 | 1.64 | 0.246 | 0.428 | 1.18095E-47 | 4.68839E-45 | |
| 4 | Glg1-ENSMUST00000169020.7-52084 | 0.70 | 1.62 | 0.259 | 0.52 | 1.71062E-91 | 6.79118E-89 | |
| 4 | Tyk2-ENSMUST00000001036.10-53294 | 0.67 | 1.59 | 0.344 | 0.631 | 6.88157E-52 | 2.73198E-49 | |
| 5 | Gzma-ENSMUST00000023897.5-73199 | 5.51 | 45.49 | 0.763 | 0.102 | 0 | 0 | |
| 5 | Klra21-NM-053151.1-234625 | 5.13 | 35.10 | 0.308 | 0.015 | 0 | 0 | |
| 5 | Klra3-ENSMUST00000111998.8-38183 | 4.95 | 30.86 | 0.275 | 0.013 | 0 | 0 | |
| 5 | Prf1-ENSMUST00000035419.5-58753 | 4.74 | 26.78 | 0.598 | 0.042 | 0 | 0 | |
| 5 | Xcl1-ENSMUST00000027860.7-5687 | 4.53 | 23.05 | 0.628 | 0.096 | 0 | 0 | |
| 5 | Klra7-ENSMUST00000049304.13-38174 | 4.34 | 20.27 | 0.646 | 0.067 | 0 | 0 | |

| | | | | | | | | |
|---|---|---|---|---|---|---|---|---|
| 5 | Eomes-ENSMUST00000035020.14-57330 | 4.29 | 19.55 | 0.495 | 0.033 | 0 | 0 | |
| 5 | Gzmb-ENSMUST00000015581.4-75403 | 4.28 | 19.48 | 0.915 | 0.19 | 0 | 0 | |
| 5 | Klra1-ENSMUST00000032288.5-38185 | 4.10 | 17.14 | 0.264 | 0.023 | 0 | 0 | |
| 5 | Txk-ENSMUST00000198464.2-28791 | 4.02 | 16.22 | 0.499 | 0.039 | 0 | 0 | NK interacting |
| 5 | Tnfrsf9-ENSMUST00000030808.9-25835 | 3.98 | 15.83 | 0.756 | 0.13 | 0 | 0 | |
| 5 | Cst7-ENSMUST00000089200.2-12723 | 3.96 | 15.53 | 0.645 | 0.085 | 0 | 0 | |
| 5 | Il2rb-ENSMUST00000089398.7-77753 | 3.85 | 14.46 | 0.953 | 0.216 | 0 | 0 | |
| 5 | Ctsw-ENSMUST00000025844.4-87236 | 3.65 | 12.58 | 0.848 | 0.117 | 0 | 0 | |
| 5 | Nkg7-ENSMUST00000070518.3-42115 | 3.65 | 12.56 | 0.959 | 0.174 | 0 | 0 | |
| 5 | Klrc1-ENSMUST00000032270.12-38138 | 3.54 | 11.60 | 0.581 | 0.065 | 0 | 0 | |
| 5 | Trbc1-ENSMUST00000192856.5-34295 | 3.43 | 10.79 | 0.914 | 0.146 | 0 | 0 | |
| 5 | Thy1-ENSMUST00000114840.1-53998 | 3.36 | 10.27 | 0.833 | 0.251 | 0 | 0 | |
| 5 | Tbx21-ENSMUST00000001484.2-65761 | 3.30 | 9.88 | 0.516 | 0.061 | 0 | 0 | |
| 5 | Tigit-BD-custom-chr16-43648113 | 3.24 | 9.42 | 0.424 | 0.043 | 0 | 0 | |
| 6 | Cd3d-ENSMUST00000034602.7-54114 | 6.14 | 70.75 | 0.659 | 0.018 | 0 | 0 | |
| 6 | Cd8b1-ENSMUST00000065248.8-35529 | 6.11 | 69.23 | 0.429 | 0.016 | 0 | 0 | |
| 6 | Cd8a-ENSMUST00000066747.13-35536 | 5.86 | 58.16 | 0.433 | 0.019 | 0 | 0 | |
| 6 | Cd3g-ENSMUST00000002101.11-54111 | 5.75 | 53.72 | 0.75 | 0.031 | 0 | 0 | |
| 6 | Trac-ENSMUST00000198398.4-75126 | 5.62 | 49.23 | 0.795 | 0.051 | 0 | 0 | |
| 6 | Cd6-ENSMUST00000080292.11-87771 | 5.60 | 48.66 | 0.474 | 0.018 | 0 | 0 | |
| 6 | Cd3e-ENSMUST00000102832.1-54115 | 5.46 | 44.10 | 0.438 | 0.015 | 0 | 0 | |
| 6 | Cd5-ENSMUST00000025571.7-87742 | 5.41 | 42.52 | 0.401 | 0.018 | 0 | 0 | |
| 6 | Cxcr6-ENSMUST00000049810.7-57554 | 5.32 | 40.01 | 0.334 | 0.018 | 0 | 0 | |
| 6 | Tnfrsf4-ENSMUST00000030952.5-26202 | 5.11 | 34.52 | 0.295 | 0.032 | 0 | 0 | |
| 6 | Lat-ENSMUST00000032997.7-46345 | 4.76 | 27.03 | 0.778 | 0.067 | 0 | 0 | |
| 6 | Trbc2-ENSMUST00000103299.2-34304 | 4.44 | 21.74 | 0.908 | 0.128 | 0 | 0 | T cell interacting |
| 6 | Icos-ENSMUST00000102827.3-2147 | 4.19 | 18.25 | 0.234 | 0.025 | 0 | 0 | |
| 6 | Pdcd1-ENSMUST00000027507.7-3686 | 4.40 | 21.08 | 0.22 | 0.017 | 0 | 0 | |
| 6 | Cd27-ENSMUST00000032486.12-37807 | 3.94 | 15.33 | 0.329 | 0.026 | 0 | 0 | |
| 6 | Cd247-ENSMUST00000005907.11-5752 | 3.85 | 14.42 | 0.585 | 0.06 | 0 | 0 | |
| 6 | Itk-ENSMUST00000020664.12-62412 | 3.78 | 13.75 | 0.472 | 0.049 | 0 | 0 | |
| 6 | Cxcr3-ENSMUST00000056614.6-91654 | 3.58 | 11.96 | 0.444 | 0.074 | 0 | 0 | |
| 6 | Tcf7-ENSMUST00000086844.9-62791 | 3.33 | 10.09 | 0.437 | 0.065 | 0 | 0 | |
| 6 | Gimap7-ENSMUST00000052503.7-34625 | 3.29 | 9.80 | 0.347 | 0.043 | 0 | 0 | |
| 6 | Cd7-ENSMUST00000026159.5-67713 | 3.15 | 8.90 | 0.269 | 0.045 | 0 | 0 | |
| 6 | Lck-ENSMUST00000102596.7-24289 | 3.14 | 8.81 | 0.814 | 0.14 | 0 | 0 | |
| 6 | Gzmb-ENSMUST00000015581.4-75403 | 1.60 | 3.04 | 0.628 | 0.227 | 0 | 0 | |
| 7 | Cd34-ENSMUST00000016638.7-7059 | 5.92 | 60.64 | 0.75 | 0.042 | 0 | 0 | |
| 7 | Ncam1-ENSMUST00000194252.5-54263 | 5.83 | 56.88 | 0.663 | 0.023 | 0 | 0 | |
| 7 | Trib2-ENSMUST00000020922.7-68015 | 4.87 | 29.16 | 0.411 | 0.023 | 0 | 0 | |
| 7 | Fosl1-ENSMUST00000025850.5-87228 | 4.75 | 26.90 | 0.488 | 0.051 | 0 | 0 | |
| 7 | Il1rl1-ENSMUST00000097772.9-1442 | 4.74 | 26.78 | 0.259 | 0.021 | 0 | 0 | |
| 7 | Mmp9-ENSMUST00000017881.2-13675 | 4.73 | 26.62 | 0.77 | 0.066 | 0 | 0 | |
| 7 | Csf1-ENSMUST00000014743.9-18457 | 4.66 | 25.35 | 0.656 | 0.061 | 0 | 0 | |
| 7 | Fscn1-ENSMUST00000031565.14-32540 | 4.23 | 18.72 | 0.913 | 0.121 | 0 | 0 | |
| 7 | Tgfb3-ENSMUST00000003687.6-69288 | 3.86 | 14.47 | 0.596 | 0.114 | 0 | 0 | |
| 7 | Thbd-ENSMUST00000099270.4-12649 | 2.79 | 6.91 | 0.267 | 0.078 | 2.3025E-197 | 9.141E-195 | |
| 7 | Tcf4-ENSMUST00000114985.9-86599 | 2.59 | 6.04 | 0.277 | 0.083 | 2.8107E-203 | 1.1158E-200 | |
| 7 | Lgals1-ENSMUST00000089377.5-77768 | 2.22 | 4.67 | 1 | 0.999 | 0 | 0 | |
| 7 | Cd63-ENSMUST00000105229.7-61044 | 2.18 | 4.52 | 0.992 | 0.805 | 0 | 0 | |
| 7 | Kdelr1-ENSMUST00000002855.13-42761 | 1.76 | 3.38 | 0.908 | 0.649 | 0 | 0 | Stromal interacting |
| 7 | Anxa5-ENSMUST00000029266.13-15400 | 1.62 | 3.07 | 0.996 | 0.967 | 0 | 0 | |
| 7 | Tyms-ENSMUST00000026846.10-27158 | 1.42 | 2.67 | 0.633 | 0.411 | 4.2159E-150 | 1.6737E-147 | |
| 7 | Lap3-ENSMUST00000046122.10-28077 | 1.35 | 2.55 | 0.843 | 0.613 | 6.3735E-267 | 2.5303E-264 | |
| 7 | Mapk8-ENSMUST00000111945.8-74249 | 1.35 | 2.55 | 0.335 | 0.17 | 2.05699E-88 | 8.16626E-86 | |
| 7 | Mcm4-ENSMUST00000023353.3-79445 | 1.23 | 2.35 | 0.677 | 0.498 | 1.6658E-110 | 6.6134E-108 | |
| 7 | Mcm2-ENSMUST00000058011.7-36363 | 1.21 | 2.31 | 0.709 | 0.523 | 8.5532E-121 | 3.3956E-118 | |
| 7 | Havcr2-ENSMUST00000020668.14-62428 | -0.92 | -1.89 | 0.429 | 0.655 | 2.84615E-87 | 1.12992E-84 | |
| 7 | Cd48-ENSMUST00000068584.6-6088 | -0.83 | -1.78 | 0.618 | 0.831 | 3.0146E-139 | 1.1968E-136 | |
| 7 | Bcl2a1a-ENSMUST00000098485.3-55772 | -0.82 | -1.76 | 0.839 | 0.973 | 3.605E-232 | 1.4312E-229 | |
| 7 | Itgb2-ENSMUST00000000299.13-59249 | -0.81 | -1.75 | 0.798 | 0.955 | 1.4577E-203 | 5.7873E-201 | |
| 7 | Cd52-ENSMUST00000000696.6-24674 | -0.80 | -1.74 | 0.871 | 0.99 | 1.4765E-248 | 5.8617E-246 | |
| 7 | Itgax-ENSMUST00000033053.7-46808 | -0.79 | -1.73 | 0.546 | 0.7 | 5.97339E-59 | 2.37144E-56 | |
| 7 | Icam1-ENSMUST00000086399.4-53284 | -0.75 | -1.69 | 0.512 | 0.689 | 3.72464E-70 | 1.47868E-67 | |
| 7 | Dusp1-ENSMUST00000025025.6-82616 | -0.65 | -1.57 | 0.725 | 0.838 | 1.3302E-62 | 5.28088E-60 | |

| | | | | | | | | |
|---|---|---|---|---|---|---|---|---|
| 8 | S100a9-ENSMUST00000117167.1-17275 | 8.72 | 422.29 | 0.599 | 0.003 | 0 | 0 | |
| 8 | Cxcr2-ENSMUST00000106899.3-2565 | 8.17 | 288.98 | 0.401 | 0.002 | 0 | 0 | |
| 8 | S100a8-ENSMUST00000069927.9-17272 | 7.48 | 178.30 | 0.938 | 0.063 | 0 | 0 | |
| 8 | Arg2-ENSMUST00000021550.6-68965 | 4.86 | 29.10 | 0.269 | 0.011 | 0 | 0 | |
| 8 | Il1r2-ENSMUST00000027243.12-1427 | 4.03 | 16.37 | 0.795 | 0.115 | 0 | 0 | |
| 8 | Mmp9-ENSMUST00000017881.2-13675 | 3.71 | 13.06 | 0.738 | 0.069 | 0 | 0 | |
| 8 | Cxcl2-ENSMUST00000200681.3-29365 | 2.81 | 7.00 | 0.922 | 0.538 | 0 | 0 | Tlr4 induced |
| 8 | Clec4d-ENSMUST00000032240.3-37600 | 2.13 | 4.37 | 0.746 | 0.325 | 0 | 0 | |
| 8 | Csf1-ENSMUST00000014743.9-18457 | 2.02 | 4.06 | 0.267 | 0.074 | 2.6948E-186 | 1.0698E-183 | |
| 8 | Tnfsf14-ENSMUST00000005976.6-84524 | 1.48 | 2.80 | 0.309 | 0.141 | 6.9772E-88 | 2.76995E-85 | |
| 8 | Ier3-ENSMUST00000003635.6-83532 | 1.23 | 2.35 | 0.95 | 0.783 | 2.6088E-237 | 1.0357E-234 | |
| 8 | Tlr4-ENSMUST00000048096.11-22134 | 0.90 | 1.87 | 0.348 | 0.231 | 6.28526E-33 | 2.49525E-30 | |
| 8 | Il15-ENSMUST00000209363.1-50703 | 0.70 | 1.63 | 0.295 | 0.212 | 2.11176E-17 | 8.38367E-15 | |
| 8 | Nlrp3-ENSMUST00000101148.8-63198 | 0.68 | 1.60 | 0.743 | 0.588 | 3.91134E-60 | 1.5528E-57 | |
| 9 | Flt3-ENSMUST00000049324.12-32846 | 6.64 | 99.62 | 0.714 | 0.015 | 0 | 0 | |
| 9 | Ccl17-ENSMUST00000034232.2-51374 | 6.48 | 89.33 | 0.332 | 0.011 | 0 | 0 | |
| 9 | Itgae-ENSMUST00000006101.3-64270 | 5.67 | 50.87 | 0.297 | 0.016 | 0 | 0 | |
| 9 | Btla-ENSMUST00000063654.4-80651 | 5.19 | 36.46 | 0.296 | 0.008 | 0 | 0 | |
| 9 | Fscn1-ENSMUST00000031565.14-32540 | 4.43 | 21.60 | 0.354 | 0.142 | 7.5715E-108 | 3.0059E-105 | |
| 9 | Dpp4-ENSMUST00000047812.7-9521 | 4.39 | 20.93 | 0.48 | 0.025 | 0 | 0 | |
| 9 | H2-Ob-ENSMUST00000095342.9-83196 | 3.61 | 12.24 | 0.327 | 0.03 | 0 | 0 | |
| 9 | Kit-ENSMUST00000005815.6-28962 | 3.27 | 9.64 | 0.277 | 0.022 | 0 | 0 | CD86+ inflammatory |
| 9 | Il1r2-ENSMUST00000027243.12-1427 | 2.77 | 6.84 | 0.544 | 0.127 | 0 | 0 | |
| 9 | Qpct-ENSMUST00000040789.4-84843 | 2.39 | 5.26 | 0.52 | 0.162 | 9.0864E-244 | 3.6073E-241 | |
| 9 | Tlr3-ENSMUST00000209772.1-49458 | 1.49 | 2.80 | 0.33 | 0.203 | 1.49874E-30 | 5.94999E-28 | |
| 9 | Cd40-ENSMUST00000017799.11-13689 | 1.30 | 2.47 | 0.371 | 0.2 | 9.55439E-47 | 3.79309E-44 | |
| 9 | Cd86-ENSMUST00000089620.10-80399 | 1.18 | 2.27 | 0.645 | 0.392 | 6.12496E-82 | 2.43161E-79 | |
| 9 | Il2ra-ENSMUST00000028111.5-7410 | 0.94 | 1.92 | 0.26 | 0.126 | 2.16618E-38 | 8.59972E-36 | |
| 9 | Bcl6-ENSMUST00000023151.5-80026 | 0.79 | 1.72 | 0.364 | 0.185 | 3.07368E-47 | 1.22025E-44 | |
| 9 | Nod2-ENSMUST00000118370.7-51093 | 0.69 | 1.61 | 0.276 | 0.156 | 2.05712E-25 | 8.16678E-23 | |
| 10 | Chil3-ENSMUST00000063062.8-18365 | 3.56 | 11.82 | 0.537 | 0.231 | 9.1966E-99 | 3.65105E-96 | |
| 10 | Mgst1-ENSMUST00000008684.10-38555 | 3.35 | 10.18 | 0.38 | 0.153 | 2.83568E-70 | 1.12576E-67 | |
| 10 | Nod2-ENSMUST00000118370.7-51093 | 2.55 | 5.84 | 0.309 | 0.157 | 3.48454E-33 | 1.38336E-30 | |
| 10 | Tnfrsf1b-ENSMUST00000030336.10-25450 | 2.49 | 5.61 | 0.815 | 0.589 | 1.416E-148 | 5.6217E-146 | |
| 10 | Il6ra-ENSMUST00000197679.4-17033 | 1.36 | 2.57 | 0.648 | 0.659 | 7.2222E-33 | 2.86721E-30 | |
| 10 | Ifitm3 | 1.27 | 2.41 | 0.967 | 0.959 | 9.4278E-131 | 3.7429E-128 | |
| 10 | Fyb-ENSMUST00000090461.11-76514 | 1.26 | 2.39 | 0.531 | 0.549 | 1.13869E-15 | 4.52059E-13 | |
| 10 | Tlr2-ENSMUST00000029623.10-16593 | 1.20 | 2.30 | 0.737 | 0.756 | 1.13184E-38 | 4.49341E-36 | Signalling responsive |
| 10 | Itgb2-ENSMUST00000000299.13-59249 | 1.20 | 2.30 | 0.967 | 0.95 | 8.8225E-121 | 3.5025E-118 | |
| 10 | Havcr2-ENSMUST00000020668.14-62428 | 1.16 | 2.23 | 0.574 | 0.649 | 2.24339E-14 | 8.90626E-12 | |
| 10 | Tnfsf13-ENSMUST00000018896.13-63805 | 1.08 | 2.12 | 0.528 | 0.614 | 6.14391E-07 | 0.000243913 | |
| 10 | Lyz2-ENSMUST00000092163.7-60608 | 0.93 | 1.91 | 1 | 0.996 | 9.92801E-65 | 3.94142E-62 | |
| 10 | Lyn-ENSMUST00000103010.3-20309 | 0.84 | 1.78 | 0.969 | 0.962 | 3.62572E-77 | 1.43941E-74 | |
| 10 | Tgfb1-ENSMUST00000002678.9-40844 | 0.82 | 1.77 | 0.683 | 0.762 | 2.92852E-12 | 1.16262E-09 | |
| 10 | Xbp1-ENSMUST00000063084.15-61419 | 0.77 | 1.71 | 0.85 | 0.895 | 1.77458E-36 | 7.0451E-34 | |
| 10 | Casp1-ENSMUST00000027015.5-52899 | 0.75 | 1.68 | 0.526 | 0.675 | 0.001723998 | 0.684427048 | |
| 10 | Myd88-ENSMUST00000035092.6-57385 | 0.74 | 1.67 | 0.748 | 0.838 | 7.78267E-23 | 3.08972E-20 | |
| 11 | Cd79a-ENSMUST00000003469.7-40721 | 10.43 | 1376.16 | 0.743 | 0.004 | 0 | 0 | |
| 11 | Igkc-ENSMUST00000103410.2-35486 | 10.28 | 1240.20 | 0.892 | 0.009 | 0 | 0 | |
| 11 | Fcer2a-ENSMUST00000005678.5-48045 | 10.20 | 1174.37 | 0.389 | 0.001 | 0 | 0 | |
| 11 | Cd19-ENSMUST00000206325.1-46361 | 9.84 | 919.35 | 0.481 | 0.001 | 0 | 0 | |
| 11 | Iglc3-ENSMUST00000200235.1-79750 | 9.75 | 858.51 | 0.786 | 0.003 | 0 | 0 | |
| 11 | Mzb1-ENSMUST00000025211.4-85784 | 9.26 | 612.86 | 0.497 | 0.001 | 0 | 0 | |
| 11 | Pou2af1-ENSMUST00000034554.7-54351 | 8.83 | 456.04 | 0.368 | 0.002 | 0 | 0 | B cell interacting |
| 11 | Blk-ENSMUST00000014597.3-75650 | 8.40 | 338.11 | 0.272 | 0.001 | 0 | 0 | |
| 11 | Ighd-ENSMUST00000194162.5-70199 | 8.19 | 291.72 | 0.656 | 0.012 | 0 | 0 | |
| 11 | Cxcr5-ENSMUST00000179828.7-54074 | 5.87 | 58.48 | 0.259 | 0.007 | 0 | 0 | |
| 11 | H2-Ob-ENSMUST00000095342.9-83196 | 4.97 | 31.39 | 0.365 | 0.033 | 4.484E-280 | 1.7801E-277 | |
| 11 | Cd79b-ENSMUST00000167143.1-66704 | 4.81 | 27.97 | 0.63 | 0.094 | 6.2775E-306 | 2.4922E-303 | |
| 11 | Cd22-ENSMUST00000019248.12-41479 | 4.55 | 23.45 | 0.36 | 0.041 | 7.0604E-215 | 2.803E-212 | |
| 11 | Cd38-ENSMUST00000030964.5-27997 | 0.96 | 1.95 | 0.415 | 0.289 | 6.11644E-10 | 2.42823E-07 | |
